# Screening key genes and signaling pathways in COVID-19 infection and its associated complications by integrated bioinformatics analysis

**DOI:** 10.1101/2021.09.24.461631

**Authors:** Basavaraj Vastrad, Chanabasayya Vastrad

## Abstract

Severe acute respiratory syndrome corona virus 2 (SARS-CoV-2)/ coronavirus disease 2019 (COVID-19) infection is the leading cause of respiratory tract infection associated mortality worldwide. The aim of the current investigation was to identify the differentially expressed genes (DEGs) and enriched pathways in COVID-19 infection and its associated complications by bioinformatics analysis, and to provide potential targets for diagnosis and treatment. Valid next-generation sequencing (NGS) data of 93 COVID 19 samples and 100 non COVID 19 samples (GSE156063) were obtained from the Gene Expression Omnibus database. Gene ontology (GO) and REACTOME pathway enrichment analysis was conducted to identify the biological role of DEGs. In addition, a protein-protein interaction network, modules, miRNA-hub gene regulatory network, TF-hub gene regulatory network and receiver operating characteristic curve (ROC) analysis were used to identify the key genes. A total of 738 DEGs were identified, including 415 up regulated genes and 323 down regulated genes. Most of the DEGs were significantly enriched in immune system process, cell communication, immune system and signaling by NTRK1 (TRKA). Through PPI, modules, miRNA-hub gene regulatory network, TF-hub gene regulatory network analysis, ESR1, UBD, FYN, STAT1, ISG15, EGR1, ARRB2, UBE2D1, PRKDC and FOS were selected as hub genes, which were expressed in COVID-19 samples relative to those in non COVID-19 samples, respectively. Among them, ESR1, UBD, FYN, STAT1, ISG15, EGR1, ARRB2, UBE2D1, PRKDC and FOS were suggested to be diagonstic factors for COVID-19. The findings from this bioinformatics analysis study identified molecular mechanisms and the key hub genes that might contribute to COVID-19 infection and its associated complications.

## Introduction

Severe acute respiratory syndrome corona virus 2 (SARS-CoV-2)/ coronavirus disease 2019 (COVID-19) infection is one of the leading causes of respiratory tract infection [1]. COVID-19 is the fast spreading infectious disease worldwide, with a more than 200 million confirmed cases and 4.2 million confirmed death during August 2021 across nearly 200 countries according to the information given by the World Health Organization (WHO). The incidence rate of COVID-19 is high, which seriously affects the patient’s health complications, which includes diabetes mellitus [2], cardiovascular complications [3], neurological complications [4] and kidney complications [5]. However, the molecular mechanism underlying many COVID-19 and its associated complications causes remains unclear, resulting in a lack of effective treatment [6]. The higher prevalence and limited treatments of COVID-19 infection and its associated complications lead to considerable public health and economic difficulty [7]. Therefore, more reliable prognostic biomarkers should be explored as a target for improving the treatment effect and better understanding the underlying molecular mechanism of COVID-19 infection and its associated complications.

Studies have shown that genes includes ACE2 [8–12], TMPRSS2 [13–15], CD26 [16–18], HLA-DRB1 [19–23] and AKT1 [24–28] and signaling pathways includes cytokine signaling [29–31], innate immune signaling [32–36], JAK-STAT Signaling [37–41], IFN-AhR signaling [42] and STING pathway [43–44] are involved in the COVID-19 infection and its associated complications. Therefore, it is significant to further explore the molecular mechanisms of COVID-19 infection and its associated complications, and find molecular targets that can be used to diagnose early, prevent early, and treat early.

Next-generation sequencing (NGS) has thoroughly heightened the understanding of COVID-19 infection and its associated complications [45]. It provides a platform for public databases and exploring the molecular mechanisms of COVID-19 infection and its associated complications. A high-quality NGS could potentially link molecular biomarkers to the advancement, diagnosis, and treatment of COVID-19 infection and its associated complications.

In this investigation, we identified DEGs in expression profiling by high throughput sequencing data GSE156063 [46] from the Gene Expression Omnibus (GEO) database (http://www.ncbi.nlm.nih.gov/geo/) [47]. We performed Gene Ontology (GO) and REACTOME functional and pathway enrichment analyses, and constructed and analyzed a protein–protein interaction (PPI) network and isolated modules from PPI network, miRNA-hub gene regulatory network and TF-hub gene regulatory network were constructed and analyzed. Diagnostic values of hub genes were determined by receiver operating characteristic curve (ROC) analysis. Hub genes and the clinical prognosis of patients with COVID-19 infection and its associated complications were compared and analyzed with the objective of identifying novel prognostic biomarkers and therapeutic targets.

## Materials and methods

### Data resources

The expression profiling by high throughput sequencing GSE156063 [46] was downloaded from the GEO database and was used in this investigation. The dataset consisted of 93 COVID 19 samples and 100 non COVID 19 samples. The platform used was the GPL24676 Illumina NovaSeq 6000 (Homo sapiens).

### Identification of DEGs

The limma in R Bioconductor software package [48] was used to perform the identification of DEGs between COVID 19 samples and non COVID 19 samples. To precise the analysis of statistically significant genes and false-positives, we using the adjusted P-value and Benjamini and Hochberg false discovery rate method [49]. An adj P valuec<c0.05, and a |log FC (fold change) |c>c regulated genes and a |log FC (fold change) |c<-c were defined as the thresholds for DEG screening. The heatmaps of DEGs from each dataset were plotted by the gplots package in the R software. The DEGs are presented as volcano plots, generated using ggplot2 in the R software.

### GO and REACTOME pathway enrichment analysis of DEGs

The g:Profiler (http://biit.cs.ut.ee/gprofiler/) [50] is an online functional annotation tool to provide a comprehensive understanding of biological information of genes and proteins. GO term (http://www.geneontology.org) enrichment analysis is used broadly to classify the characteristic biological attributes of genes, gene products, and sequences, including biological process (BP), cell components (CC) and molecular function (MF) [51]. REACTOME (https://reactome.org/) [52] pathway enrichment analysis demonstrates the enriched signaling pathways in DEGs. P<0.05 was considered to represent statistical significance.

### Construction of the PPI network and module analysis

The IMEx interactome (https://www.imexconsortium.org/) is a public database harboring known and predicted protein-protein interactions [53]. Protein-protein interaction (PPI) is an indispensable approach for research on protein functions as it helps to clarify the interactions among proteins. In this study, the IMEx interactome database was used to construct a PPI network with an interaction score 0.4. The network was then visualized using the Cytoscape 3.8.2 (http://www.cytoscape.org/) [54]. Moreover, Network Analyzer [55], which is another plug-in of Cytoscape, was employed to study key nodes in the network with 4 methods includes node degree [56], betweenness centrality [57], stress centrality [58] and closeness centrality [59] are completed to explore the hub genes that were contained in the PPI network. Significant modules and hub genes were further analyzed and visualized with app PEWCC1 (http://apps.cytoscape.org/apps/PEWCC1) [60] plugged in Cytoscape. The two significant modules were displayed to show the density of nodes by IMEx interactome.

### MiRNA-hub gene regulatory network construction

MiRNAs controls gene expression under defined disease conditions through interaction with hub genes was analyzed. We applied miRNet database (https://www.mirnet.ca/) [61] to integrate 14 miRNA databases (TarBase, miRTarBase, miRecords, miRanda (S mansoni only), miR2Disease, HMDD, PhenomiR, SM2miR, PharmacomiR, EpimiR, starBase, TransmiR, ADmiRE, and TAM 2.0). We visualized miRNA-hub gene regulatory network by employing Cytoscape 3.8.2 [54].

### TF-hub gene regulatory network construction

TFs controls gene expression under defined disease conditions through interaction with hub genes was analyzed. We applied NetworkAnalyst database (https://www.networkanalyst.ca/) [62] to integrate TF database (JASPER). We visualized TF-hub gene regulatory network by employing Cytoscape 3.8.2 [54].

### Validation of hub genes by receiver operating characteristic curve (ROC) analysis

Receiver operating characteristic (ROC) analysis was used to assess the diagnostic value of hub genes in this research using pROC package in R statistical software [63]. Area under curve (AUC) analysis was implemented to determine the diagnostic value of hub genes. The AUC value > 0.8 showed a good diagnostic value for COVID 19.

## Results

### Identification of DEGs

A total of 738 DEGs (415 up regulated genes and 323 down regulated genes) were identified between COVID 19 samples and non COVID 19 samples according to methods described above. The heatmap of the DEGs has been shown in Fig. 1. The up and down regulated genes were listed in Table 1. The volcano plot showed (Fig. 2) that DEGs (up and down regulated genes) with the most significant logFC, and were used to further identify potential biomarkers.

**Fig. 1.**
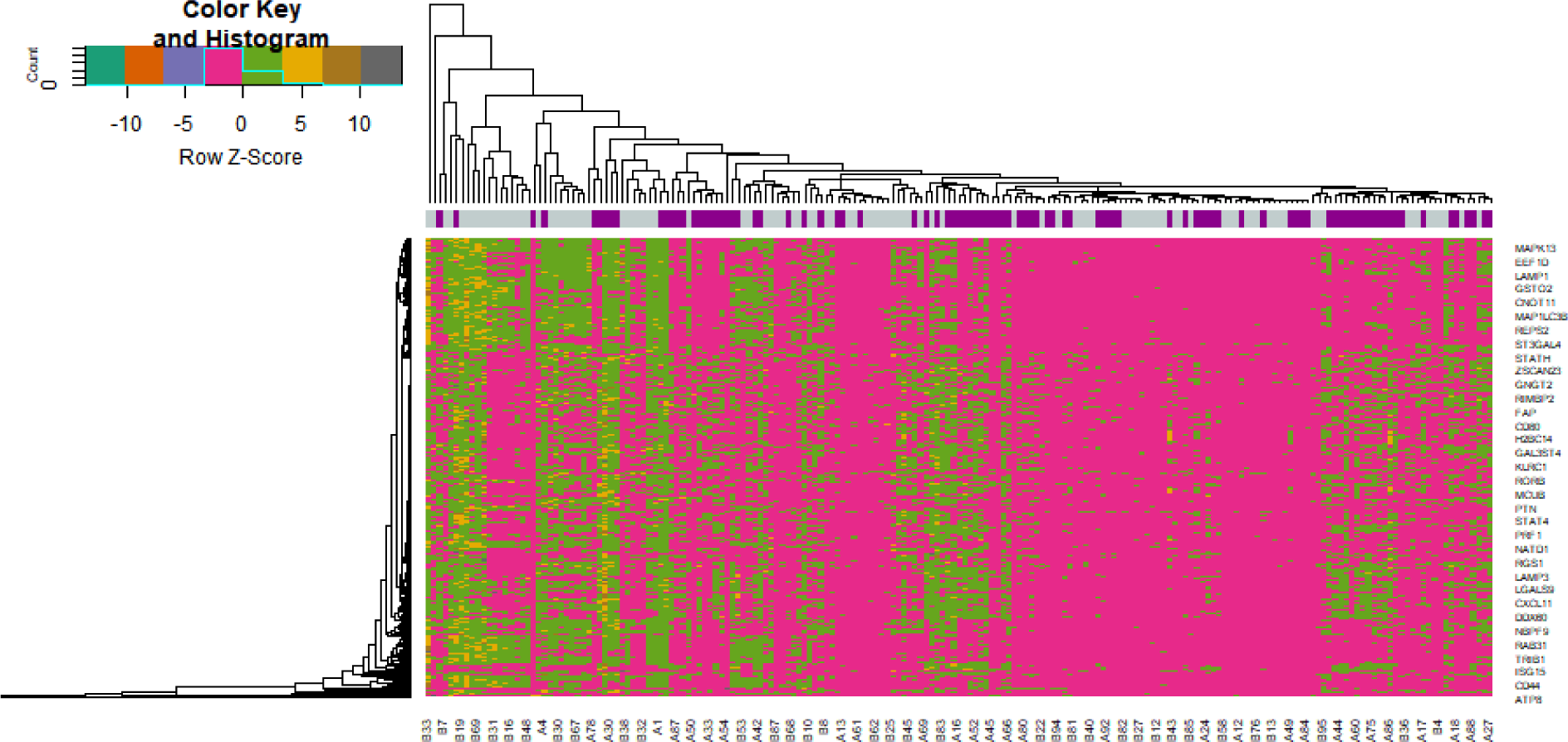
Heat map of differentially expressed genes. Legend on the top left indicate log fold change of genes. (A1 – A93= COVID 19 samples; B1 – B100= non COVID 19 samples)

**Fig. 2.**
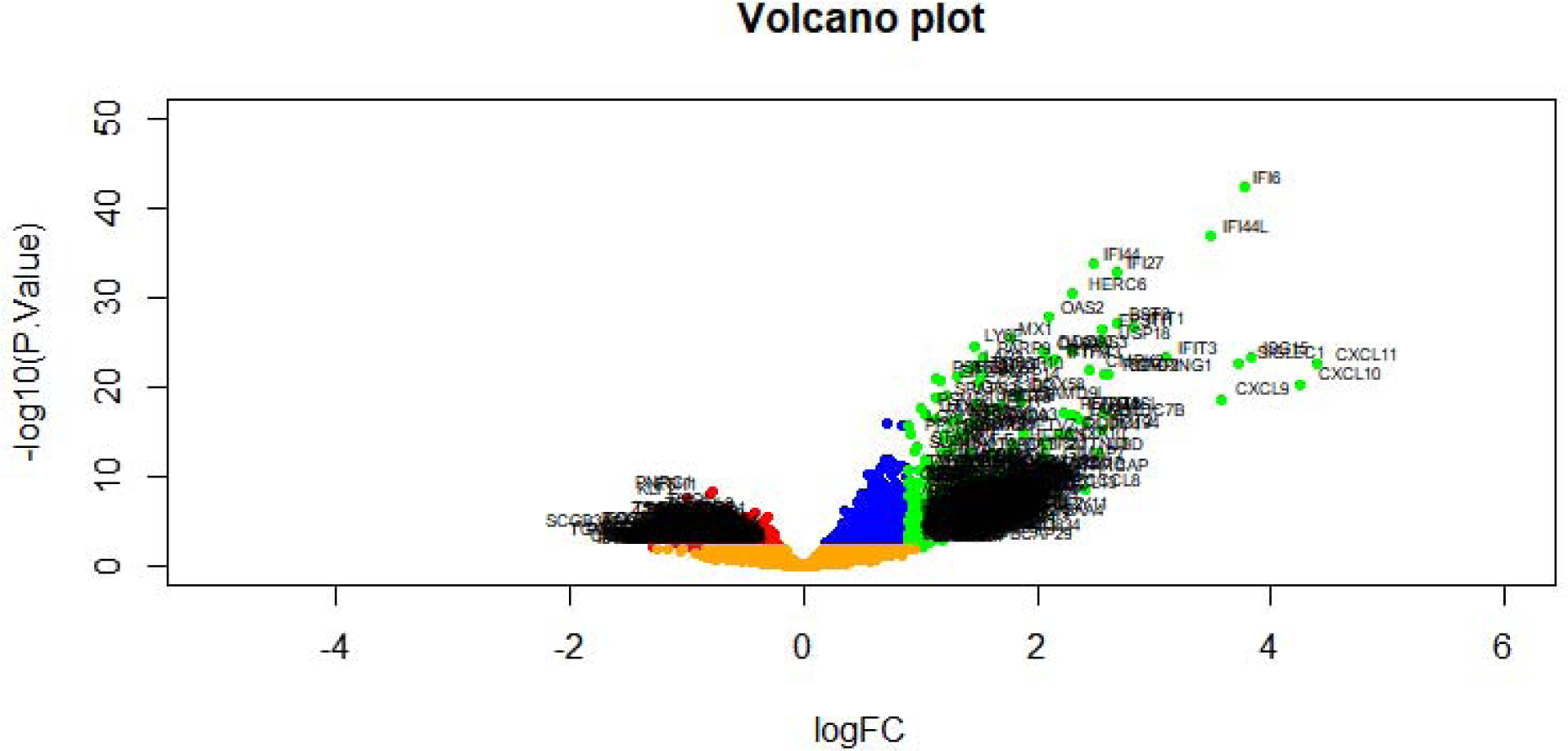
Volcano plot of differentially expressed genes. Genes with a significant change of more than two-fold were selected. Green dot represented up regulated significant genes and red dot represented down regulated significant genes.

### GO and REACTOME pathway enrichment analysis of DEGs

The consistently up and down regulated genes were clustered via the online website g:Profiler for the GO and REACTOME pathway enrichment analyses of DEGs in COVID 19. Three GO category results are presented in Table 2. GO terms cover BP, MF and CC categories. As to the BP, DEGs were significantly enriched in immune system process, response to stimulus, cell communication and localization. For the CC, DEGs were enriched in the cell periphery, plasma membrane, cytosol and intracellular anatomical structure. In terms of the MF, DEGs were enriched in binding, identical protein binding, protein binding and ion binding. Pathway enrichment results are presented in Table 3. The REACTOME pathway enrichment analysis revealed that DEGs were highly associated with the immune system, HDACs deacetylate histones, signaling by NTRK1 (TRKA) and innate immune system.

### Construction of the PPI network and module analysis

PPI network of DEGs was downloaded from IMEx interactome and further analyzed by Cytoscape. The PPI network included 5132 nodes and 12268 edges (Fig.3). The nodes such as ESR1, UBD, FYN, STAT1, ISG15, EGR1, ARRB2, UBE2D1, PRKDC and FOS with high node degree, betweenness centrality, stress centrality and closeness centrality were screened as hub proteins in the PPI network and listed in Table 4. In order to improve analysis of the PPI network, 2 significant modules were selected using the PEWCC1 plugin. Module 1 consisted of 16 nodes and 39 edges(Fig.4A), in which up regulated genes, including IFITM1, EIF2AK2, IFIT1, ISG15, IFIT2, GBP1, STAT2, IRF7 and STAT1. Module 2 consisted of 10 nodes and 27 edges (Fig.4B), in which down regulated genes, including GABARAPL1, GABARAP, RASSF5 and MAP1LC3B. Module 1 was mainly involved in immune system, infectious disease, immune system process, response to stimulus, binding, cell periphery and plasma membrane. Module 2 was mainly involved in cell communication, membrane trafficking, localization and cytosol.

**Fig. 3.**
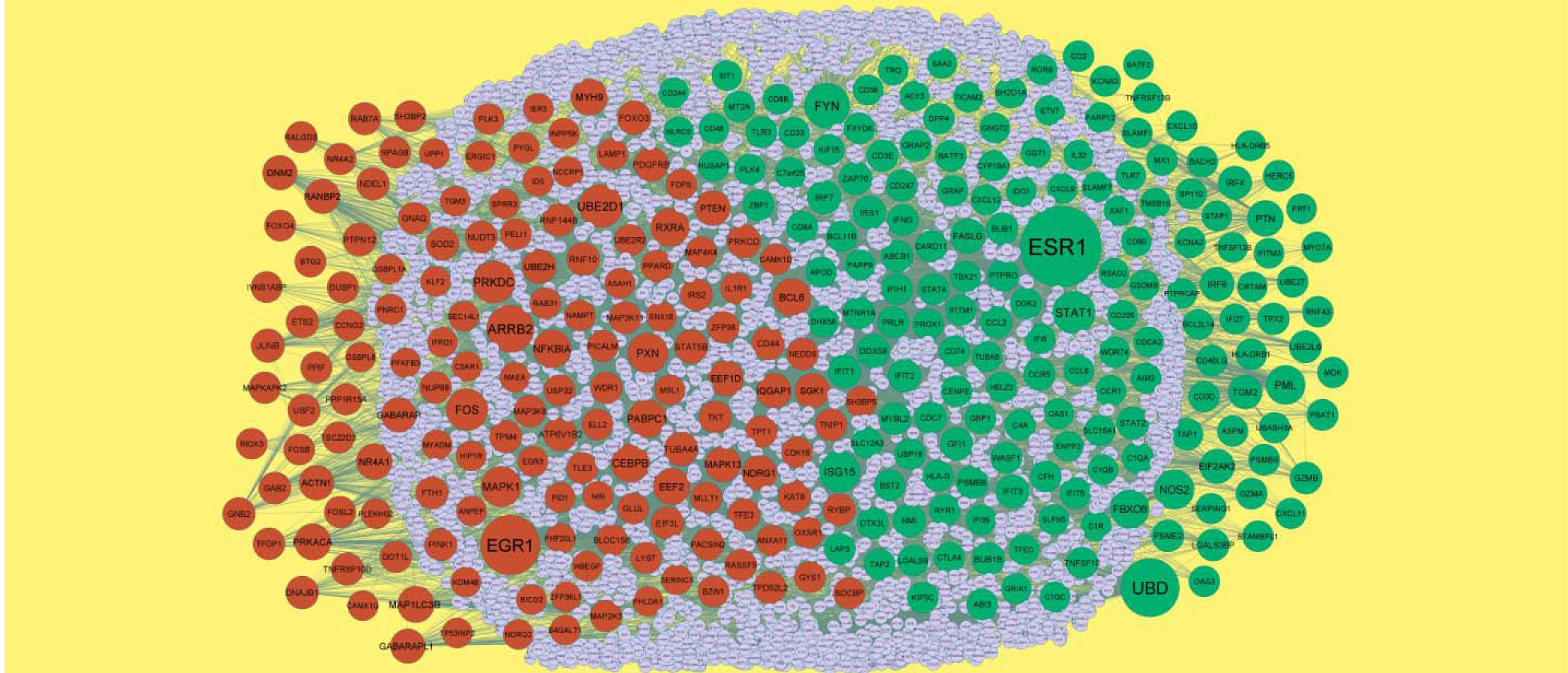
PPI network of DEGs. Up regulated genes are marked in green; down regulated genes are marked in red

**Fig. 4.**
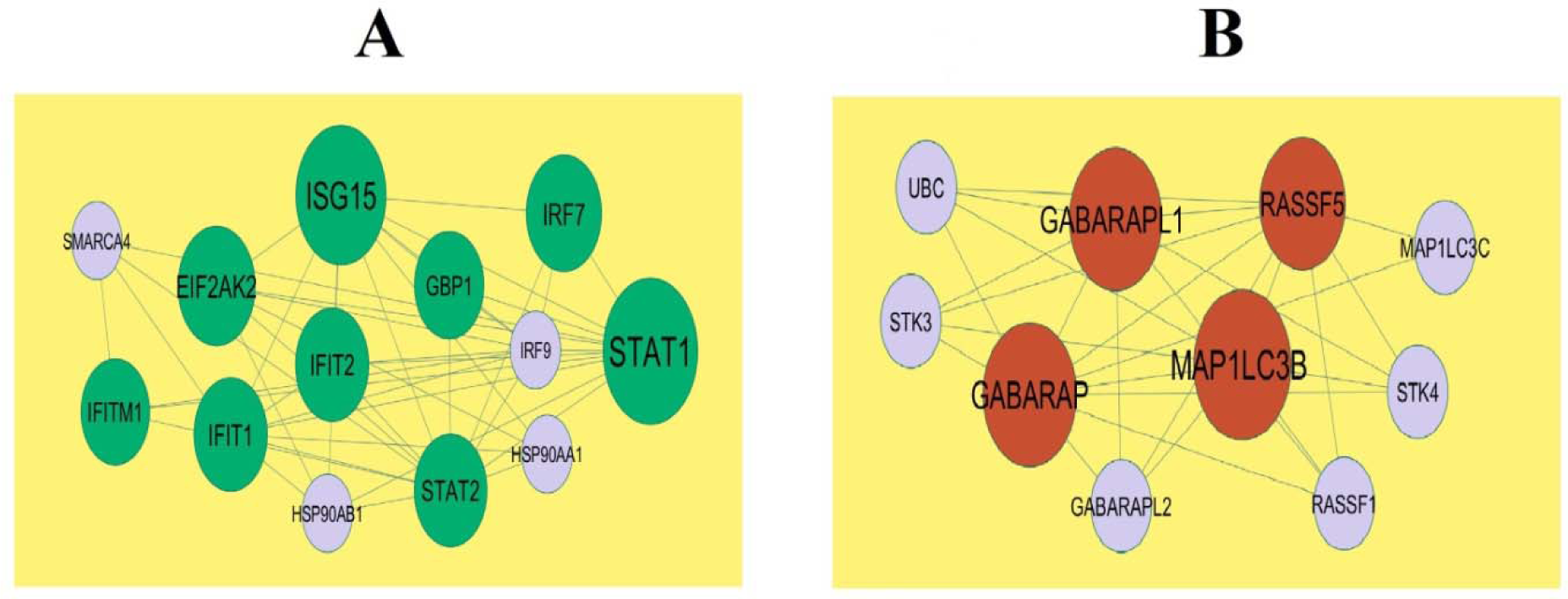
Modules of isolated form PPI of DEGs. (A) The most significant module was obtained from PPI networ with 16 nodes and 39 edges for up regulated genes (B) The most significant module was obtained from PPI network with 10 nodes and 27 edges for down regulated genes. Up regulated genes are marked in green; down regulated genes are marked in red

### MiRNA-hub gene regulatory network construction

The miRNA-hub gene regulatory network is shown in Fig. 5, which has 2626 nodes (hub genes:327; miRNAs: 2299) and 18613 interactions. The nodes with higher degrees are listed in Table 5, including IRF4 was the potential target of 140 miRNAs (ex; hsa-mir-4484); ESR1 was the potential target of 98 miRNAs (ex; hsa-mir-3668); EIF2AK2 was the potential target of 98 miRNAs (ex; hsa-mir- 190b); STAT1 was the potential target of 63 miRNAs (ex; hsa-mir-4693-5p); FYN was the potential target of 61 miRNAs (ex; hsa-mir-4326); MAPK1 was the potential target of 286 miRNAs (ex; hsa-mir-1470); EEF2 was the potential target of 229 miRNAs (ex; hsa-mir-2110); MYH9 was the potential target of 226 miRNAs (ex; hsa-mir-3615); PRKDC was the potential target of 164 miRNAs (ex; hsa-mir-4511); PABPC1 was the potential target of 151 miRNAs (ex; hsa-mir-1825).

**Fig. 5.**
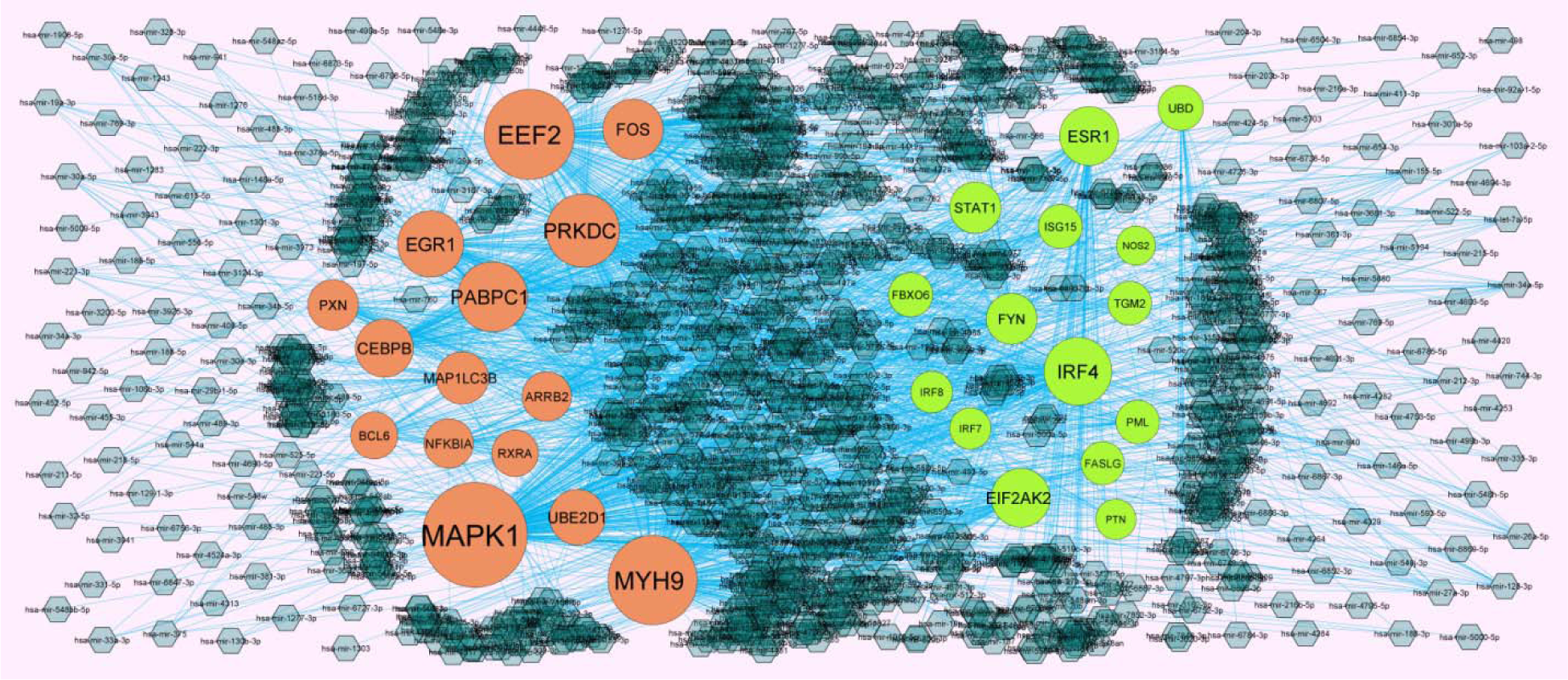
Target gene - miRNA regulatory network between target genes. The blue color diamond nodes represent the key miRNAs; up regulated genes are marked in green; down regulated genes are marked in red.

### TF-hub gene regulatory network construction

The TF-hub gene regulatory network is shown in Fig. 6, which has 59 nodes (hub genes:84; TFs:328) and 2719 interactions. The nodes with higher degrees are listed in Table 5, including ESR1 was the potential target of 22 TFs (ex; STAT3); IRF7 was the potential target of 14 TFs (ex; KLF5); IRF8 was the potential target of 12 TFs (ex; MAX); EIF2AK2 was the potential target of 10 TFs (ex; POU2F2); FYN was the potential target of 10 TFs (ex; ARID3A); BCL6 was the potential target of 17 TFs (ex; SREBF2); CEBPB was the potential target of 12 TFs (ex; E2F6); EGR1 was the potential target of 12 TFs (ex; SRF); RXRA was the potential target of 12 TFs (ex; GATA3); ARRB2 was the potential target of 12 TFs (ex; USF1).

**Fig. 6.**
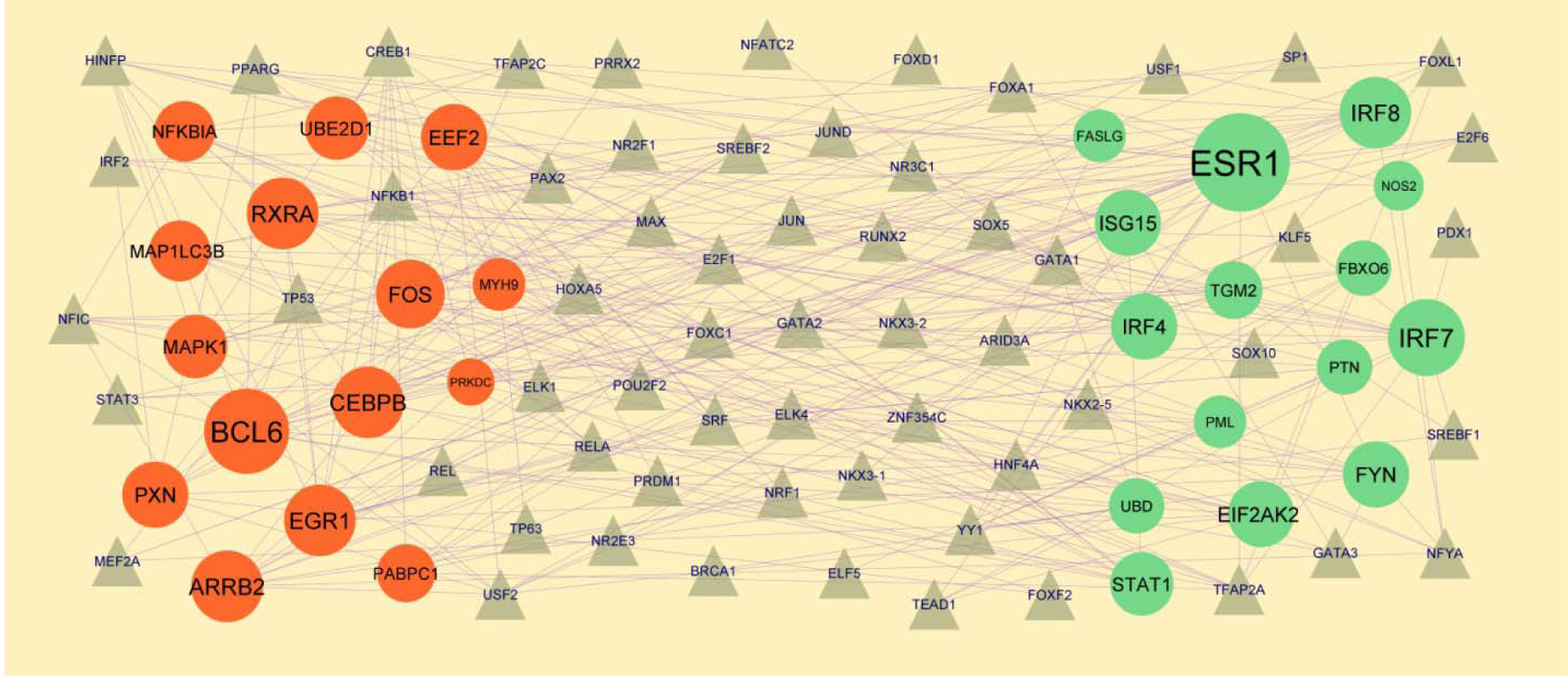
Target gene - TF regulatory network between target genes. The gray color triangle nodes represent the key TFs; up regulated genes are marked in green; down regulated genes are marked in red.

### Validation of hub genes by receiver operating characteristic curve (ROC) analysis

The ROC curves (Fig.7) of hub genes showed that their AUC as follows: ESR1 (0.913), UBD (0.928), FYN (0.938), STAT1 (0.931), ISG15 (0.946), EGR1 (0.913), ARRB2 (0.902), UBE2D1 (0.920), PRKDC (0.917) and FOS (0.924).

**Fig. 7.**
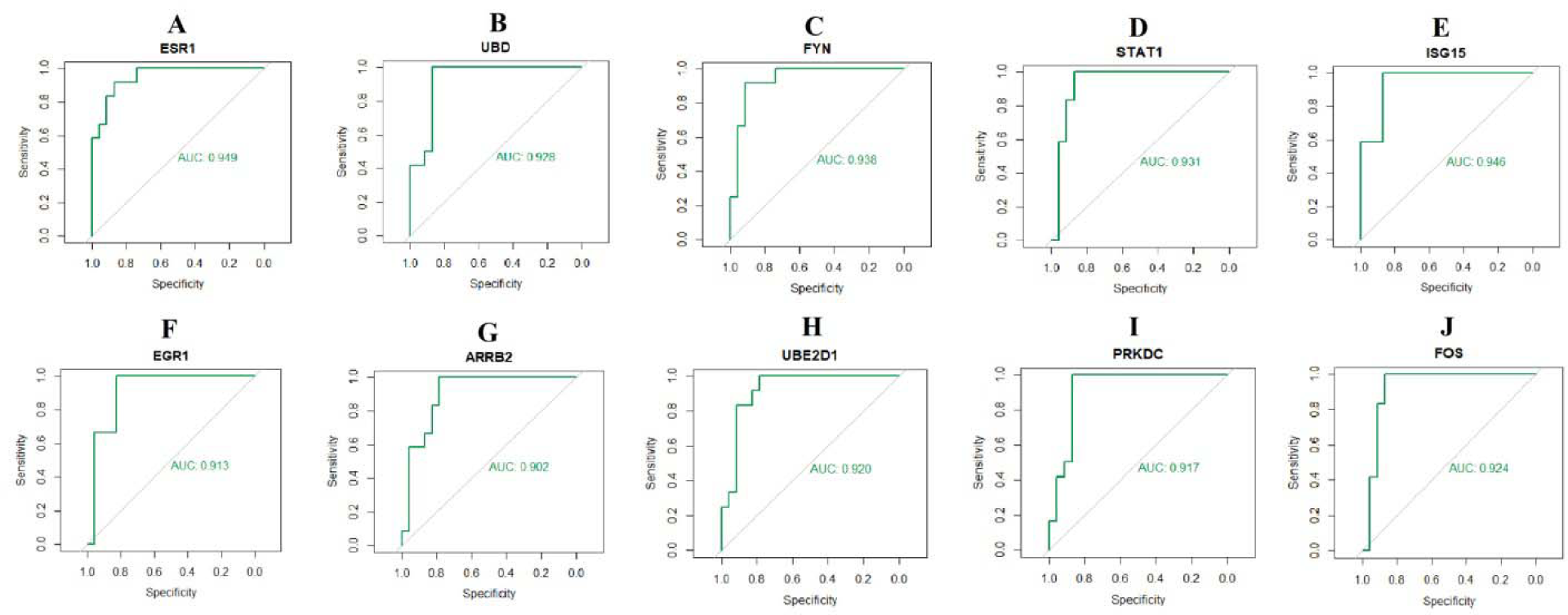
ROC curve analyses of hub genes. A) ESR1 B) UBD C) FYN D) STAT1 E) ISG15 F) EGR1 G) ARRB2 H) UBE2D1 I) PRKDC J) FOS

Since ROC curves had good specificity and sensitivity, ESR1, UBD, FYN, STAT1, ISG15, EGR1, ARRB2, UBE2D1, PRKDC and FOS had excellent diagnostic values for distinguishing COVID 19 samples and non COVID 19 samples.

## Discussion

Although many relevant studies of COVID-19 infection and its associated complications have been performed, early diagnosis, efficacy of treatment and prognosis for COVID-19 infection and its associated complications remain poorly determined. For prognosis, diagnosis and treatment, it is necessary to further understand the molecular mechanisms resulting in occurrence and advancement. Due to the advancement of NGS technology, the genetic modifications due to disease development can be detected, indicating gene targets for diagnosis, therapy and prognosis of infectious diseases. In the present investigation, a total of 738 DEGs between COVID-19 samples and non COVID-19 samples were identified, consisting of 415 up regulated and 323 down regulated genes. CXCL11 [64], CXCL10 [65], ISG15 [66], SIGLEC1 [67] and CXCL9 [64] are identified as a new type of COVID-19 susceptibility genes. CXCL10 might play a role in progression of diabetes mellitus [68]. CXCL10 [69], ISG15 [70], SIGLEC1 [71], CXCL9 [72], IFIT3 [73], KLF15 [74] and SPRR3 [75] have a potential role in the diagnosis and treatment of cardiovascular diseases. CXCL10 [76], SIGLEC1 [77] and IL1R2 [78] have been suggested to be associated with neurological diseases. CXCL9 [79], SIGLEC1 [80], IL1R2 [81], KLF15 [82] and CD207 [83] are important in the development of kidney diseases..

GO and REACTOME pathway enrichment pathway analyses were performed using g:Profiler. Immune system [84], HDACs deacetylate histones [85], adaptive immune system [86], innate immune system [87] and neutrophil degranulation [88] were linked with progression of COVID-19. Immune system [89], HDACs deacetylate histones [90], adaptive immune system [91], infectious disease [92], innate immune system [93] and neutrophil degranulation [94] were liable for progression of diabetes mellitus. Immune system [95], HDACs deacetylate histones [96], adaptive immune system [97], infectious disease [98], innate immune system [99] and neutrophil degranulation [100] were involved in progression of cardiovascular diseases. Immune system [101], HDACs deacetylate histones [102], adaptive immune system [103], innate immune system [104] and membrane trafficking [105] were linked with progression of neurological diseases. Immune system [106], infectious disease [107] and innate immune system [108] were associated with advancement of kidney diseases. IFI27 [109], UBD (ubiquitin D) [110], STAT1 [111], MT2A [112], CAMK4 [113], FASLG (Fas ligand) [114], ISG20 [115], ADGRF5 [116], HLA-G [117], MDK (midkine) [118], C4A [119], RGS1 [120], C1R [121], CTLA4 [122], CD38 [123], CX3CL1 [124], CD247 [125], SFRP1 [126], HLA DRB1 [23], AIM2 [127], TLR7 [128], TNFSF13B [129], CCR1 [130], TAP2 [131], APOL1 [132], CXCL12 [133], CFH (complement factor H) [134], RARRES2 [135], FYN (FYN proto-oncogene, Src family tyrosine kinase) [136], NLRC5 [137], CD74 [138], ACE2 [139], ABCB1 [140], POSTN (periostin) [141], TGM2 [142], COL4A3 [143], SLC12A3 [144] , IRS1 [145], APOL3 [146], STAT4 [147], CXCL8 [148], KLF2 [149], ADM (adrenomedullin) [150], EGR1 [151], TRIB1 [152], SOD2 [153], OXSR1 [154], IL6R [155], CD44 [156], LRG1 [157], PFKFB2 [158], PACSIN2 [159], MYH9 [160], DOT1L [161], CXCL16 [162], PTEN (phosphatase and tensin homolog) [163], FOXO3 [164], DDIT4 [165], PINK1 [166], IQGAP1 [167], ADIPOR1 [168], MTMR3 [169], USF2 [170], SGK1 [171], GSTO2 [172], ETS2 [173], TKT (transketolase) [174], CMIP (c-Maf inducing protein) [175], THBD (thrombomodulin) [176], ALPL (alkaline phosphatase, biomineralization associated) [177], GPX3 [178] and ELL2 [179] might serve as molecular markers for kidney diseases. OASL (2’-5’-oligoadenylate synthetase like) [180], IFIT2 [181], LAG3 [182], IFITM1 [183], IFITM3 [184], MX1 [185], IFI35 [186], TRIM22 [187], CCR1 [188], ZBP1 [189], FCGR3A [190], BPIFA1 [191], PSMB8 [192], CALHM6 [193], ETV7 [194], PARP12 [195], BCL6 [196] and DNAJB1 [197] were mainly involved in the various viral infections. CCL8 [198], GZMB (granzyme B) [199], CCL2 [200], IDO1 [201], CD80 [202], IFNG (interferon gamma) [203], STAT1 [204], CCR5 [205], SAA1 [206], MT2A [207], DPP4 [208], FASLG (Fas ligand) [209], CMKLR1 [210], ICOS (inducible T cell costimulator) [211], PDCD1 [212], SUCNR1 [213], IRF7 [214], HLA-F [215], HLA-G [216], MDK (midkine) [217], TIGIT (T cell immunoreceptor with Ig and ITIM domains) [218], CTLA4 [219], CD38 [220] TLR3 [221], CX3CL1 [222], SFRP1 [223], HLA-DRB1 [21], AIM2 [224], TLR7 [225], IRF8 [226], SLAMF7 [227], BCL11B [228], APOD (apolipoprotein D) [229], APOL1 [230], CXCL12 [231], CFH (complement factor H) [232], RARRES2 [233], RNASE6 [234], IL32 [235], NLRC5 [236], CD74 [237], ACE2 [238], FAP (fibroblast activation protein alpha) [239], ABCB1 [240], PCDH17 [241], POSTN (periostin) [242], KCNA2 [243], CD72 [244], TIMP4 [245], ESR1 [246], IRS1 [247], CMPK2 [248], IGFBP4 [249], CYP19A1 [250], CASQ2 [251], STAT4 [252], PPM1K [253], APOC1 [254], CXCL8 [255], KLF2 [256], ADM (adrenomedullin) [257], HBEGF (heparin binding EGF like growth factor) [258], KLF7 [259], DSC2 [260], DUSP1 [261], EGR1 [262], ADRB3 [263], ZFP36 [264], EGR3 [265], TRIB1 [266], LITAF (lipopolysaccharide induced TNF factor) [267], SOD2 [268], NR4A1 [269], BCL6 [270], IL6R [271], CD44 [272], MAP2K3 [273], NFKBIA (NFKB inhibitor alpha) [274], LRG1 [275], PFKFB2 [276], MAP4K4 [277], NR4A2 [278], IRS2 [279], TNIP1 [280], PTPN12 [281], DOT1L [282], CXCL16 [283], MAP3K11 [284], SORT1 [285], FOXO3 [286], SEMA4A [287], PINK1 [288], ADIPOR1 [289], SGK1 [290], MAPK1 [291], KLHL24 [292], ZFP36 [293], PFKFB3 [294], GYS1 [295], ETS2 [296], WDR1 [297], THBD (thrombomodulin) [298], ALPL (alkaline phosphatase, biomineralization associated) [299], FOSL2 [300] and GPX3 [301] contributes to the progression of cardiovascular diseases. CCL8 [302], LAG3 [303], CXCL13 [304], CCL2 [305], DDX58 [306], C1QA [307], C1QB [308], IFNG (interferon gamma) [309], STAT1 [310], CCR5 [311], HLA-DMB [312], CMKLR1 [313], LAMP3 [314], HLA-F [315], HLA-G [316], MDK (midkine) [317], IRF4 [318], C4A [319], EIF2AK2 [320], HLA-DRB5 [321], GSDMB (gasdermin B) [322], CD244 [323], C1R [324], CTLA4 [325], CD38 [326], CX3CL1 [327], HLA-DRB1 [328], AIM2 [329], TLR7 [330], IRF8 [331], TMEM176B [332], BCL11B [333], TAP2 [334], APOD (apolipoprotein D) [335], CXCL12 [336], CFH (complement factor H) [337], IL32 [338], NLRC5

[339], CD33 [340], ACE2 [341], KCNA3 [342], ABCB1 [343], GABRB2 [344]. MLC1 [345], KCNA2 [346], TIMP4 [347], EXOC3L4 [348], SAMD9L [349], TPX2 [350], MGAT3 [351], TMEM187 [352], ABI3 [353], TRANK1 [354], CYP19A1 [355], SH2D2A [356], WASF1 [357], FBXO6 [358], KIF5C [359], APOC1 [360], CXCL8 [361], KLF2 [362], ADM (adrenomedullin) [363], EGR1 [364], ADRB3 [365], EGR3 [366], ANPEP (alanylaminopeptidase, membrane) [367], SCN2A [368], KREMEN1 [369], NEDD9 [370], GPR85 [371], NDRG1 [372], SOD2 [373], ARRB2 [374], IL6R [375], GAB2 [376], NDEL1 [377], CEBPB (CCAAT enhancer binding protein beta) [378], FGD4 [379], INPP5A [380], SLC12A6 [381], PRKDC (protein kinase, DNA-activated, catalytic subunit) [382], MAP4K4 [383], PER1 [384], HIP1R [385], NR4A2 [386], INPP5K [387], IRS2 [388], ATP6V1B2 [389], PTEN (phosphatase and tensin homolog) [390], PELI1 [391], ZMIZ1 [392], NDRG2 [393], FOXO3 [394], DDIT4 [395], PINK1 [396], CPEB4 [397], ADIPOR1 [398], GLUL (glutamate-ammonia ligase) [399], PRKCD (protein kinase C delta) [400], USF2 [401], SGK1 [402], WDR45 [403], RAB7A [404], MAPK1 [405], FTH1 [406], AMPD2 [407], PICALM (phosphatidylinositol binding clathrin assembly protein) [408], UBAP1 [409], TUBA4A [410], TFE3 [411], EEF2 [412], UBE2H [413], CMIP (c-Maf inducing protein) [414], ANXA11 [415], THBD (thrombomodulin) [416], TREML2 [417], SLC25A37 [418], GPX3 [419] and KAT8 [420] have been shown to be activated in neurological diseases. LAG3 [421], IFITM3 [422], OAS1 [423], CXCL13 [424], CCL2 [425], IDO1 [426], CD80 [427], MX1 [428], IFNG (interferon gamma) [429], IFIH1 [430], PARP9 [431], STAT1 [432], KLRC2 [433], CCR5 [434], PARP14 [435], DPP4 [436], CAMK4 [437], HLA-G [438], MDK (midkine) [439], CD38 [440], TLR3 [441], CX3CL1 [442], HLA-DRB1 [19], TLR7 [443], IL7 [444], APOL1 [445], TNFRSF13B [446], STAT2 [447], IL32 [448], CH25H [449], CD33 [450], ACE2 [451], ABCB1 [452], CXCL8 [453], KLF2 [454], ADM (adrenomedullin) [455], C5AR1 [456], IL6R [457] and SDCBP (syndecan binding protein) [458] were found to be involved in the progression of COVID-19. GZMB (granzyme B) [459], OAS1 [460], CCL2 [461], IDO1 [462], CD80 [463], GZMA (granzyme A) [464], IFNG (interferon gamma) [465], IFIH1 [466], STAT1 [467], CCR5 [468], ADCY5 [469], MT2A [470], DPP4 [471], FASLG (Fas ligand) [472], HLA-DMB [473], CD3D [474], CMKLR1 [475], ICOS (inducible T cell costimulator) [476], PDCD1 [477], SUCNR1 [478], IRF7 [479], DEFB1 [480], HLA-G [481], ZAP70 [482], UBASH3A [483], HLA-DMA [484], TAP1 [485], CTLA4 [486], CD38 [487], TLR3 [488], CX3CL1 [489], CD247 [490], NOS2 [491], HLA-DRB1 [20], AIM2 [492], CFB (complement factor B) [493], TLR7 [494], SLAMF1 [495], BPIFA1 [496], CD300E [497], TAP2 [498], CD48 [499], APOL1 [500], BACH2 [501], CD226 [502], CXCL12 [503], STAT2 [504], RARRES2 [505], IL32 [506], ACE2 [507], CD72 [508], TGM2 [509], CD52 [510], COL4A3 [511], SLC12A3 [512], GPC4 [513], IRS1 [514], CYP19A1 [515], PROX1 [516], STAT4 [517], SEC16B [518], CXCL8 [519], KLF2 [520], DGAT2 [521], ADM (adrenomedullin) [522], IL1R1 [523], EGR1 [524], ADRB3 [525], SOD2 [526], ARRB2 [527], NR4A1 [528], IL6R [529], CD44 [530], LRG1 [531], FOXO4 [532], MAP4K4 [533], MYH9 [534], IRS2 [535], SULF2 [536], CXCL16 [537], FOXO3 [538], PINK1 [539], ADIPOR1 [540], GLUL (glutamate-ammonia ligase) [541], SGK1 [542], PFKFB3 [543], GYS1 [544], TP53INP2 [545], CMIP (c-Maf inducing protein) [546], STATH (statherin) [547], FOSL2 [548], GPX3 [549], CAMK1D [550] and RNASEK (ribonuclease K) [551] were identified to be closely associated with diabetes mellitus.

The hub genes from PPI network and modules defined in this study might lead to the progression of potential diagnostic, prognostic, or therapeutic biomarkers for COVID-19 infection and its associated complications. UBE2D1, FOS (Fos proto-oncogene, AP-1 transcription factor subunit), IFIT1, GBP1, GABARAPL1, GABARAP (GABA type A receptor-associated protein), RASSF5 and MAP1LC3B were might be the novel biomarkers for COVID-19 infection and its associated complications.

miRNA-hub gene regulatory network and TF-hub gene regulatory network can be regarded as key to the understanding COVID-19 infection and its associated complications and might also lead to new therapeutic approaches. hsa-mir-4484 [552], hsa-mir-4511 [553], STAT3 [554], KLF5 [555], SRF (Serum response factor) [556], GATA3 [557] and USF1 [558] have been implicated as a principal mediator of kidney diseases. Studies have reported that hsa-mir-190b [559], hsa-mir-4693-5p [560], hsa-mir-1470 [560], hsa-mir-1825 [561], STAT3 [562], SREBF2 [563] and SRF (Serum response factor) [564] are necessary for neurological diseases. hsa-mir-4693-5p [565] and STAT3 [432] expression have been identified in COVID-19 infection. hsa-mir-4326 [566], hsa-mir-1470 [567], hsa-mir-2110 [567], hsa-mir-3615 [568], hsa-mir-1825 [569], STAT3 [570], KLF5 [571], SREBF2 [572], E2F6 [573], GATA3 [574] and USF1 [575] have been shown to be activated in cardiovascular diseases. hsa-mir-3615 [576], hsa-mir-1825 [577], STAT3 [578], KLF5 [579], SREBF2 [580], GATA3 [Huda et al. 2018] and USF1 [Meex et al. 2008] were identified to be closely associated in patients with diabetes mellitus. PABPC1, RXRA (retinoid X receptor alpha), hsa-mir-3668, MAX (myc-associated factor X), POU2F2 and ARID3A were might be the novel biomarkers for COVID-19 infection and its associated complications.

In conclusion, the current investigation provided a comprehensive bioinformatics analysis of DEGs in COVID-19 infection. Analysis of these genes provided information regarding the molecular mechanisms of COVID-19 infection and its associated complications, and significant biomarkers or targets for the diagnosis and treatment of COVID-19 infection and its associated complications. However, further molecular biological experiments are required to confirm the function of the DEGs and pathways in COVID-19 infection and its associated complications.

## Acknowledgement

I thank Charles Langelier, University of California San Francisco, Medicine, San Francisco, CA, USA, very much, the author who deposited their profiling by high throughput sequencing dataset GSE156063, into the public GEO database.

## Ethical approval

This article does not contain any studies with human participants or animals performed by any of the authors.

## Informed consent

No informed consent because this study does not contain human or animals participants.

## Availability of data and materials

The datasets supporting the conclusions of this article are available in the GEO (Gene Expression Omnibus) (https://www.ncbi.nlm.nih.gov/geo/) repository. [(GSE156063) https://www.ncbi.nlm.nih.gov/geo/query/acc.cgi?acc=GSE156063)]

## Consent for publication

Not applicable.

## Author Contributions

B. V. - Writing original draft, and review and editing

C. V. - Software and investigation

## Authors

Basavaraj Vastrad ORCID ID: <u>0000-0003-2202-7637</u> Chanabasayya Vastrad ORCID ID: <u>0000-0003-3615-4450</u>

