## Supplementary material for "Screening key genes and signaling pathways in COVID-19 infection and its associated complications by integrated bioinformatics analysis": Tables

**Table 1** The statistical metrics for key differentially expressed genes (DEGs)

| **Gene Symbol** | **logFC** | **pValue** | **adj.P.Val** | **tvalue** | **Regulations** | **GeneName** |
| --- | --- | --- | --- | --- | --- | --- |
| CXCL11 | 4.401406 | 2.57E-23 | 1.87E-20 | 11.33776 | Up | C-X-C motif chemokine ligand 11 |
| CXCL10 | 4.252551 | 4.53E-21 | 2.19E-18 | 10.58757 | Up | C-X-C motif chemokine ligand 10 |
| ISG15 | 3.825839 | 5.15E-24 | 4.57E-21 | 11.56876 | Up | ISG15 ubiquitin like modifier |
| IFI6 | 3.773996 | 4.72E-43 | 7.55E-39 | 17.79855 | Up | interferon alpha inducible protein 6 |
| SIGLEC1 | 3.715637 | 1.95E-23 | 1.48E-20 | 11.37744 | Up | sialic acid binding Ig like lectin 1 |
| CXCL9 | 3.571786 | 2.15E-19 | 9.3E-17 | 10.01823 | Up | C-X-C motif chemokine ligand 9 |
| IFI44L | 3.477758 | 1.13E-37 | 9.04E-34 | 16.02092 | Up | interferon induced protein 44 like |
| IFIT3 | 3.104665 | 4.23E-24 | 4.1E-21 | 11.59678 | Up | interferon induced protein with tetratricopeptide repeats 3 |
| IFIT1 | 2.830657 | 2.02E-27 | 4.02E-24 | 12.68592 | Up | interferon induced protein with tetratricopeptide repeats 1 |
| BST2 | 2.680852 | 7.94E-28 | 1.81E-24 | 12.81766 | Up | bone marrow stromal cell antigen 2 |
| IFI27 | 2.674207 | 1.39E-33 | 5.57E-30 | 14.68738 | Up | interferon alpha inducible protein 27 |
| RSAD2 | 2.60122 | 4.23E-22 | 2.41E-19 | 10.93301 | Up | radical S-adenosyl methionine domain containing 2 |
| OASL | 2.578652 | 7.91E-18 | 2.94E-15 | 9.477793 | Up | 2'-5'-oligoadenylate synthetase like |
| SERPING1 | 2.57786 | 3.41E-22 | 2.02E-19 | 10.96417 | Up | serpin family G member 1 |
| IFIT2 | 2.564315 | 5.96E-16 | 1.62E-13 | 8.814323 | Up | interferon induced protein with tetratricopeptide repeats 2 |
| USP18 | 2.563927 | 1.19E-25 | 1.74E-22 | 12.10636 | Up | ubiquitin specific peptidase 18 |
| EPSTI1 | 2.552049 | 3.77E-27 | 6.69E-24 | 12.59731 | Up | epithelial stromal interaction 1 |
| UBD | 2.521131 | 2.37E-13 | 4.99E-11 | 7.858649 | Up | ubiquitin D |
| IFI44 | 2.483417 | 1.92E-34 | 1.02E-30 | 14.9671 | Up | interferon induced protein 44 |
| CMPK2 | 2.442471 | 1.33E-22 | 8.84E-20 | 11.10071 | Up | cytidine/uridine monophosphate kinase 2 |
| KLHDC7B | 2.416369 | 5.09E-17 | 1.59E-14 | 9.194299 | Up | kelch domain containing 7B |
| CCL8 | 2.408896 | 3.1E-09 | 3.02E-07 | 6.205329 | Up | C-C motif chemokine ligand 8 |
| LAG3 | 2.349605 | 3.81E-17 | 1.24E-14 | 9.238763 | Up | lymphocyte activating 3 |
| PLVAP | 2.330148 | 2.11E-17 | 7.19E-15 | 9.328442 | Up | plasmalemma vesicle associated protein |
| HERC6 | 2.303403 | 3.03E-31 | 9.68E-28 | 13.92863 | Up | HECT and RLD domain containing E3 ubiquitin protein ligase family member 6 |
| OAS3 | 2.3013 | 1.06E-24 | 1.3E-21 | 11.79514 | Up | 2'-5'-oligoadenylate synthetase 3 |
| IFITM1 | 2.298229 | 1.01E-17 | 3.58E-15 | 9.441038 | Up | interferon induced transmembrane protein 1 |
| RUFY4 | 2.260049 | 1.64E-15 | 4.09E-13 | 8.656511 | Up | RUN and FYVE domain containing 4 |
| PLAAT2 | 2.221665 | 8.55E-18 | 3.1E-15 | 9.466031 | Up | phospholipase A and acyltransferase 2 |
| GZMB | 2.205584 | 3.31E-11 | 4.86E-09 | 7.024641 | Up | granzyme B |
| CCDC194 | 2.18719 | 1.33E-15 | 3.49E-13 | 8.688581 | Up | coiled-coil domain containing 194 |
| RTP4 | 2.153042 | 6.9E-24 | 5.8E-21 | 11.52677 | Up | receptor transporter protein 4 |
| IFITM3 | 2.120491 | 1.48E-23 | 1.18E-20 | 11.41728 | Up | interferon induced transmembrane protein 3 |
| OAS2 | 2.091757 | 1.6E-28 | 4.27E-25 | 13.04386 | Up | 2'-5'-oligoadenylate synthetase 2 |
| GIMAP7 | 2.076202 | 1.96E-12 | 3.72E-10 | 7.508424 | Up | GTPase, IMAP family member 7 |
| PTPRCAP | 2.075752 | 7.66E-11 | 1.05E-08 | 6.877829 | Up | protein tyrosine phosphatase receptor type C associated protein |
| ANXA10 | 2.069933 | 1.91E-14 | 4.37E-12 | 8.266857 | Up | annexin A10 |
| SMTNL1 | 2.060448 | 2.93E-13 | 5.92E-11 | 7.82434 | Up | smoothelin like 1 |
| OAS1 | 2.055079 | 1.6E-24 | 1.7E-21 | 11.73649 | Up | 2'-5'-oligoadenylate synthetase 1 |
| DDX60 | 2.051508 | 1.23E-24 | 1.41E-21 | 11.77317 | Up | DExD/H-box helicase 60 |
| TFEC | 2.015633 | 3.07E-09 | 3.01E-07 | 6.207019 | Up | transcription factor EC |
| CXCL13 | 2.001792 | 5.85E-09 | 5.25E-07 | 6.085515 | Up | C-X-C motif chemokine ligand 13 |
| CCL2 | 1.922935 | 9.35E-07 | 4.76E-05 | 5.063183 | Up | C-C motif chemokine ligand 2 |
| PRF1 | 1.921821 | 7.22E-10 | 7.9E-08 | 6.475198 | Up | perforin 1 |
| SAA4 | 1.902739 | 4.4E-08 | 3.21E-06 | 5.693817 | Up | serum amyloid A4, constitutive |
| ETV7 | 1.880738 | 2.07E-15 | 4.93E-13 | 8.619913 | Up | ETS variant transcription factor 7 |
| OTOF | 1.86563 | 6.04E-10 | 6.8E-08 | 6.507751 | Up | otoferlin |
| SAMD9L | 1.854541 | 3.87E-19 | 1.63E-16 | 9.930956 | Up | sterile alpha motif domain containing 9 like |
| GIMAP8 | 1.84869 | 2.21E-11 | 3.43E-09 | 7.094874 | Up | GTPase, IMAP family member 8 |
| IDO1 | 1.841864 | 7.39E-11 | 1.02E-08 | 6.884227 | Up | indoleamine 2,3-dioxygenase 1 |
| CD80 | 1.839225 | 6.32E-09 | 5.64E-07 | 6.070935 | Up | CD80 molecule |
| GIMAP5 | 1.832542 | 3.26E-11 | 4.83E-09 | 7.027451 | Up | GTPase, IMAP family member 5 |
| DDX58 | 1.818209 | 6.74E-20 | 3.17E-17 | 10.19033 | Up | DExD/H-box helicase 58 |
| HERC5 | 1.789057 | 2.81E-14 | 6.32E-12 | 8.205216 | Up | HECT and RLD domain containing E3 ubiquitin protein ligase 5 |
| GZMA | 1.782817 | 2.1E-10 | 2.67E-08 | 6.698487 | Up | granzyme A |
| C1QA | 1.773784 | 2.35E-11 | 3.62E-09 | 7.084184 | Up | complement C1q A chain |
| BATF2 | 1.764917 | 2.23E-13 | 4.75E-11 | 7.86879 | Up | basic leucine zipper ATF-like transcription factor 2 |
| MX1 | 1.763338 | 3.86E-26 | 6.16E-23 | 12.26722 | Up | MX dynamin like GTPase 1 |
| GZMH | 1.754885 | 9.71E-10 | 1.02E-07 | 6.420881 | Up | granzyme H |
| C1QB | 1.740957 | 2.16E-10 | 2.72E-08 | 6.693226 | Up | complement C1q B chain |
| IFIT5 | 1.683532 | 1.72E-18 | 6.87E-16 | 9.707937 | Up | interferon induced protein with tetratricopeptide repeats 5 |
| GIMAP6 | 1.682011 | 9.61E-10 | 1.02E-07 | 6.422861 | Up | GTPase, IMAP family member 6 |
| ACE2 | 1.678445 | 1.54E-11 | 2.56E-09 | 7.157836 | Up | angiotensin I converting enzyme 2 |
| TRO | 1.670472 | 1.4E-10 | 1.84E-08 | 6.770644 | Up | trophinin |
| PRRX2 | 1.660273 | 2.83E-10 | 3.47E-08 | 6.645352 | Up | paired related homeobox 2 |
| PTPRO | 1.650953 | 1.55E-10 | 1.99E-08 | 6.753348 | Up | protein tyrosine phosphatase receptor type O |
| KCNA3 | 1.64492 | 1.01E-08 | 8.56E-07 | 5.981651 | Up | potassium voltage-gated channel subfamily A member 3 |
| NKG7 | 1.642654 | 3.97E-08 | 2.94E-06 | 5.714211 | Up | natural killer cell granule protein 7 |
| H2AC4 | 1.633797 | 2.05E-09 | 2.08E-07 | 6.28235 | Up | H2A clustered histone 4 |
| GIMAP4 | 1.631037 | 7.71E-09 | 6.81E-07 | 6.032911 | Up | GTPase, IMAP family member 4 |
| IL4I1 | 1.628141 | 5.9E-08 | 4.17E-06 | 5.635588 | Up | interleukin 4 induced 1 |
| H2AC16 | 1.625668 | 7.5E-10 | 8.15E-08 | 6.468328 | Up | H2A clustered histone 16 |
| IFNG | 1.611797 | 2.1E-07 | 1.31E-05 | 5.377281 | Up | interferon gamma |
| SLC38A5 | 1.610227 | 1.14E-08 | 9.48E-07 | 5.957179 | Up | solute carrier family 38 member 5 |
| APOL3 | 1.609793 | 2.62E-13 | 5.36E-11 | 7.842663 | Up | apolipoprotein L3 |
| PCSK5 | 1.609514 | 8.7E-08 | 5.94E-06 | 5.557486 | Up | proproteinconvertasesubtilisin/kexin type 5 |
| SAA2-SAA4 | 1.599454 | 6.09E-06 | 0.000236 | 4.647977 | Up | SAA2-SAA4 readthrough |
| H3C7 | 1.599106 | 6.05E-08 | 4.26E-06 | 5.630552 | Up | H3 clustered histone 7 |
| H2AC19 | 1.596973 | 1.05E-05 | 0.000369 | 4.5218 | Up | H2A clustered histone 19 |
| IFIH1 | 1.596331 | 4.7E-18 | 1.79E-15 | 9.556547 | Up | interferon induced with helicase C domain 1 |
| JCHAIN | 1.595812 | 5.33E-06 | 0.00021 | 4.678371 | Up | joining chain of multimeric IgA and IgM |
| TICAM2 | 1.581335 | 4.2E-07 | 2.4E-05 | 5.233376 | Up | toll like receptor adaptor molecule 2 |
| CD2 | 1.57353 | 3.05E-08 | 2.33E-06 | 5.766258 | Up | CD2 molecule |
| LRRN2 | 1.57115 | 5.41E-08 | 3.88E-06 | 5.652736 | Up | leucine rich repeat neuronal 2 |
| ZNF831 | 1.568621 | 4.26E-07 | 2.42E-05 | 5.230346 | Up | zinc finger protein 831 |
| H3C12 | 1.557481 | 3.2E-08 | 2.42E-06 | 5.756689 | Up | H3 clustered histone 12 |
| SLC43A1 | 1.555024 | 1.04E-09 | 1.09E-07 | 6.408465 | Up | solute carrier family 43 member 1 |
| MUC13 | 1.541793 | 4.56E-09 | 4.16E-07 | 6.132722 | Up | mucin 13, cell surface associated |
| IGLL5 | 1.541699 | 0.000463 | 0.006583 | 3.560105 | Up | immunoglobulin lambda like polypeptide 5 |
| ACOD1 | 1.540224 | 2.55E-05 | 0.000749 | 4.311696 | Up | aconitate decarboxylase 1 |
| ADAMDEC1 | 1.538375 | 2.82E-10 | 3.47E-08 | 6.645769 | Up | ADAM like decysin 1 |
| PARP9 | 1.533732 | 4.36E-24 | 4.1E-21 | 11.59255 | Up | poly(ADP-ribose) polymerase family member 9 |
| C1QC | 1.53228 | 1.27E-10 | 1.69E-08 | 6.788098 | Up | complement C1q C chain |
| MYBL2 | 1.52936 | 8.79E-08 | 5.97E-06 | 5.555484 | Up | MYB proto-oncogene like 2 |
| STAT1 | 1.528207 | 3.26E-22 | 2E-19 | 10.97078 | Up | signal transducer and activator of transcription 1 |
| KLRC2 | 1.526789 | 3.22E-07 | 1.91E-05 | 5.289043 | Up | killer cell lectin like receptor C2 |
| UBQLNL | 1.513469 | 1.81E-10 | 2.32E-08 | 6.724706 | Up | ubiquilin like |
| H2AC14 | 1.513406 | 3.11E-08 | 2.37E-06 | 5.762362 | Up | H2A clustered histone 14 |
| FAP | 1.507317 | 6.58E-06 | 0.00025 | 4.630106 | Up | fibroblast activation protein alpha |
| CCR5 | 1.506698 | 2.01E-09 | 2.05E-07 | 6.286076 | Up | C-C motif chemokine receptor 5 (gene/pseudogene) |
| PARP14 | 1.501339 | 2.82E-21 | 1.41E-18 | 10.65663 | Up | poly(ADP-ribose) polymerase family member 14 |
| NRSN1 | 1.499224 | 7.81E-08 | 5.4E-06 | 5.57921 | Up | neurensin 1 |
| PTN | 1.495135 | 3.02E-08 | 2.32E-06 | 5.76828 | Up | pleiotrophin |
| UBE2L6 | 1.487839 | 7.71E-19 | 3.16E-16 | 9.827978 | Up | ubiquitin conjugating enzyme E2 L6 |
| BTN3A3 | 1.477444 | 1.32E-16 | 3.85E-14 | 9.047461 | Up | butyrophilin subfamily 3 member A3 |
| SAA1 | 1.475907 | 1.03E-05 | 0.000365 | 4.525706 | Up | serum amyloid A1 |
| SAA2 | 1.466291 | 7.87E-06 | 0.000294 | 4.588937 | Up | serum amyloid A2 |
| H1-5 | 1.461102 | 1.02E-06 | 5.09E-05 | 5.045246 | Up | H1.5 linker histone, cluster member |
| C4B | 1.460596 | 3.53E-09 | 3.38E-07 | 6.180924 | Up | complement C4B (Chido blood group) |
| LGR6 | 1.457593 | 4.14E-10 | 4.9E-08 | 6.576465 | Up | leucine rich repeat containing G protein-coupled receptor 6 |
| LY6E | 1.456796 | 2.48E-25 | 3.3E-22 | 12.00248 | Up | lymphocyte antigen 6 family member E |
| LAP3 | 1.453984 | 4.06E-23 | 2.82E-20 | 11.27188 | Up | leucineaminopeptidase 3 |
| SAMD9 | 1.451563 | 2.29E-16 | 6.3E-14 | 8.963159 | Up | sterile alpha motif domain containing 9 |
| PDCD1LG2 | 1.448347 | 8.92E-09 | 7.63E-07 | 6.004903 | Up | programmed cell death 1 ligand 2 |
| H2BC11 | 1.446626 | 1.92E-09 | 1.97E-07 | 6.295053 | Up | H2B clustered histone 11 |
| BOLL | 1.443221 | 4.22E-09 | 3.92E-07 | 6.147325 | Up | boule homolog, RNA binding protein |
| IFI35 | 1.436444 | 8.37E-17 | 2.52E-14 | 9.118111 | Up | interferon induced protein 35 |
| ADCY5 | 1.430895 | 4.58E-07 | 2.59E-05 | 5.21523 | Up | adenylatecyclase 5 |
| HLA-DOB | 1.425989 | 8.32E-07 | 4.3E-05 | 5.08832 | Up | major histocompatibility complex, class II, DO beta |
| MT2A | 1.421462 | 2.47E-13 | 5.12E-11 | 7.852393 | Up | metallothionein 2A |
| WDR74 | 1.418729 | 5.7E-11 | 7.99E-09 | 6.92983 | Up | WD repeat domain 74 |
| FPR3 | 1.416451 | 1.04E-10 | 1.42E-08 | 6.822927 | Up | formyl peptide receptor 3 |
| FBXO39 | 1.414865 | 1.34E-07 | 8.81E-06 | 5.469165 | Up | F-box protein 39 |
| DPP4 | 1.413951 | 2.68E-06 | 0.000115 | 4.833276 | Up | dipeptidyl peptidase 4 |
| TNFSF10 | 1.413089 | 2.73E-22 | 1.75E-19 | 10.99633 | Up | TNF superfamily member 10 |
| H3C2 | 1.407793 | 2.95E-06 | 0.000125 | 4.811757 | Up | H3 clustered histone 2 |
| HAPLN3 | 1.406707 | 1.04E-08 | 8.8E-07 | 5.974626 | Up | hyaluronan and proteoglycan link protein 3 |
| TBX21 | 1.406435 | 9.38E-07 | 4.76E-05 | 5.062474 | Up | T-box transcription factor 21 |
| H2BC14 | 1.394612 | 9.16E-07 | 4.68E-05 | 5.067688 | Up | H2B clustered histone 14 |
| H2BC17 | 1.39358 | 2.26E-07 | 1.4E-05 | 5.362739 | Up | H2B clustered histone 17 |
| H2AC12 | 1.387431 | 2.66E-08 | 2.07E-06 | 5.793394 | Up | H2A clustered histone 12 |
| PPAN-P2RY11 | 1.384448 | 1.07E-06 | 5.33E-05 | 5.034009 | Up | PPAN-P2RY11 readthrough |
| FAM83A | 1.384419 | 6.27E-12 | 1.14E-09 | 7.311617 | Up | family with sequence similarity 83 member A |
| PSMB9 | 1.383189 | 1.31E-15 | 3.49E-13 | 8.69114 | Up | proteasome 20S subunit beta 9 |
| CAMK4 | 1.383067 | 6.21E-07 | 3.36E-05 | 5.150833 | Up | calcium/calmodulin dependent protein kinase IV |
| FASLG | 1.379234 | 2.35E-08 | 1.85E-06 | 5.817488 | Up | Fas ligand |
| DHX58 | 1.376598 | 1.96E-15 | 4.74E-13 | 8.628561 | Up | DExH-box helicase 58 |
| HLA-DMB | 1.376028 | 3.7E-09 | 3.48E-07 | 6.171933 | Up | major histocompatibility complex, class II, DM beta |
| ABCB1 | 1.375395 | 1.51E-06 | 7.05E-05 | 4.959 | Up | ATP binding cassette subfamily B member 1 |
| H4C13 | 1.366251 | 2.57E-07 | 1.57E-05 | 5.335674 | Up | H4 clustered histone 13 |
| MS4A4A | 1.366197 | 7.9E-09 | 6.94E-07 | 6.02822 | Up | membrane spanning 4-domains A4A |
| RNF43 | 1.358987 | 0.000178 | 0.003349 | 3.81965 | Up | ring finger protein 43 |
| BTN3A1 | 1.357653 | 6.58E-12 | 1.18E-09 | 7.303358 | Up | butyrophilin subfamily 3 member A1 |
| CD3D | 1.35711 | 4.15E-07 | 2.38E-05 | 5.235751 | Up | CD3d molecule |
| CD3E | 1.355388 | 3.51E-07 | 2.05E-05 | 5.271208 | Up | CD3e molecule |
| IGSF9B | 1.34921 | 1.06E-05 | 0.00037 | 4.520117 | Up | immunoglobulin superfamily member 9B |
| P2RX5 | 1.34915 | 2.45E-05 | 0.00073 | 4.321142 | Up | purinergic receptor P2X 5 |
| CMKLR1 | 1.343192 | 1.84E-06 | 8.25E-05 | 4.916279 | Up | chemerin chemokine-like receptor 1 |
| ICOS | 1.341532 | 5.98E-07 | 3.25E-05 | 5.158662 | Up | inducible T cell costimulator |
| CALHM6 | 1.340715 | 3.04E-06 | 0.000128 | 4.805062 | Up | calcium homeostasis modulator family member 6 |
| H2BC3 | 1.340252 | 1.47E-07 | 9.48E-06 | 5.451527 | Up | H2B clustered histone 3 |
| GBP1 | 1.339844 | 3.54E-10 | 4.22E-08 | 6.604802 | Up | guanylate binding protein 1 |
| H4C14 | 1.336797 | 4.17E-12 | 7.75E-10 | 7.380672 | Up | H4 clustered histone 14 |
| GRAP2 | 1.333151 | 3.44E-06 | 0.000141 | 4.777415 | Up | GRB2 related adaptor protein 2 |
| XAF1 | 1.321697 | 4.12E-17 | 1.32E-14 | 9.226759 | Up | XIAP associated factor 1 |
| PRLR | 1.317945 | 4.41E-06 | 0.000177 | 4.721282 | Up | prolactin receptor |
| TRIM22 | 1.310769 | 6.88E-22 | 3.79E-19 | 10.86216 | Up | tripartite motif containing 22 |
| TPX2 | 1.308316 | 4.31E-09 | 3.98E-07 | 6.143296 | Up | TPX2 microtubule nucleation factor |
| LAMP3 | 1.307967 | 1.74E-15 | 4.27E-13 | 8.647092 | Up | lysosomal associated membrane protein 3 |
| IQSEC3 | 1.304031 | 1.73E-05 | 0.000545 | 4.404463 | Up | IQ motif and Sec7 domain ArfGEF 3 |
| H3C3 | 1.303964 | 5.36E-07 | 2.96E-05 | 5.182123 | Up | H3 clustered histone 3 |
| NMI | 1.296479 | 8.7E-15 | 2.02E-12 | 8.392554 | Up | N-myc and STAT interactor |
| EOMES | 1.292873 | 1.67E-06 | 7.65E-05 | 4.937302 | Up | eomesodermin |
| COL5A1 | 1.285448 | 2.94E-05 | 0.000831 | 4.276862 | Up | collagen type V alpha 1 chain |
| THEMIS | 1.275577 | 2.3E-06 | 0.0001 | 4.866994 | Up | thymocyte selection associated |
| PDCD1 | 1.274703 | 2.83E-05 | 0.000806 | 4.286 | Up | programmed cell death 1 |
| ISG20 | 1.273865 | 1.07E-09 | 1.11E-07 | 6.402836 | Up | interferon stimulated exonuclease gene 20 |
| NT5C3A | 1.26918 | 5.78E-17 | 1.77E-14 | 9.174944 | Up | 5'-nucleotidase, cytosolic IIIA |
| SUCNR1 | 1.267652 | 1.63E-06 | 7.5E-05 | 4.942182 | Up | succinate receptor 1 |
| PSAT1 | 1.266509 | 2.18E-08 | 1.74E-06 | 5.832345 | Up | phosphoserine aminotransferase 1 |
| PLK4 | 1.263388 | 2.72E-07 | 1.64E-05 | 5.324066 | Up | polo like kinase 4 |
| IRF7 | 1.261576 | 4.31E-08 | 3.16E-06 | 5.698151 | Up | interferon regulatory factor 7 |
| HLA-F | 1.260994 | 6.67E-14 | 1.46E-11 | 8.065758 | Up | major histocompatibility complex, class I, F |
| MGAT3 | 1.257607 | 7.14E-08 | 4.98E-06 | 5.597203 | Up | beta-1,4-mannosyl-glycoprotein 4-beta-N-acetylglucosaminyltransferase |
| GABRB2 | 1.245678 | 4.01E-06 | 0.000162 | 4.742989 | Up | gamma-aminobutyric acid type A receptor beta2 subunit |
| PPM1J | 1.241277 | 1.68E-07 | 1.07E-05 | 5.424044 | Up | protein phosphatase, Mg2+/Mn2+ dependent 1J |
| HESX1 | 1.235182 | 4E-08 | 2.95E-06 | 5.712576 | Up | HESX homeobox 1 |
| LGALS3BP | 1.228479 | 1.17E-19 | 5.34E-17 | 10.10869 | Up | galectin 3 binding protein |
| GPR171 | 1.228274 | 1.15E-05 | 0.000398 | 4.500169 | Up | G protein-coupled receptor 171 |
| ADGRF5 | 1.218011 | 2.69E-06 | 0.000115 | 4.832296 | Up | adhesion G protein-coupled receptor F5 |
| PARP12 | 1.215013 | 1.4E-15 | 3.61E-13 | 8.681068 | Up | poly(ADP-ribose) polymerase family member 12 |
| ATP10A | 1.213572 | 3.81E-09 | 3.56E-07 | 6.166388 | Up | ATPase phospholipid transporting 10A (putative) |
| DEFB1 | 1.211219 | 7.3E-07 | 3.86E-05 | 5.11622 | Up | defensin beta 1 |
| SHFL | 1.210974 | 4.35E-15 | 1.02E-12 | 8.502576 | Up | shiftless antiviral inhibitor of ribosomal frameshifting |
| CD8A | 1.20866 | 5.74E-07 | 3.14E-05 | 5.167479 | Up | CD8a molecule |
| H3C8 | 1.207391 | 1.01E-06 | 5.05E-05 | 5.047585 | Up | H3 clustered histone 8 |
| GRAP | 1.206435 | 0.000182 | 0.003405 | 3.813859 | Up | GRB2 related adaptor protein |
| KRT39 | 1.205387 | 3.05E-07 | 1.83E-05 | 5.300073 | Up | keratin 39 |
| HLA-G | 1.201932 | 2.92E-08 | 2.26E-06 | 5.774564 | Up | major histocompatibility complex, class I, G |
| HSH2D | 1.200504 | 1.84E-11 | 2.93E-09 | 7.127183 | Up | hematopoietic SH2 domain containing |
| CD8B | 1.198553 | 1.4E-05 | 0.000463 | 4.454297 | Up | CD8b molecule |
| C17orf67 | 1.196268 | 1.67E-07 | 1.06E-05 | 5.425354 | Up | chromosome 17 open reading frame 67 |
| FUOM | 1.195372 | 1.11E-06 | 5.46E-05 | 5.026106 | Up | fucosemutarotase |
| MDK | 1.1953 | 3.03E-13 | 6.06E-11 | 7.818336 | Up | midkine |
| GGT1 | 1.194131 | 0.001344 | 0.013767 | 3.252355 | Up | gamma-glutamyltransferase 1 |
| H3C10 | 1.181443 | 5.05E-07 | 2.8E-05 | 5.194589 | Up | H3 clustered histone 10 |
| CACNA2D1 | 1.181064 | 0.000174 | 0.003292 | 3.826508 | Up | calcium voltage-gated channel auxiliary subunit alpha2delta 1 |
| RBM11 | 1.178008 | 3.22E-08 | 2.43E-06 | 5.755448 | Up | RNA binding motif protein 11 |
| IRF4 | 1.176965 | 3.94E-05 | 0.00105 | 4.204992 | Up | interferon regulatory factor 4 |
| SIRPG | 1.175205 | 2.49E-05 | 0.000737 | 4.316981 | Up | signal regulatory protein gamma |
| ASCL3 | 1.174818 | 9.33E-06 | 0.000335 | 4.549529 | Up | achaete-scute family bHLH transcription factor 3 |
| C4A | 1.173587 | 1.27E-06 | 6.13E-05 | 4.997223 | Up | complement C4A (Rodgers blood group) |
| EIF2AK2 | 1.169837 | 1.64E-21 | 8.44E-19 | 10.73595 | Up | eukaryotic translation initiation factor 2 alpha kinase 2 |
| SH2D1A | 1.168984 | 5.76E-05 | 0.001391 | 4.11067 | Up | SH2 domain containing 1A |
| ZAP70 | 1.168554 | 1.4E-05 | 0.000463 | 4.455069 | Up | zeta chain of T cell receptor associated protein kinase 70 |
| TIGIT | 1.163349 | 1.57E-05 | 0.000501 | 4.428093 | Up | T cell immunoreceptor with Ig and ITIM domains |
| RGS1 | 1.163058 | 1.59E-05 | 0.000506 | 4.424014 | Up | regulator of G protein signaling 1 |
| ZNF683 | 1.160545 | 7.74E-05 | 0.001797 | 4.036373 | Up | zinc finger protein 683 |
| KIF15 | 1.157673 | 3.77E-06 | 0.000154 | 4.756825 | Up | kinesin family member 15 |
| PCDH17 | 1.153155 | 3.96E-06 | 0.000161 | 4.74579 | Up | protocadherin 17 |
| HLA-DRB5 | 1.152815 | 1.24E-06 | 6.03E-05 | 5.001415 | Up | major histocompatibility complex, class II, DR beta 5 |
| GSDMB | 1.149462 | 1.79E-11 | 2.9E-09 | 7.131056 | Up | gasdermin B |
| UBASH3A | 1.14733 | 8.6E-06 | 0.000315 | 4.568519 | Up | ubiquitin associated and SH3 domain containing A |
| H2AC17 | 1.146451 | 4.91E-07 | 2.75E-05 | 5.200351 | Up | H2A clustered histone 17 |
| LOC114483834 | 1.145693 | 0.000267 | 0.004444 | 3.711776 | Up | ABCF2-H2B readthrough |
| CD244 | 1.143792 | 1.27E-05 | 0.00043 | 4.477359 | Up | CD244 molecule |
| RORB | 1.142761 | 1.04E-07 | 6.98E-06 | 5.520529 | Up | RAR related orphan receptor B |
| IGFBP4 | 1.14086 | 2.63E-07 | 1.6E-05 | 5.331239 | Up | insulin like growth factor binding protein 4 |
| MUSK | 1.140619 | 0.000245 | 0.004204 | 3.73506 | Up | muscle associated receptor tyrosine kinase |
| C1R | 1.139833 | 1.38E-06 | 6.52E-05 | 4.9784 | Up | complement C1r |
| STAP1 | 1.139635 | 1.79E-07 | 1.13E-05 | 5.411085 | Up | signal transducing adaptor family member 1 |
| KLRC1 | 1.13937 | 1.44E-06 | 6.77E-05 | 4.969682 | Up | killer cell lectin like receptor C1 |
| CD69 | 1.133738 | 0.000119 | 0.002469 | 3.924855 | Up | CD69 molecule |
| PSME2 | 1.133014 | 9.12E-22 | 4.86E-19 | 10.82122 | Up | proteasome activator subunit 2 |
| PML | 1.129201 | 3.61E-17 | 1.2E-14 | 9.246657 | Up | promyelocyticleukemia |
| SLAMF8 | 1.126474 | 0.000327 | 0.005142 | 3.656317 | Up | SLAM family member 8 |
| AGR3 | 1.126324 | 8.54E-07 | 4.39E-05 | 5.082692 | Up | anterior gradient 3, protein disulphide isomerase family member |
| LKAAEAR1 | 1.126315 | 3.51E-05 | 0.000959 | 4.233504 | Up | LKAAEAR motif containing 1 |
| SPATS2L | 1.12537 | 1.62E-19 | 7.19E-17 | 10.06046 | Up | spermatogenesis associated serine rich 2 like |
| RIMBP2 | 1.123332 | 0.000218 | 0.00388 | 3.765968 | Up | RIMS binding protein 2 |
| WARS1 | 1.119747 | 1.65E-11 | 2.72E-09 | 7.145302 | Up | tryptophanyl-tRNAsynthetase 1 |
| CENPE | 1.115371 | 4.91E-05 | 0.001244 | 4.150575 | Up | centromere protein E |
| HLA-DMA | 1.114999 | 4.91E-12 | 9.02E-10 | 7.353076 | Up | major histocompatibility complex, class II, DM alpha |
| SLC16A1 | 1.113249 | 2.69E-07 | 1.62E-05 | 5.32649 | Up | solute carrier family 16 member 1 |
| TMEM255A | 1.111071 | 6.43E-06 | 0.000246 | 4.635496 | Up | transmembrane protein 255A |
| TAP1 | 1.111069 | 2.41E-10 | 3.01E-08 | 6.674058 | Up | transporter 1, ATP binding cassette subfamily B member |
| H2AC13 | 1.106711 | 2.65E-05 | 0.000771 | 4.302394 | Up | H2A clustered histone 13 |
| CTLA4 | 1.105201 | 0.000159 | 0.00306 | 3.850854 | Up | cytotoxic T-lymphocyte associated protein 4 |
| CD38 | 1.102368 | 1.45E-08 | 1.17E-06 | 5.912062 | Up | CD38 molecule |
| TLR3 | 1.097491 | 4.84E-11 | 6.84E-09 | 6.958566 | Up | toll like receptor 3 |
| POSTN | 1.092424 | 0.001064 | 0.011754 | 3.32165 | Up | periostin |
| CX3CL1 | 1.092129 | 4.93E-07 | 2.75E-05 | 5.19956 | Up | C-X3-C motif chemokine ligand 1 |
| PRH1-PRR4 | 1.091218 | 3.71E-05 | 0.001007 | 4.219694 | Up | PRH1-PRR4 readthrough |
| VAMP5 | 1.086383 | 1.91E-08 | 1.53E-06 | 5.857824 | Up | vesicle associated membrane protein 5 |
| HDX | 1.085964 | 1.32E-05 | 0.000442 | 4.468047 | Up | highly divergent homeobox |
| PLLP | 1.084269 | 6.06E-06 | 0.000236 | 4.649178 | Up | plasmolipin |
| CD247 | 1.083968 | 0.000305 | 0.004877 | 3.675427 | Up | CD247 molecule |
| GBP5 | 1.082263 | 0.000139 | 0.002799 | 3.88492 | Up | guanylate binding protein 5 |
| KLRC4 | 1.076409 | 8.57E-05 | 0.001949 | 4.010218 | Up | killer cell lectin like receptor C4 |
| MYO7A | 1.075836 | 0.000133 | 0.002688 | 3.896456 | Up | myosin VIIA |
| ENDOD1 | 1.075522 | 5.84E-08 | 4.15E-06 | 5.637511 | Up | endonuclease domain containing 1 |
| NOS2 | 1.073796 | 0.000104 | 0.002246 | 3.961719 | Up | nitric oxide synthase 2 |
| MUC19 | 1.071408 | 0.001711 | 0.016404 | 3.179378 | Up | mucin 19, oligomeric |
| TMEM187 | 1.070957 | 8.82E-06 | 0.000322 | 4.562525 | Up | transmembrane protein 187 |
| H2BC7 | 1.06985 | 2.16E-06 | 9.49E-05 | 4.880699 | Up | H2B clustered histone 7 |
| ASPM | 1.064392 | 2.43E-05 | 0.000727 | 4.322537 | Up | abnormal spindle microtubule assembly |
| DUOXA2 | 1.06383 | 8.26E-06 | 0.000304 | 4.57778 | Up | dual oxidase maturation factor 2 |
| SFRP1 | 1.060562 | 7.31E-05 | 0.00171 | 4.050696 | Up | secreted frizzled related protein 1 |
| CDCA7 | 1.059395 | 4.49E-05 | 0.001155 | 4.172964 | Up | cell division cycle associated 7 |
| HLA-DRB1 | 1.057518 | 2.37E-09 | 2.39E-07 | 6.255374 | Up | major histocompatibility complex, class II, DR beta 1 |
| DGKI | 1.055618 | 0.000122 | 0.002509 | 3.919005 | Up | diacylglycerol kinase iota |
| ABI3 | 1.055241 | 5.07E-05 | 0.001269 | 4.142891 | Up | ABI family member 3 |
| SP140 | 1.054482 | 0.00017 | 0.003231 | 3.832679 | Up | SP140 nuclear body protein |
| TRANK1 | 1.053793 | 1.37E-08 | 1.12E-06 | 5.921916 | Up | tetratricopeptide repeat and ankyrin repeat containing 1 |
| AIM2 | 1.052474 | 0.00047 | 0.006662 | 3.55603 | Up | absent in melanoma 2 |
| CFB | 1.051696 | 1.41E-12 | 2.79E-10 | 7.562965 | Up | complement factor B |
| TLR7 | 1.050929 | 5.06E-06 | 0.0002 | 4.690096 | Up | toll like receptor 7 |
| MARCHF1 | 1.046483 | 1.44E-06 | 6.77E-05 | 4.969182 | Up | membrane associated ring-CH-type finger 1 |
| TNFSF13B | 1.043139 | 9.2E-05 | 0.002047 | 3.992169 | Up | TNF superfamily member 13b |
| SLAMF1 | 1.043091 | 0.000778 | 0.009461 | 3.412962 | Up | signaling lymphocytic activation molecule family member 1 |
| DTX3L | 1.042672 | 1.16E-17 | 4.01E-15 | 9.420384 | Up | deltex E3 ubiquitin ligase 3L |
| BUB1 | 1.042601 | 3.7E-06 | 0.000151 | 4.760713 | Up | BUB1 mitotic checkpoint serine/threonine kinase |
| IRF8 | 1.039334 | 0.000142 | 0.00284 | 3.878924 | Up | interferon regulatory factor 8 |
| PLAAT4 | 1.039212 | 7.38E-12 | 1.3E-09 | 7.283749 | Up | phospholipase A and acyltransferase 4 |
| H2AC11 | 1.037053 | 1.37E-05 | 0.000456 | 4.459439 | Up | H2A clustered histone 11 |
| SLAMF7 | 1.0367 | 5.5E-05 | 0.001341 | 4.122384 | Up | SLAM family member 7 |
| CCR1 | 1.035127 | 0.003333 | 0.02598 | 2.971015 | Up | C-C motif chemokine receptor 1 |
| CYP19A1 | 1.03203 | 0.000447 | 0.006447 | 3.570182 | Up | cytochrome P450 family 19 subfamily A member 1 |
| GFI1 | 1.031841 | 2.27E-05 | 0.000686 | 4.339154 | Up | growth factor independent 1 transcriptional repressor |
| NUPR1 | 1.03178 | 3.09E-10 | 3.76E-08 | 6.629512 | Up | nuclear protein 1, transcriptional regulator |
| UBE2T | 1.031599 | 8.28E-07 | 4.29E-05 | 5.089484 | Up | ubiquitin conjugating enzyme E2 T |
| TPSAB1 | 1.029666 | 0.005271 | 0.035016 | 2.82105 | Up | tryptase alpha/beta 1 |
| NUSAP1 | 1.027552 | 8.51E-05 | 0.001938 | 4.012021 | Up | nucleolar and spindle associated protein 1 |
| IL7 | 1.026475 | 1.1E-06 | 5.42E-05 | 5.028224 | Up | interleukin 7 |
| PERM1 | 1.026132 | 0.000126 | 0.002574 | 3.910663 | Up | PPARGC1 and ESRR induced regulator, muscle 1 |
| ZBP1 | 1.025941 | 1.81E-05 | 0.000565 | 4.393986 | Up | Z-DNA binding protein 1 |
| DOK2 | 1.024526 | 0.000311 | 0.004946 | 3.669943 | Up | docking protein 2 |
| SH2D2A | 1.022026 | 0.000112 | 0.002369 | 3.94161 | Up | SH2 domain containing 2A |
| FCGR3A | 1.021586 | 0.00031 | 0.004935 | 3.670848 | Up | Fc fragment of IgG receptor IIIa |
| BPIFA1 | 1.017903 | 0.004576 | 0.031791 | 2.867945 | Up | BPI fold containing family A member 1 |
| B4GALT6 | 1.017433 | 9.87E-07 | 4.99E-05 | 5.051571 | Up | beta-1,4-galactosyltransferase 6 |
| RHEX | 1.016556 | 1.41E-05 | 0.000465 | 4.452569 | Up | regulator of hemoglobinization and erythroid cell expansion |
| TMEM176B | 1.015269 | 8.81E-05 | 0.001989 | 4.003184 | Up | transmembrane protein 176B |
| FAM241B | 1.014896 | 2.51E-07 | 1.54E-05 | 5.340666 | Up | family with sequence similarity 241 member B |
| CD300E | 1.01489 | 0.002163 | 0.019405 | 3.10745 | Up | CD300e molecule |
| MLC1 | 1.012654 | 0.002677 | 0.022462 | 3.040789 | Up | modulator of VRAC current 1 |
| PSMB8 | 1.010733 | 2.32E-18 | 9.06E-16 | 9.662568 | Up | proteasome 20S subunit beta 8 |
| H4C11 | 1.009858 | 1.03E-07 | 6.91E-06 | 5.524312 | Up | H4 clustered histone 11 |
| BCL11B | 1.009779 | 3.71E-05 | 0.001007 | 4.219823 | Up | BAF chromatin remodeling complex subunit BCL11B |
| TAP2 | 1.009615 | 4.6E-08 | 3.34E-06 | 5.685134 | Up | transporter 2, ATP binding cassette subfamily B member |
| ZNF311 | 1.008804 | 4E-06 | 0.000162 | 4.743075 | Up | zinc finger protein 311 |
| TEDC2 | 1.00586 | 4.21E-05 | 0.0011 | 4.188573 | Up | tubulin epsilon and delta complex 2 |
| BATF3 | 1.005358 | 0.000141 | 0.002828 | 3.88105 | Up | basic leucine zipper ATF-like transcription factor 3 |
| CD48 | 1.001842 | 0.000649 | 0.008317 | 3.465044 | Up | CD48 molecule |
| UBQLN3 | 1.000697 | 3.92E-05 | 0.001048 | 4.206225 | Up | ubiquilin 3 |
| CYP2C18 | 0.999334 | 1.85E-05 | 0.000572 | 4.388811 | Up | cytochrome P450 family 2 subfamily C member 18 |
| SYT13 | 0.996966 | 3.78E-05 | 0.00102 | 4.215291 | Up | synaptotagmin 13 |
| APOD | 0.995956 | 4.16E-05 | 0.001089 | 4.191974 | Up | apolipoprotein D |
| BCL2L14 | 0.993597 | 3.63E-08 | 2.71E-06 | 5.731854 | Up | BCL2 like 14 |
| MX2 | 0.992711 | 1.16E-06 | 5.71E-05 | 5.015852 | Up | MX dynamin like GTPase 2 |
| GIMAP1 | 0.991907 | 1.82E-05 | 0.000566 | 4.392546 | Up | GTPase, IMAP family member 1 |
| PYHIN1 | 0.99186 | 7.96E-06 | 0.000297 | 4.586198 | Up | pyrin and HIN domain family member 1 |
| PROX1 | 0.990926 | 0.00025 | 0.004258 | 3.729761 | Up | prosperohomeobox 1 |
| MB21D2 | 0.990392 | 2.73E-05 | 0.000786 | 4.295117 | Up | Mab-21 domain containing 2 |
| CCDC103 | 0.990158 | 0.000268 | 0.00445 | 3.711104 | Up | coiled-coil domain containing 103 |
| SERPINI1 | 0.988812 | 1.32E-05 | 0.000441 | 4.468814 | Up | serpin family I member 1 |
| ZNF367 | 0.988292 | 1.06E-05 | 0.00037 | 4.520874 | Up | zinc finger protein 367 |
| APOL1 | 0.986536 | 1.92E-11 | 3.01E-09 | 7.119218 | Up | apolipoprotein L1 |
| H2BC10 | 0.984986 | 2.82E-05 | 0.000804 | 4.28716 | Up | H2B clustered histone 10 |
| CLCNKB | 0.984494 | 0.000549 | 0.007433 | 3.512335 | Up | chloride voltage-gated channel Kb |
| KCNA2 | 0.980876 | 5.09E-05 | 0.001269 | 4.141694 | Up | potassium voltage-gated channel subfamily A member 2 |
| GBP4 | 0.976728 | 4.36E-10 | 5.09E-08 | 6.566822 | Up | guanylate binding protein 4 |
| H4C6 | 0.976582 | 6.79E-05 | 0.001613 | 4.069274 | Up | H4 clustered histone 6 |
| CD40LG | 0.975434 | 5.67E-05 | 0.001378 | 4.114519 | Up | CD40 ligand |
| BACH2 | 0.975149 | 0.000753 | 0.009288 | 3.422335 | Up | BTB domain and CNC homolog 2 |
| SP110 | 0.974814 | 1.15E-08 | 9.48E-07 | 5.956979 | Up | SP110 nuclear body protein |
| CD226 | 0.972552 | 1.9E-06 | 8.48E-05 | 4.909144 | Up | CD226 molecule |
| CXCL12 | 0.971945 | 0.000816 | 0.009774 | 3.39927 | Up | C-X-C motif chemokine ligand 12 |
| TNFRSF13B | 0.971733 | 0.000201 | 0.003651 | 3.788497 | Up | TNF receptor superfamily member 13B |
| DPP10 | 0.971373 | 1.55E-05 | 0.0005 | 4.4298 | Up | dipeptidyl peptidase like 10 |
| STAT2 | 0.970844 | 4.31E-14 | 9.58E-12 | 8.136094 | Up | signal transducer and activator of transcription 2 |
| ANKRD20A2 | 0.969825 | 0.000131 | 0.002661 | 3.900363 | Up | ankyrin repeat domain 20 family member A2 |
| CD72 | 0.968515 | 1.87E-05 | 0.000577 | 4.386396 | Up | CD72 molecule |
| CFH | 0.96751 | 6.39E-10 | 7.14E-08 | 6.497561 | Up | complement factor H |
| MCUB | 0.967317 | 8.06E-07 | 4.2E-05 | 5.09509 | Up | mitochondrial calcium uniporter dominant negative beta subunit |
| TGM2 | 0.967147 | 6.73E-10 | 7.47E-08 | 6.487992 | Up | transglutaminase 2 |
| WASF1 | 0.965169 | 2.71E-05 | 0.000785 | 4.296437 | Up | WASP family member 1 |
| H2BC13 | 0.964089 | 0.000358 | 0.005474 | 3.631719 | Up | H2B clustered histone 13 |
| CRTAM | 0.963133 | 2.74E-05 | 0.000786 | 4.293927 | Up | cytotoxic and regulatory T cell molecule |
| CASQ2 | 0.961354 | 0.004203 | 0.03005 | 2.895852 | Up | calsequestrin 2 |
| RARRES2 | 0.957694 | 5.9E-05 | 0.001415 | 4.104914 | Up | retinoic acid receptor responder 2 |
| GNLY | 0.957639 | 0.000431 | 0.006248 | 3.58068 | Up | granulysin |
| DUS4L-BCAP29 | 0.957304 | 0.002552 | 0.021804 | 3.055816 | Up | DUS4L-BCAP29 readthrough |
| RNASE6 | 0.955791 | 3.25E-05 | 0.000899 | 4.252596 | Up | ribonuclease A family member k6 |
| CDC7 | 0.954561 | 1.3E-05 | 0.000437 | 4.47147 | Up | cell division cycle 7 |
| CD52 | 0.953347 | 0.000691 | 0.008705 | 3.447229 | Up | CD52 molecule |
| GNGT2 | 0.952694 | 0.000341 | 0.005283 | 3.644913 | Up | G protein subunit gamma transducin 2 |
| GRIK1 | 0.95212 | 0.000148 | 0.002922 | 3.868572 | Up | glutamate ionotropic receptor kainate type subunit 1 |
| CARD11 | 0.951886 | 2.12E-06 | 9.33E-05 | 4.884984 | Up | caspase recruitment domain family member 11 |
| COL4A3 | 0.951406 | 0.0028 | 0.023088 | 3.02652 | Up | collagen type IV alpha 3 chain |
| H3C13 | 0.949323 | 4.38E-05 | 0.001135 | 4.17917 | Up | H3 clustered histone 13 |
| C7orf25 | 0.948401 | 0.000263 | 0.004394 | 3.715987 | Up | chromosome 7 open reading frame 25 |
| IL15RA | 0.948202 | 1.98E-06 | 8.82E-05 | 4.89982 | Up | interleukin 15 receptor subunit alpha |
| FXYD6 | 0.947726 | 2.27E-05 | 0.000686 | 4.339392 | Up | FXYD domain containing ion transport regulator 6 |
| CPLX2 | 0.947697 | 0.001468 | 0.014604 | 3.225895 | Up | complexin 2 |
| C1orf54 | 0.944952 | 2.42E-05 | 0.000723 | 4.324384 | Up | chromosome 1 open reading frame 54 |
| SLFN5 | 0.944853 | 1.2E-13 | 2.59E-11 | 7.97021 | Up | schlafen family member 5 |
| FYN | 0.944334 | 8.2E-06 | 0.000302 | 4.579549 | Up | FYN proto-oncogene, Src family tyrosine kinase |
| RASGRF2 | 0.941455 | 4.05E-05 | 0.001074 | 4.198131 | Up | Ras protein specific guanine nucleotide releasing factor 2 |
| IL32 | 0.939257 | 5.89E-05 | 0.001414 | 4.105383 | Up | interleukin 32 |
| HASPIN | 0.93806 | 0.000507 | 0.007088 | 3.534909 | Up | histone H3 associated protein kinase |
| HELZ2 | 0.938057 | 1.59E-07 | 1.02E-05 | 5.434373 | Up | helicase with zinc finger 2 |
| RYR1 | 0.936541 | 0.003676 | 0.027722 | 2.939435 | Up | ryanodine receptor 1 |
| MS4A7 | 0.936077 | 6.43E-07 | 3.46E-05 | 5.143377 | Up | membrane spanning 4-domains A7 |
| TUBA8 | 0.935841 | 5.7E-05 | 0.001382 | 4.113383 | Up | tubulin alpha 8 |
| PLEK2 | 0.935841 | 1.94E-07 | 1.22E-05 | 5.393574 | Up | pleckstrin 2 |
| BUB1B | 0.935804 | 0.000405 | 0.005974 | 3.597412 | Up | BUB1 mitotic checkpoint serine/threonine kinase B |
| PAK1IP1 | 0.935039 | 1.73E-06 | 7.83E-05 | 4.930142 | Up | PAK1 interacting protein 1 |
| ADGRB2 | 0.933954 | 3.88E-05 | 0.001042 | 4.208588 | Up | adhesion G protein-coupled receptor B2 |
| H2BC6 | 0.931813 | 2.85E-06 | 0.000122 | 4.819691 | Up | H2B clustered histone 6 |
| ZSCAN23 | 0.929206 | 0.001098 | 0.012004 | 3.312596 | Up | zinc finger and SCAN domain containing 23 |
| PBDC1 | 0.928296 | 7.26E-09 | 6.44E-07 | 6.044436 | Up | polysaccharide biosynthesis domain containing 1 |
| SHISA8 | 0.928111 | 0.000183 | 0.003407 | 3.813359 | Up | shisa family member 8 |
| SLC12A3 | 0.928031 | 0.000215 | 0.003837 | 3.769814 | Up | solute carrier family 12 member 3 |
| NLRC5 | 0.927988 | 1.08E-07 | 7.18E-06 | 5.514151 | Up | NLR family CARD domain containing 5 |
| CDCA2 | 0.927654 | 0.000292 | 0.004716 | 3.687852 | Up | cell division cycle associated 2 |
| H2AC7 | 0.926416 | 0.000186 | 0.003441 | 3.808835 | Up | H2A clustered histone 7 |
| TREX1 | 0.926046 | 4.23E-10 | 4.98E-08 | 6.572283 | Up | three prime repair exonuclease 1 |
| PDE1C | 0.925174 | 0.000443 | 0.006402 | 3.57285 | Up | phosphodiesterase 1C |
| TIMP4 | 0.924467 | 0.000108 | 0.002308 | 3.950523 | Up | TIMP metallopeptidase inhibitor 4 |
| MAP10 | 0.924329 | 2.73E-05 | 0.000786 | 4.294699 | Up | microtubule associated protein 10 |
| ESR1 | 0.921366 | 0.00205 | 0.018726 | 3.123955 | Up | estrogen receptor 1 |
| GPC4 | 0.920969 | 1.39E-05 | 0.000462 | 4.456212 | Up | glypican 4 |
| ENPP2 | 0.920702 | 5.75E-05 | 0.001389 | 4.111422 | Up | ectonucleotidepyrophosphatase/phosphodiesterase 2 |
| FGD2 | 0.920517 | 1.89E-06 | 8.47E-05 | 4.909976 | Up | FYVE, RhoGEF and PH domain containing 2 |
| RAMP1 | 0.919355 | 0.000273 | 0.004508 | 3.70592 | Up | receptor activity modifying protein 1 |
| GAL3ST4 | 0.919244 | 1.13E-05 | 0.000391 | 4.504813 | Up | galactose-3-O-sulfotransferase 4 |
| SLC26A1 | 0.918847 | 0.003472 | 0.026659 | 2.957844 | Up | solute carrier family 26 member 1 |
| GOLGA8Q | 0.917389 | 0.000119 | 0.002463 | 3.925848 | Up | golgin A8 family member Q |
| MTNR1A | 0.916886 | 0.000501 | 0.007012 | 3.538459 | Up | melatonin receptor 1A |
| H2AC20 | 0.91586 | 8.43E-10 | 8.98E-08 | 6.446779 | Up | H2A clustered histone 20 |
| TMEM42 | 0.914646 | 0.000104 | 0.002249 | 3.961002 | Up | transmembrane protein 42 |
| FBXO6 | 0.914172 | 3.33E-10 | 4E-08 | 6.615821 | Up | F-box protein 6 |
| ACY3 | 0.913938 | 0.000108 | 0.002308 | 3.950726 | Up | aminoacylase 3 |
| TOX | 0.91385 | 0.000877 | 0.010286 | 3.378409 | Up | thymocyte selection associated high mobility group box |
| ZNF682 | 0.913427 | 2.07E-05 | 0.000635 | 4.361432 | Up | zinc finger protein 682 |
| SIT1 | 0.913282 | 0.000936 | 0.010749 | 3.359276 | Up | signaling threshold regulating transmembrane adaptor 1 |
| STAT4 | 0.91098 | 0.000472 | 0.006686 | 3.554792 | Up | signal transducer and activator of transcription 4 |
| CD96 | 0.910756 | 9.02E-05 | 0.002012 | 3.997339 | Up | CD96 molecule |
| PPM1K | 0.909529 | 1.61E-15 | 4.07E-13 | 8.659512 | Up | protein phosphatase, Mg2+/Mn2+ dependent 1K |
| STAMBPL1 | 0.909026 | 2.22E-05 | 0.000675 | 4.34438 | Up | STAM binding protein like 1 |
| PET117 | 0.907766 | 2.19E-05 | 0.000668 | 4.34745 | Up | PET117 cytochrome c oxidase chaperone |
| RNASE1 | 0.906595 | 0.00051 | 0.007099 | 3.533442 | Up | ribonuclease A family member 1, pancreatic |
| IRS1 | 0.906406 | 2.12E-06 | 9.33E-05 | 4.885291 | Up | insulin receptor substrate 1 |
| CH25H | 0.906185 | 0.000379 | 0.005724 | 3.615754 | Up | cholesterol 25-hydroxylase |
| SEC16B | 0.905825 | 1.09E-05 | 0.000379 | 4.512972 | Up | SEC16 homolog B, endoplasmic reticulum export factor |
| CD33 | 0.905786 | 0.000285 | 0.00464 | 3.694461 | Up | CD33 molecule |
| EXOC3L4 | 0.905027 | 0.000219 | 0.003888 | 3.764555 | Up | exocyst complex component 3 like 4 |
| LGALS9 | 0.904739 | 1.58E-16 | 4.5E-14 | 9.020451 | Up | galectin 9 |
| TMSB10 | 0.903894 | 2.44E-11 | 3.68E-09 | 7.077674 | Up | thymosin beta 10 |
| CD74 | 0.902704 | 5.16E-10 | 5.84E-08 | 6.536563 | Up | CD74 molecule |
| TACR1 | 0.902382 | 0.001704 | 0.016385 | 3.180656 | Up | tachykinin receptor 1 |
| KIF5C | 0.901762 | 0.000353 | 0.005431 | 3.635467 | Up | kinesin family member 5C |
| APOL2 | 0.900875 | 1.39E-08 | 1.13E-06 | 5.9197 | Up | apolipoprotein L2 |
| APOC1 | 0.900681 | 0.001169 | 0.012487 | 3.293952 | Up | apolipoprotein C1 |
| TGM3 | -1.47834 | 0.000646 | 0.008303 | -3.46619 | Down | transglutaminase 3 |
| SCGB3A1 | -1.36843 | 9.64E-05 | 0.002119 | -3.98021 | Down | secretoglobin family 3A member 1 |
| IL1R2 | -1.2965 | 0.007366 | 0.04433 | -2.70757 | Down | interleukin 1 receptor type 2 |
| KLF15 | -1.20548 | 0.000549 | 0.007433 | -3.51242 | Down | Kruppel like factor 15 |
| FOSB | -1.19521 | 4.11E-05 | 0.001083 | -4.1945 | Down | FosB proto-oncogene, AP-1 transcription factor subunit |
| CD207 | -1.15391 | 0.000515 | 0.007122 | -3.53075 | Down | CD207 molecule |
| TULP2 | -1.15196 | 0.00269 | 0.022504 | -3.0392 | Down | TUB like protein 2 |
| RGPD2 | -1.07807 | 0.004861 | 0.033294 | -2.84797 | Down | RANBP2 like and GRIP domain containing 2 |
| SPRR3 | -1.07407 | 0.003937 | 0.028932 | -2.91721 | Down | small proline rich protein 3 |
| PDGFRB | -1.07186 | 0.001018 | 0.011372 | -3.33464 | Down | platelet derived growth factor receptor beta |
| CXCL8 | -1.06707 | 0.003106 | 0.024817 | -2.99357 | Down | C-X-C motif chemokine ligand 8 |
| MUC7 | -1.055 | 0.005844 | 0.037514 | -2.78645 | Down | mucin 7, secreted |
| BEST1 | -1.04013 | 0.002215 | 0.019712 | -3.10003 | Down | bestrophin 1 |
| CAMK1G | -1.03532 | 0.004051 | 0.029425 | -2.90789 | Down | calcium/calmodulin dependent protein kinase IG |
| SRSF12 | -1.01792 | 0.002864 | 0.023443 | -3.0194 | Down | serine and arginine rich splicing factor 12 |
| KRT13 | -1.00058 | 0.005628 | 0.03665 | -2.7991 | Down | keratin 13 |
| KLF2 | -0.98914 | 2.29E-08 | 1.82E-06 | -5.82208 | Down | Kruppel like factor 2 |
| PADI2 | -0.97338 | 0.0009 | 0.01046 | -3.37077 | Down | peptidyl arginine deiminase 2 |
| DGAT2 | -0.93702 | 0.005999 | 0.038269 | -2.77758 | Down | diacylglycerol O-acyltransferase 2 |
| THBD | -0.93625 | 0.003057 | 0.024538 | -2.99861 | Down | thrombomodulin |
| FOS | -0.89665 | 0.001294 | 0.013392 | -3.26371 | Down | Fos proto-oncogene, AP-1 transcription factor subunit |
| ADM | -0.87386 | 0.002488 | 0.021365 | -3.06371 | Down | adrenomedullin |
| C17orf107 | -0.86984 | 0.004991 | 0.033833 | -2.83924 | Down | chromosome 17 open reading frame 107 |
| MUC21 | -0.85619 | 0.002935 | 0.023868 | -3.01161 | Down | mucin 21, cell surface associated |
| HBEGF | -0.85044 | 0.002272 | 0.020037 | -3.0921 | Down | heparin binding EGF like growth factor |
| DCUN1D3 | -0.83153 | 5.2E-05 | 0.001291 | -4.13616 | Down | defective in cullinneddylation 1 domain containing 3 |
| ATP8 | -0.83079 | 7.05E-05 | 0.001661 | -4.05996 | Down | ATP synthase F0 subunit 8 |
| STATH | -0.81643 | 0.006636 | 0.041179 | -2.74335 | Down | statherin |
| BTBD19 | -0.81195 | 0.00112 | 0.012161 | -3.3067 | Down | BTB domain containing 19 |
| KLF7 | -0.81111 | 1.55E-05 | 0.000499 | -4.43075 | Down | Kruppel like factor 7 |
| DSC2 | -0.79948 | 0.000114 | 0.002392 | -3.9371 | Down | desmocollin 2 |
| ICAM3 | -0.79631 | 0.000294 | 0.00474 | -3.68565 | Down | intercellular adhesion molecule 3 |
| DUSP1 | -0.79617 | 4.07E-05 | 0.001076 | -4.19696 | Down | dual specificity phosphatase 1 |
| LOXHD1 | -0.79464 | 0.007319 | 0.044242 | -2.70977 | Down | lipoxygenase homology domains 1 |
| FTH1 | -0.79153 | 8.32E-09 | 7.23E-07 | -6.01825 | Down | ferritin heavy chain 1 |
| PNRC1 | -0.78805 | 3.64E-09 | 3.46E-07 | -6.17547 | Down | proline rich nuclear receptor coactivator 1 |
| NAMPT | -0.78417 | 0.005423 | 0.035674 | -2.81155 | Down | nicotinamidephosphoribosyltransferase |
| IL1R1 | -0.78387 | 8.07E-06 | 0.0003 | -4.58321 | Down | interleukin 1 receptor type 1 |
| EGR1 | -0.78372 | 0.000884 | 0.010347 | -3.37586 | Down | early growth response 1 |
| TREML2 | -0.76808 | 0.008381 | 0.048314 | -2.66284 | Down | triggering receptor expressed on myeloid cells like 2 |
| ADRB3 | -0.76534 | 0.001114 | 0.012113 | -3.30832 | Down | adrenoceptor beta 3 |
| ALPL | -0.76002 | 0.000591 | 0.007804 | -3.49159 | Down | alkaline phosphatase, biomineralization associated |
| AMPD2 | -0.75693 | 0.000279 | 0.004578 | -3.69953 | Down | adenosine monophosphate deaminase 2 |
| FCER1A | -0.75615 | 0.008706 | 0.049511 | -2.64956 | Down | Fc fragment of IgE receptor Ia |
| ARHGEF40 | -0.74698 | 0.000164 | 0.003152 | -3.84141 | Down | Rho guanine nucleotide exchange factor 40 |
| ZFP36 | -0.73956 | 0.00066 | 0.008423 | -3.46029 | Down | ZFP36 ring finger protein |
| EGR3 | -0.73321 | 0.008221 | 0.047609 | -2.66957 | Down | early growth response 3 |
| BTG2 | -0.73249 | 0.000255 | 0.004312 | -3.7242 | Down | BTG anti-proliferation factor 2 |
| ZSWIM4 | -0.73012 | 3.02E-06 | 0.000128 | -4.80662 | Down | zinc finger SWIM-type containing 4 |
| SEC14L6 | -0.72167 | 0.000339 | 0.005271 | -3.6463 | Down | SEC14 like lipid binding 6 |
| OR2B11 | -0.72005 | 0.005408 | 0.035648 | -2.81247 | Down | olfactory receptor family 2 subfamily B member 11 |
| TMEM71 | -0.71806 | 0.003467 | 0.026644 | -2.95839 | Down | transmembrane protein 71 |
| C5AR2 | -0.71584 | 0.000961 | 0.010963 | -3.35151 | Down | complement component 5a receptor 2 |
| FOSL2 | -0.71292 | 1.91E-05 | 0.00059 | -4.38054 | Down | FOS like 2, AP-1 transcription factor subunit |
| IER3 | -0.71147 | 3.81E-05 | 0.001027 | -4.21335 | Down | immediate early response 3 |
| ANPEP | -0.70961 | 4.8E-05 | 0.001224 | -4.1561 | Down | alanylaminopeptidase, membrane |
| ZNF219 | -0.70613 | 4.82E-06 | 0.000191 | -4.70116 | Down | zinc finger protein 219 |
| CASS4 | -0.70362 | 0.003212 | 0.025338 | -2.98288 | Down | Cas scaffold protein family member 4 |
| NCCRP1 | -0.70258 | 0.004876 | 0.033336 | -2.84698 | Down | NCCRP1, F-box associated domain containing |
| TRIB1 | -0.70221 | 0.001394 | 0.014121 | -3.24149 | Down | tribblespseudokinase 1 |
| PLEKHG2 | -0.69201 | 0.002074 | 0.018849 | -3.12036 | Down | pleckstrin homology and RhoGEF domain containing G2 |
| GK3P | -0.69041 | 0.005238 | 0.034877 | -2.82311 | Down | glycerol kinase 3 pseudogene |
| PFKFB3 | -0.68222 | 0.008618 | 0.049214 | -2.65313 | Down | 6-phosphofructo-2-kinase/fructose-2,6-biphosphatase 3 |
| LYPD3 | -0.66375 | 0.005074 | 0.034126 | -2.83371 | Down | LY6/PLAUR domain containing 3 |
| PYGL | -0.66104 | 3.2E-07 | 1.91E-05 | -5.29031 | Down | glycogen phosphorylase L |
| SCN2A | -0.65842 | 0.002758 | 0.022886 | -3.03127 | Down | sodium voltage-gated channel alpha subunit 2 |
| A2ML1 | -0.65205 | 0.008659 | 0.049346 | -2.65144 | Down | alpha-2-macroglobulin like 1 |
| LITAF | -0.64871 | 0.001169 | 0.012487 | -3.29393 | Down | lipopolysaccharide induced TNF factor |
| ABHD5 | -0.64294 | 6.84E-06 | 0.00026 | -4.62127 | Down | abhydrolase domain containing 5 |
| TSC22D3 | -0.63973 | 2.59E-06 | 0.000112 | -4.8405 | Down | TSC22 domain family member 3 |
| KREMEN1 | -0.63687 | 5.49E-05 | 0.001341 | -4.1227 | Down | kringle containing transmembrane protein 1 |
| NEDD9 | -0.63269 | 0.00293 | 0.023838 | -3.01216 | Down | neural precursor cell expressed, developmentally down-regulated 9 |
| ZNF442 | -0.63162 | 0.00247 | 0.021275 | -3.06605 | Down | zinc finger protein 442 |
| C5AR1 | -0.63159 | 0.000449 | 0.006465 | -3.56895 | Down | complement C5a receptor 1 |
| JUNB | -0.62547 | 0.00411 | 0.02962 | -2.90321 | Down | JunB proto-oncogene, AP-1 transcription factor subunit |
| CTDSP1 | -0.62535 | 0.000231 | 0.004041 | -3.7512 | Down | CTD small phosphatase 1 |
| IFRD1 | -0.61939 | 0.006351 | 0.039861 | -2.75827 | Down | interferon related developmental regulator 1 |
| GPR85 | -0.61483 | 0.004632 | 0.032097 | -2.86393 | Down | G protein-coupled receptor 85 |
| GYS1 | -0.61203 | 0.003398 | 0.026318 | -2.96483 | Down | glycogen synthase 1 |
| NDRG1 | -0.60552 | 0.000261 | 0.004375 | -3.71795 | Down | N-myc downstream regulated 1 |
| SOD2 | -0.6012 | 0.007303 | 0.044236 | -2.71055 | Down | superoxide dismutase 2 |
| ABTB1 | -0.60117 | 0.000526 | 0.007201 | -3.52478 | Down | ankyrin repeat and BTB domain containing 1 |
| DENND5A | -0.59892 | 0.003162 | 0.025098 | -2.98791 | Down | DENN domain containing 5A |
| ARRB2 | -0.59646 | 0.002132 | 0.01921 | -3.11193 | Down | arrestin beta 2 |
| OXSR1 | -0.59472 | 0.000116 | 0.002423 | -3.93181 | Down | oxidative stress responsive kinase 1 |
| ATG9B | -0.5935 | 0.000582 | 0.007714 | -3.49615 | Down | autophagy related 9B |
| SLC25A37 | -0.58982 | 0.008419 | 0.048459 | -2.66129 | Down | solute carrier family 25 member 37 |
| UPP1 | -0.58603 | 0.004169 | 0.02991 | -2.89856 | Down | uridinephosphorylase 1 |
| NR4A1 | -0.58478 | 0.001571 | 0.0154 | -3.20538 | Down | nuclear receptor subfamily 4 group A member 1 |
| GPCPD1 | -0.5831 | 0.000793 | 0.009574 | -3.40763 | Down | glycerophosphocholinephosphodiesterase 1 |
| LAMB3 | -0.5818 | 0.004963 | 0.033719 | -2.84107 | Down | laminin subunit beta 3 |
| IVNS1ABP | -0.58168 | 0.002567 | 0.021899 | -3.05395 | Down | influenza virus NS1A binding protein |
| IMPDH1 | -0.58013 | 6.18E-06 | 0.000239 | -4.64451 | Down | inosine monophosphate dehydrogenase 1 |
| METRNL | -0.57826 | 0.00137 | 0.013939 | -3.24672 | Down | meteorin like, glial cell differentiation regulator |
| TCN1 | -0.57779 | 0.006792 | 0.041813 | -2.73543 | Down | transcobalamin 1 |
| PLEKHG6 | -0.57288 | 0.001999 | 0.018352 | -3.13171 | Down | pleckstrin homology and RhoGEF domain containing G6 |
| BCL6 | -0.57252 | 0.000547 | 0.007418 | -3.51374 | Down | BCL6 transcription repressor |
| IL6R | -0.57246 | 0.000611 | 0.007963 | -3.48244 | Down | interleukin 6 receptor |
| ATP2B1 | -0.5717 | 0.001263 | 0.013194 | -3.27087 | Down | ATPase plasma membrane Ca2+ transporting 1 |
| CD44 | -0.5714 | 0.004209 | 0.030058 | -2.89537 | Down | CD44 molecule (Indian blood group) |
| KLLN | -0.56584 | 0.000684 | 0.008631 | -3.4499 | Down | killin, p53 regulated DNA replication inhibitor |
| PHLDA1 | -0.56267 | 0.001149 | 0.012344 | -3.29893 | Down | pleckstrin homology like domain family A member 1 |
| COL6A3 | -0.56185 | 0.006948 | 0.042489 | -2.72764 | Down | collagen type VI alpha 3 chain |
| TLE3 | -0.55702 | 0.001969 | 0.018212 | -3.13635 | Down | TLE family member 3, transcriptional corepressor |
| PPIF | -0.55572 | 0.004779 | 0.032882 | -2.85362 | Down | peptidylprolylisomerase F |
| MAP2K3 | -0.55394 | 0.000259 | 0.004355 | -3.71976 | Down | mitogen-activated protein kinase kinase 3 |
| GAB2 | -0.5507 | 0.006989 | 0.042671 | -2.72565 | Down | GRB2 associated binding protein 2 |
| NDEL1 | -0.54893 | 0.001085 | 0.011929 | -3.31587 | Down | nudE neurodevelopment protein 1 like 1 |
| OPA3 | -0.54683 | 4.36E-05 | 0.001134 | -4.18028 | Down | outer mitochondrial membrane lipid metabolism regulator OPA3 |
| JOSD1 | -0.54542 | 0.0011 | 0.012027 | -3.31182 | Down | Josephin domain containing 1 |
| PLB1 | -0.54522 | 0.005709 | 0.036981 | -2.79427 | Down | phospholipase B1 |
| REPS2 | -0.54355 | 1.35E-05 | 0.000451 | -4.46299 | Down | RALBP1 associated Eps domain containing 2 |
| TNFRSF10D | -0.54115 | 0.007126 | 0.04333 | -2.71899 | Down | TNF receptor superfamily member 10d |
| NFKBIA | -0.54054 | 0.007014 | 0.042809 | -2.72442 | Down | NFKB inhibitor alpha |
| PI3 | -0.53928 | 0.001287 | 0.013352 | -3.26536 | Down | peptidase inhibitor 3 |
| OR2L2 | -0.5338 | 0.008356 | 0.048213 | -2.6639 | Down | olfactory receptor family 2 subfamily L member 2 |
| GSTO2 | -0.53008 | 0.00374 | 0.027951 | -2.93388 | Down | glutathione S-transferase omega 2 |
| ST3GAL4 | -0.52905 | 0.003713 | 0.027827 | -2.93621 | Down | ST3 beta-galactoside alpha-2,3-sialyltransferase 4 |
| LRG1 | -0.52771 | 0.008031 | 0.046922 | -2.67768 | Down | leucine rich alpha-2-glycoprotein 1 |
| TP53INP2 | -0.52568 | 6.08E-06 | 0.000236 | -4.64824 | Down | tumor protein p53 inducible nuclear protein 2 |
| CEBPB | -0.52347 | 0.001649 | 0.015958 | -3.19071 | Down | CCAAT enhancer binding protein beta |
| USP32 | -0.52337 | 0.002161 | 0.019405 | -3.10772 | Down | ubiquitin specific peptidase 32 |
| SLC45A4 | -0.52164 | 0.000429 | 0.006246 | -3.58195 | Down | solute carrier family 45 member 4 |
| KATNBL1 | -0.51907 | 0.000403 | 0.00596 | -3.59889 | Down | katanin regulatory subunit B1 like 1 |
| PNPLA8 | -0.51402 | 0.00014 | 0.002803 | -3.88419 | Down | patatin like phospholipase domain containing 8 |
| MAPK13 | -0.51365 | 0.001167 | 0.01248 | -3.29447 | Down | mitogen-activated protein kinase 13 |
| FGD4 | -0.51363 | 0.002003 | 0.018366 | -3.13109 | Down | FYVE, RhoGEF and PH domain containing 4 |
| CD302 | -0.50813 | 0.006418 | 0.040184 | -2.75474 | Down | CD302 molecule |
| INPP5A | -0.50485 | 0.000956 | 0.01091 | -3.35321 | Down | inositol polyphosphate-5-phosphatase A |
| MYADM | -0.50339 | 4.83E-05 | 0.001228 | -4.15485 | Down | myeloid associated differentiation marker |
| GPX3 | -0.50185 | 0.00043 | 0.006248 | -3.5807 | Down | glutathione peroxidase 3 |
| PLK3 | -0.49613 | 0.004072 | 0.029525 | -2.90618 | Down | polo like kinase 3 |
| SHKBP1 | -0.49605 | 0.004754 | 0.032787 | -2.85533 | Down | SH3KBP1 binding protein 1 |
| SLC12A6 | -0.49412 | 0.004123 | 0.029697 | -2.90215 | Down | solute carrier family 12 member 6 |
| CNOT11 | -0.49107 | 0.002198 | 0.01962 | -3.10242 | Down | CCR4-NOT transcription complex subunit 11 |
| CPD | -0.488 | 0.000706 | 0.008859 | -3.44083 | Down | carboxypeptidase D |
| PRKDC | -0.48758 | 0.000507 | 0.007088 | -3.53471 | Down | protein kinase, DNA-activated, catalytic subunit |
| MBOAT2 | -0.48227 | 0.00078 | 0.009476 | -3.41214 | Down | membrane bound O-acyltransferase domain containing 2 |
| PID1 | -0.48159 | 0.00501 | 0.03391 | -2.83792 | Down | phosphotyrosine interaction domain containing 1 |
| RALGDS | -0.4811 | 3.26E-06 | 0.000136 | -4.78946 | Down | ral guanine nucleotide dissociation stimulator |
| RABGEF1 | -0.47821 | 0.000767 | 0.009382 | -3.41727 | Down | RAB guanine nucleotide exchange factor 1 |
| RASA2 | -0.47776 | 0.002722 | 0.022682 | -3.03548 | Down | RAS p21 protein activator 2 |
| PFKFB2 | -0.47681 | 0.000241 | 0.004154 | -3.73992 | Down | 6-phosphofructo-2-kinase/fructose-2,6-biphosphatase 2 |
| FAM53C | -0.47554 | 0.000236 | 0.004093 | -3.74529 | Down | family with sequence similarity 53 member C |
| MAML3 | -0.47245 | 0.000172 | 0.003273 | -3.82896 | Down | mastermind like transcriptional coactivator 3 |
| PPP1R15A | -0.46809 | 0.001775 | 0.016877 | -3.16816 | Down | protein phosphatase 1 regulatory subunit 15A |
| STAT5B | -0.46796 | 0.007994 | 0.046789 | -2.67931 | Down | signal transducer and activator of transcription 5B |
| LFNG | -0.46785 | 0.003845 | 0.028522 | -2.9249 | Down | LFNG O-fucosylpeptide 3-beta-N-acetylglucosaminyltransferase |
| FOXO4 | -0.46785 | 0.001931 | 0.017953 | -3.14236 | Down | forkhead box O4 |
| CEP68 | -0.46753 | 0.0017 | 0.016352 | -3.18146 | Down | centrosomal protein 68 |
| NIN | -0.46502 | 9.79E-06 | 0.000349 | -4.53826 | Down | ninein |
| MAN2B1 | -0.46365 | 1.22E-05 | 0.000419 | -4.48669 | Down | mannosidase alpha class 2B member 1 |
| BICD2 | -0.46337 | 4.37E-06 | 0.000176 | -4.72323 | Down | BICD cargo adaptor 2 |
| SPAG9 | -0.46006 | 0.000438 | 0.006344 | -3.57619 | Down | sperm associated antigen 9 |
| NIBAN1 | -0.45525 | 0.00108 | 0.011891 | -3.31741 | Down | niban apoptosis regulator 1 |
| MAP4K4 | -0.4546 | 0.000732 | 0.009111 | -3.43054 | Down | mitogen-activated protein kinase kinasekinasekinase 4 |
| MTURN | -0.45158 | 0.00053 | 0.007239 | -3.52257 | Down | maturin, neural progenitor differentiation regulator homolog |
| PER1 | -0.45133 | 0.000111 | 0.002353 | -3.94413 | Down | period circadian regulator 1 |
| CDC42EP3 | -0.44984 | 0.002209 | 0.019679 | -3.10081 | Down | CDC42 effector protein 3 |
| ETS2 | -0.44904 | 0.00178 | 0.01689 | -3.16738 | Down | ETS proto-oncogene 2, transcription factor |
| MANSC1 | -0.44787 | 0.006674 | 0.041333 | -2.74142 | Down | MANSC domain containing 1 |
| HIP1R | -0.44744 | 0.001356 | 0.013823 | -3.2496 | Down | huntingtin interacting protein 1 related |
| NR4A2 | -0.44532 | 0.003766 | 0.028091 | -2.93163 | Down | nuclear receptor subfamily 4 group A member 2 |
| HHLA1 | -0.44407 | 0.0004 | 0.005925 | -3.60075 | Down | HERV-H LTR-associating 1 |
| MAEA | -0.44405 | 7.51E-05 | 0.001753 | -4.04404 | Down | macrophage erythroblast attacher |
| SH3BP2 | -0.4436 | 0.005716 | 0.036994 | -2.79388 | Down | SH3 domain binding protein 2 |
| PACSIN2 | -0.44102 | 0.000822 | 0.00982 | -3.39702 | Down | protein kinase C and casein kinase substrate in neurons 2 |
| PICALM | -0.43931 | 0.000875 | 0.010274 | -3.37899 | Down | phosphatidylinositol binding clathrin assembly protein |
| INPP5K | -0.43911 | 0.002037 | 0.018636 | -3.12588 | Down | inositol polyphosphate-5-phosphatase K |
| MYH9 | -0.4341 | 0.001152 | 0.012346 | -3.29839 | Down | myosin heavy chain 9 |
| EIF3L | -0.43251 | 0.000558 | 0.007514 | -3.50807 | Down | eukaryotic translation initiation factor 3 subunit L |
| IRS2 | -0.43244 | 0.000379 | 0.005717 | -3.61633 | Down | insulin receptor substrate 2 |
| KDM4B | -0.42986 | 0.002404 | 0.020934 | -3.0745 | Down | lysine demethylase 4B |
| UBAP1 | -0.42924 | 6.92E-05 | 0.001636 | -4.06448 | Down | ubiquitin associated protein 1 |
| NBPF9 | -0.42916 | 0.003767 | 0.028091 | -2.93151 | Down | NBPF member 9 |
| SSH2 | -0.42856 | 0.004931 | 0.033584 | -2.84325 | Down | slingshot protein phosphatase 2 |
| ABHD12B | -0.42706 | 0.003739 | 0.027951 | -2.93394 | Down | abhydrolase domain containing 12B |
| PCDHB1 | -0.42614 | 0.005683 | 0.036912 | -2.79585 | Down | protocadherin beta 1 |
| SDCBP | -0.42408 | 0.001025 | 0.011418 | -3.33283 | Down | syndecan binding protein |
| RASSF5 | -0.42344 | 0.005139 | 0.034489 | -2.82948 | Down | Ras association domain family member 5 |
| TPD52L2 | -0.42323 | 1.35E-06 | 6.43E-05 | -4.98332 | Down | TPD52 like 2 |
| TPM4 | -0.42223 | 0.000186 | 0.003446 | -3.80814 | Down | tropomyosin 4 |
| SH3BP5 | -0.42121 | 0.002223 | 0.019753 | -3.09895 | Down | SH3 domain binding protein 5 |
| RAB31 | -0.42061 | 0.003548 | 0.02711 | -2.95095 | Down | RAB31, member RAS oncogene family |
| KLHL2 | -0.42058 | 0.000272 | 0.0045 | -3.70694 | Down | kelch like family member 2 |
| TNIP1 | -0.41705 | 0.004106 | 0.029617 | -2.90353 | Down | TNFAIP3 interacting protein 1 |
| TMCO3 | -0.41689 | 0.000515 | 0.007122 | -3.5305 | Down | transmembrane and coiled-coil domains 3 |
| SULF2 | -0.41581 | 0.004203 | 0.03005 | -2.89586 | Down | sulfatase 2 |
| EIF2D | -0.4151 | 0.000356 | 0.005455 | -3.63319 | Down | eukaryotic translation initiation factor 2D |
| MORC4 | -0.41342 | 0.0051 | 0.034269 | -2.83203 | Down | MORC family CW-type zinc finger 4 |
| OSBPL1A | -0.41211 | 0.003202 | 0.025288 | -2.98392 | Down | oxysterol binding protein like 1A |
| PTPN12 | -0.4119 | 0.003463 | 0.02664 | -2.95873 | Down | protein tyrosine phosphatase non-receptor type 12 |
| DOT1L | -0.41137 | 0.00764 | 0.045333 | -2.69498 | Down | DOT1 like histone lysine methyltransferase |
| C16orf72 | -0.40652 | 3.06E-05 | 0.000857 | -4.26681 | Down | chromosome 16 open reading frame 72 |
| ATP6V1B2 | -0.40561 | 0.00533 | 0.035283 | -2.81729 | Down | ATPase H+ transporting V1 subunit B2 |
| CDC42BPG | -0.40461 | 0.004898 | 0.033433 | -2.84545 | Down | CDC42 binding protein kinase gamma |
| CXCL16 | -0.40444 | 0.002518 | 0.02157 | -3.06005 | Down | C-X-C motif chemokine ligand 16 |
| PTEN | -0.40275 | 0.004977 | 0.033768 | -2.84017 | Down | phosphatase and tensin homolog |
| PELI1 | -0.40258 | 0.003722 | 0.027859 | -2.93541 | Down | pellino E3 ubiquitin protein ligase 1 |
| GXYLT2 | -0.4003 | 0.004254 | 0.030302 | -2.89195 | Down | glucosidexylosyltransferase 2 |
| MAP3K20 | -0.39795 | 0.007986 | 0.046763 | -2.67963 | Down | mitogen-activated protein kinase kinasekinase 20 |
| ERGIC1 | -0.39654 | 7.63E-05 | 0.001778 | -4.03977 | Down | endoplasmic reticulum-golgi intermediate compartment 1 |
| ACTN1 | -0.39536 | 0.007246 | 0.04399 | -2.71324 | Down | actinin alpha 1 |
| OSBPL8 | -0.39507 | 0.001578 | 0.015443 | -3.20399 | Down | oxysterol binding protein like 8 |
| NATD1 | -0.39499 | 0.001023 | 0.011406 | -3.33334 | Down | N-acetyltransferase domain containing 1 |
| RXRA | -0.39432 | 0.001122 | 0.012161 | -3.30607 | Down | retinoid X receptor alpha |
| SEC14L1 | -0.39413 | 0.000463 | 0.006583 | -3.56033 | Down | SEC14 like lipid binding 1 |
| SYAP1 | -0.38996 | 3.2E-05 | 0.000889 | -4.25609 | Down | synapse associated protein 1 |
| MAP3K11 | -0.38943 | 0.004389 | 0.030941 | -2.88168 | Down | mitogen-activated protein kinase kinasekinase 11 |
| NIBAN2 | -0.38681 | 0.006264 | 0.039467 | -2.76298 | Down | niban apoptosis regulator 2 |
| STEAP4 | -0.38453 | 0.003898 | 0.028768 | -2.92046 | Down | STEAP4 metalloreductase |
| CCNG2 | -0.384 | 0.001118 | 0.012157 | -3.30704 | Down | cyclin G2 |
| MAPKAPK2 | -0.38391 | 0.000604 | 0.00792 | -3.48542 | Down | MAPK activated protein kinase 2 |
| CCNQ | -0.38366 | 0.005702 | 0.036965 | -2.79468 | Down | cyclin Q |
| NUDT3 | -0.38348 | 0.004299 | 0.030544 | -2.88847 | Down | nudix hydrolase 3 |
| FRAT2 | -0.38298 | 0.007429 | 0.044476 | -2.70464 | Down | FRAT regulator of WNT signaling pathway 2 |
| TLCD2 | -0.3801 | 0.000278 | 0.004566 | -3.7005 | Down | TLC domain containing 2 |
| ERF | -0.37846 | 0.000263 | 0.004394 | -3.71566 | Down | ETS2 repressor factor |
| ZMIZ1 | -0.37414 | 0.004463 | 0.031289 | -2.87621 | Down | zinc finger MIZ-type containing 1 |
| TUBA4A | -0.374 | 0.00868 | 0.049431 | -2.65059 | Down | tubulin alpha 4a |
| SLC44A2 | -0.37399 | 0.000184 | 0.003417 | -3.81169 | Down | solute carrier family 44 member 2 |
| SORT1 | -0.37329 | 0.001334 | 0.013703 | -3.25471 | Down | sortilin 1 |
| LYST | -0.37303 | 0.005249 | 0.034931 | -2.82245 | Down | lysosomal trafficking regulator |
| PHF20L1 | -0.37185 | 0.004569 | 0.031759 | -2.86842 | Down | PHD finger protein 20 like 1 |
| PIM3 | -0.36926 | 0.00086 | 0.010147 | -3.3841 | Down | Pim-3 proto-oncogene, serine/threonine kinase |
| NDRG2 | -0.36889 | 0.000738 | 0.009146 | -3.42832 | Down | NDRG family member 2 |
| STX3 | -0.36716 | 0.00614 | 0.038903 | -2.76973 | Down | syntaxin 3 |
| ELL2 | -0.36667 | 0.00281 | 0.023157 | -3.02541 | Down | elongation factor for RNA polymerase II 2 |
| TPT1 | -0.36627 | 0.000566 | 0.007586 | -3.50374 | Down | tumor protein, translationally-controlled 1 |
| R3HDM4 | -0.36105 | 0.005795 | 0.037307 | -2.78924 | Down | R3H domain containing 4 |
| NUP98 | -0.35983 | 0.003404 | 0.026339 | -2.96426 | Down | nucleoporin 98 |
| FOXO3 | -0.35774 | 1.45E-05 | 0.000472 | -4.44649 | Down | forkhead box O3 |
| DDIT4 | -0.356 | 0.000593 | 0.007822 | -3.49059 | Down | DNA damage inducible transcript 4 |
| ZC3H3 | -0.35548 | 0.00825 | 0.047726 | -2.66836 | Down | zinc finger CCCH-type containing 3 |
| VPS37B | -0.35522 | 0.000756 | 0.009288 | -3.42149 | Down | VPS37B subunit of ESCRT-I |
| TBC1D14 | -0.35285 | 0.001397 | 0.014146 | -3.24078 | Down | TBC1 domain family member 14 |
| TFE3 | -0.35236 | 0.004362 | 0.030857 | -2.88372 | Down | transcription factor binding to IGHM enhancer 3 |
| MAP1LC3B | -0.3501 | 9.94E-06 | 0.000353 | -4.53486 | Down | microtubule associated protein 1 light chain 3 beta |
| FRMD4B | -0.34955 | 0.004943 | 0.033639 | -2.84242 | Down | FERM domain containing 4B |
| H3-3A | -0.34767 | 0.000148 | 0.002916 | -3.8696 | Down | H3.3 histone A |
| EEF2 | -0.34745 | 0.001604 | 0.015577 | -3.19905 | Down | eukaryotic translation elongation factor 2 |
| ASAH1 | -0.34715 | 5.29E-05 | 0.001309 | -4.13197 | Down | N-acylsphingosineamidohydrolase 1 |
| MAP3K8 | -0.34658 | 0.004083 | 0.029576 | -2.90532 | Down | mitogen-activated protein kinase kinasekinase 8 |
| SEMA4A | -0.34648 | 0.00489 | 0.033408 | -2.84599 | Down | semaphorin 4A |
| PXN | -0.34593 | 0.000275 | 0.004529 | -3.70385 | Down | paxillin |
| ARFGAP3 | -0.34357 | 0.000236 | 0.004098 | -3.7445 | Down | ADP ribosylation factor GTPase activating protein 3 |
| RNF144B | -0.34274 | 0.002253 | 0.019946 | -3.09477 | Down | ring finger protein 144B |
| UBE2R2 | -0.34042 | 0.00023 | 0.004029 | -3.75232 | Down | ubiquitin conjugating enzyme E2 R2 |
| NBPF14 | -0.34008 | 0.001432 | 0.014368 | -3.23326 | Down | NBPF member 14 |
| RHOQ | -0.34 | 0.002276 | 0.020061 | -3.09155 | Down | ras homolog family member Q |
| IRF2BP2 | -0.33749 | 0.000184 | 0.003417 | -3.81134 | Down | interferon regulatory factor 2 binding protein 2 |
| GABARAPL1 | -0.33722 | 0.000609 | 0.007959 | -3.48311 | Down | GABA type A receptor associated protein like 1 |
| RAB3D | -0.33666 | 0.002283 | 0.020095 | -3.09067 | Down | RAB3D, member RAS oncogene family |
| PINK1 | -0.33551 | 0.000119 | 0.002463 | -3.92585 | Down | PTEN induced kinase 1 |
| SERINC3 | -0.33472 | 0.000521 | 0.007167 | -3.52743 | Down | serine incorporator 3 |
| PABPC1 | -0.33444 | 0.000567 | 0.007589 | -3.50338 | Down | poly(A) binding protein cytoplasmic 1 |
| ZNF185 | -0.33247 | 0.006736 | 0.041572 | -2.73826 | Down | zinc finger protein 185 with LIM domain |
| UBE2D1 | -0.33144 | 0.004895 | 0.033426 | -2.84567 | Down | ubiquitin conjugating enzyme E2 D1 |
| CPEB4 | -0.32973 | 0.003715 | 0.027827 | -2.93608 | Down | cytoplasmic polyadenylation element binding protein 4 |
| KLHL21 | -0.32456 | 0.000783 | 0.009487 | -3.41137 | Down | kelch like family member 21 |
| IDI1 | -0.31705 | 0.000668 | 0.008477 | -3.45662 | Down | isopentenyl-diphosphate delta isomerase 1 |
| GABARAP | -0.31514 | 0.000191 | 0.00352 | -3.801 | Down | GABA type A receptor-associated protein |
| IQGAP1 | -0.3113 | 0.001355 | 0.013817 | -3.24999 | Down | IQ motif containing GTPase activating protein 1 |
| TTPAL | -0.31108 | 0.005399 | 0.035625 | -2.81302 | Down | alpha tocopherol transfer protein like |
| BLOC1S6 | -0.31029 | 0.003798 | 0.028251 | -2.92889 | Down | biogenesis of lysosomal organelles complex 1 subunit 6 |
| ZFP36L1 | -0.30501 | 0.006474 | 0.040439 | -2.75178 | Down | ZFP36 ring finger protein like 1 |
| RIOK3 | -0.30287 | 0.002094 | 0.018965 | -3.11748 | Down | RIO kinase 3 |
| ADIPOR1 | -0.30265 | 3.3E-06 | 0.000137 | -4.78639 | Down | adiponectin receptor 1 |
| PPARD | -0.3019 | 0.003562 | 0.027186 | -2.94963 | Down | peroxisome proliferator activated receptor delta |
| SLC25A40 | -0.30097 | 0.005503 | 0.03599 | -2.80666 | Down | solute carrier family 25 member 40 |
| YPEL3 | -0.29906 | 0.007547 | 0.045029 | -2.69923 | Down | yippee like 3 |
| SNX18 | -0.29867 | 0.002638 | 0.02223 | -3.04539 | Down | sorting nexin 18 |
| WDR1 | -0.2972 | 0.001094 | 0.011994 | -3.31345 | Down | WD repeat domain 1 |
| MSL1 | -0.29603 | 0.004073 | 0.029525 | -2.90612 | Down | MSL complex subunit 1 |
| TKT | -0.29601 | 0.001792 | 0.016955 | -3.1653 | Down | transketolase |
| EIF4EBP2 | -0.29523 | 0.001382 | 0.014026 | -3.24409 | Down | eukaryotic translation initiation factor 4E binding protein 2 |
| MTMR3 | -0.29205 | 0.007323 | 0.044242 | -2.70958 | Down | myotubularin related protein 3 |
| EIF1 | -0.29128 | 7.23E-05 | 0.001698 | -4.05365 | Down | eukaryotic translation initiation factor 1 |
| RANBP2 | -0.28835 | 0.00612 | 0.038791 | -2.77084 | Down | RAN binding protein 2 |
| GLUL | -0.28774 | 0.001254 | 0.013146 | -3.27311 | Down | glutamate-ammonia ligase |
| DNM2 | -0.28768 | 0.005212 | 0.0348 | -2.82482 | Down | dynamin 2 |
| PRKCD | -0.28536 | 0.00501 | 0.03391 | -2.83797 | Down | protein kinase C delta |
| USF2 | -0.28423 | 0.000968 | 0.010992 | -3.34957 | Down | upstream transcription factor 2, c-fos interacting |
| DOP1B | -0.28185 | 0.007237 | 0.043955 | -2.71365 | Down | DOP1 leucine zipper like protein B |
| STK38L | -0.27657 | 0.006341 | 0.039827 | -2.7588 | Down | serine/threonine kinase 38 like |
| TFDP1 | -0.27631 | 0.001671 | 0.016116 | -3.18663 | Down | transcription factor Dp-1 |
| ZSWIM6 | -0.2704 | 0.006811 | 0.04191 | -2.73444 | Down | zinc finger SWIM-type containing 6 |
| YPEL5 | -0.26608 | 0.000443 | 0.006402 | -3.57266 | Down | yippee like 5 |
| CAMK1D | -0.26524 | 0.004032 | 0.029403 | -2.90946 | Down | calcium/calmodulin dependent protein kinase ID |
| GNAQ | -0.2631 | 0.003636 | 0.027507 | -2.94302 | Down | G protein subunit alpha q |
| SGK1 | -0.26184 | 0.004447 | 0.03122 | -2.87736 | Down | serum/glucocorticoid regulated kinase 1 |
| UBE2H | -0.26171 | 0.003338 | 0.026007 | -2.97052 | Down | ubiquitin conjugating enzyme E2 H |
| RNF10 | -0.2591 | 0.003397 | 0.026318 | -2.96489 | Down | ring finger protein 10 |
| CMIP | -0.25797 | 0.000785 | 0.009508 | -3.41052 | Down | c-Maf inducing protein |
| WBP2 | -0.25642 | 0.0003 | 0.004812 | -3.68016 | Down | WW domain binding protein 2 |
| IDS | -0.25232 | 0.002736 | 0.022752 | -3.03385 | Down | iduronate 2-sulfatase |
| ANXA11 | -0.2512 | 0.005195 | 0.034762 | -2.82586 | Down | annexin A11 |
| RBM47 | -0.2503 | 0.004286 | 0.030478 | -2.88947 | Down | RNA binding motif protein 47 |
| KAT8 | -0.24712 | 0.004379 | 0.030931 | -2.88245 | Down | lysine acetyltransferase 8 |
| RNF141 | -0.24646 | 0.001755 | 0.016717 | -3.17162 | Down | ring finger protein 141 |
| WDR45 | -0.24604 | 0.004381 | 0.030937 | -2.88224 | Down | WD repeat domain 45 |
| CASC3 | -0.24371 | 0.002782 | 0.023038 | -3.02854 | Down | CASC3 exon junction complex subunit |
| RAB7A | -0.24268 | 0.002251 | 0.019946 | -3.09494 | Down | RAB7A, member RAS oncogene family |
| MLLT1 | -0.24026 | 0.007512 | 0.044855 | -2.70083 | Down | MLLT1 super elongation complex subunit |
| DNAJC5 | -0.23976 | 0.002324 | 0.02033 | -3.08502 | Down | DnaJ heat shock protein family (Hsp40) member C5 |
| B4GALT1 | -0.23721 | 0.004137 | 0.029747 | -2.90107 | Down | beta-1,4-galactosyltransferase 1 |
| ZCCHC14 | -0.23514 | 0.006633 | 0.041176 | -2.7435 | Down | zinc finger CCHC-type containing 14 |
| BZW1 | -0.23476 | 0.003892 | 0.028748 | -2.92093 | Down | basic leucine zipper and W2 domains 1 |
| RYBP | -0.23443 | 0.005895 | 0.037786 | -2.78349 | Down | RING1 and YY1 binding protein |
| GNB2 | -0.23408 | 0.001941 | 0.018019 | -3.14088 | Down | G protein subunit beta 2 |
| EEF1D | -0.23373 | 0.00569 | 0.036912 | -2.79544 | Down | eukaryotic translation elongation factor 1 delta |
| MAPK1 | -0.23171 | 0.002436 | 0.021124 | -3.07031 | Down | mitogen-activated protein kinase 1 |
| KLHL24 | -0.23087 | 0.008307 | 0.048029 | -2.66594 | Down | kelch like family member 24 |
| PPP4R1 | -0.23022 | 0.005951 | 0.038035 | -2.78032 | Down | protein phosphatase 4 regulatory subunit 1 |
| CHP1 | -0.22577 | 0.002763 | 0.02291 | -3.03076 | Down | calcineurin like EF-hand protein 1 |
| CDK16 | -0.21855 | 0.005028 | 0.033925 | -2.83678 | Down | cyclin dependent kinase 16 |
| FDPS | -0.21774 | 0.003361 | 0.026106 | -2.96837 | Down | farnesyldiphosphate synthase |
| OSBPL2 | -0.21684 | 0.006654 | 0.041245 | -2.74241 | Down | oxysterol binding protein like 2 |
| PRKACA | -0.21604 | 0.005 | 0.033866 | -2.83864 | Down | protein kinase cAMP-activated catalytic subunit alpha |
| LAMP1 | -0.20879 | 0.006045 | 0.038424 | -2.775 | Down | lysosomal associated membrane protein 1 |
| DNAJB1 | -0.20621 | 0.00732 | 0.044242 | -2.70975 | Down | DnaJ heat shock protein family (Hsp40) member B1 |
| CCNI | -0.20561 | 0.005766 | 0.037193 | -2.79098 | Down | cyclin I |
| RNASEK | -0.20408 | 0.007377 | 0.04435 | -2.70706 | Down | ribonuclease K |

**Table 2** The enriched GO terms of the up and down regulated differentially expressed genes

| **GO ID** | **CATEGORY** | **GO Name** | **Adjusted**  **p value** | **negative**  **log10**  **of adjusted**  **p value** | **Gene Count** | **Gene** |
| --- | --- | --- | --- | --- | --- | --- |
| **Up regulated genes** | | | | | | |
| GO:0002376 | BP | immune system process | 1.62E-50 | 49.78916342 | 207 | CXCL11,CXCL10,ISG15,IFI6,CXCL9,IFI44L,IFIT3,IFIT1,BST2,IFI27,RSAD2,OASL,SERPING1,IFIT2,USP18,UBD,IFI44,CCL8,LAG3,PLVAP,OAS3,IFITM1,GZMB,RTP4,IFITM3,OAS2,OAS1,DDX60,CXCL13,CCL2,PRF1,IDO1,CD80,DDX58,HERC5,GZMA,C1QA,BATF2,MX1,C1QB,IFIT5,PTPRO,IL4I1,IFNG,H3C7,IFIH1,JCHAIN,TICAM2,CD2,H3C12,MUC13,IGLL5,ACOD1,ADAMDEC1,PARP9,C1QC,STAT1,KLRC2,CCR5,PARP14,PTN,BTN3A3,SAA1,C4B,PDCD1LG2,H2BC11,IFI35,ADCY5,HLA-DOB,MT2A,FPR3,DPP4,TNFSF10,H3C2,TBX21,PSMB9,CAMK4,FASLG,DHX58,HLA-DMB,H4C13,BTN3A1,CD3D,CD3E,CMKLR1,ICOS,GBP1,H4C14,GRAP2,XAF1,TRIM22,LAMP3,H3C3,NMI,EOMES,THEMIS,PDCD1,ISG20,NT5C3A,SUCNR1,IRF7,HLA-F,GPR171,ADGRF5,DEFB1,SHFL,CD8A,H3C8,HLA-G,CD8B,MDK,GGT1,H3C10,IRF4,SIRPG,C4A,EIF2AK2,SH2D1A,ZAP70,TIGIT,RGS1,ZNF683,KIF15,HLA-DRB5,GSDMB,UBASH3A,CD244,C1R,STAP1,KLRC1,PSME2,PML,SLAMF8,CENPE,HLA-DMA,SLC16A1,TAP1,CTLA4,CD38,TLR3,CX3CL1,CD247,GBP5,NOS2,H2BC7,SFRP1,HLA-DRB1,AIM2,CFB,TLR7,MARCHF1,TNFSF13B,SLAMF1,DTX3L,IRF8,SLAMF7,CCR1,GFI1,IL7,ZBP1,DOK2,FCGR3A,BPIFA1,RHEX,TMEM176B,CD300E,PSMB8,H4C11,BCL11B,TAP2,BATF3,CD48,APOD,MX2,PYHIN1,APOL1,H2BC10,GBP4,H4C6,CD40LG,BACH2,CD226,CXCL12,TNFRSF13B,STAT2,CFH,CRTAM,RARRES2,GNLY,RNASE6,CARD11,H3C13,IL15RA,CPLX2,FYN,IL32,H2BC6,NLRC5,ENPP2,RAMP1,TOX,SIT1,CD96,CH25H,CD33,LGALS9,CD74 |
| GO:0050896 | BP | response to stimulus | 1.87E-17 | 16.72713193 | 286 | CXCL11,CXCL10,ISG15,IFI6,SIGLEC1,CXCL9,IFI44L,IFIT3,IFIT1,BST2,IFI27,RSAD2,OASL,SERPING1,IFIT2,USP18,UBD,IFI44,CMPK2,CCL8,LAG3,PLVAP,OAS3,IFITM1,RUFY4,GZMB,RTP4,IFITM3,OAS2,PTPRCAP,SMTNL1,OAS1,DDX60,TFEC,CXCL13,CCL2,PRF1,SAA4,CD80,DDX58,HERC5,GZMA,C1QA,BATF2,MX1,C1QB,IFIT5,ACE2,PTPRO,IL4I1,IFNG,APOL3,SAA2-SAA4,H3C7,IFIH1,JCHAIN,TICAM2,CD2,LRRN2,H3C12,MUC13,IGLL5,ACOD1,ADAMDEC1,PARP9,C1QC,MYBL2,STAT1,KLRC2,FAP,CCR5,PARP14,PTN,UBE2L6,BTN3A3,SAA1,SAA2,C4B,LGR6,LY6E,PDCD1LG2,H2BC11,IFI35,ADCY5,HLA-DOB,MT2A,FPR3,DPP4,TNFSF10,H3C2,TBX21,PPAN-P2RY11,FAM83A,PSMB9,CAMK4,FASLG,DHX58,HLA-DMB,ABCB1,H4C13,RNF43,BTN3A1,CD3D,CD3E,P2RX5,CMKLR1,ICOS,GBP1,H4C14,GRAP2,XAF1,PRLR,TRIM22,TPX2,LAMP3,IQSEC3,H3C3,NMI,EOMES,COL5A1,THEMIS,PDCD1,ISG20,NT5C3A,SUCNR1,IRF7,HLA-F,MGAT3,GABRB2,HESX1,LGALS3BP,GPR171,ADGRF5,DEFB1,SHFL,CD8A,H3C8,GRAP,HLA-G,HSH2D,CD8B,MDK,GGT1,H3C10,CACNA2D1,RBM11,IRF4,SIRPG,C4A,EIF2AK2,SH2D1A,ZAP70,RGS1,ZNF683,HLA-DRB5,GSDMB,UBASH3A,CD244,RORB,IGFBP4,MUSK,C1R,STAP1,KLRC1,PSME2,PML,SLAMF8,HLA-DMA,SLC16A1,TAP1,CTLA4,CD38,TLR3,POSTN,CX3CL1,PLLP,CD247,GBP5,NOS2,H2BC7,ASPM,DUOXA2,SFRP1,HLA-DRB1,DGKI,ABI3,SP140,AIM2,CFB,TLR7,MARCHF1,TNFSF13B,SLAMF1,DTX3L,BUB1,IRF8,SLAMF7,CCR1,CYP19A1,GFI1,NUPR1,UBE2T,TPSAB1,IL7,PERM1,ZBP1,DOK2,SH2D2A,FCGR3A,BPIFA1,RHEX,CD300E,MLC1,PSMB8,H4C11,BCL11B,TAP2,TEDC2,BATF3,CD48,CYP2C18,SYT13,APOD,BCL2L14,MX2,PYHIN1,PROX1,APOL1,H2BC10,GBP4,H4C6,CD40LG,BACH2,CD226,CXCL12,TNFRSF13B,STAT2,CFH,TGM2,WASF1,CRTAM,CASQ2,RARRES2,GNLY,RNASE6,CDC7,GNGT2,GRIK1,CARD11,COL4A3,H3C13,IL15RA,CPLX2,FYN,RASGRF2,IL32,HASPIN,RYR1,PLEK2,BUB1B,PAK1IP1,ADGRB2,H2BC6,SHISA8,NLRC5,PDE1C,TIMP4,ESR1,GPC4,ENPP2,FGD2,RAMP1,MTNR1A,FBXO6,SIT1,STAT4,CD96,IRS1,CH25H,CD33,LGALS9,CD74,TACR1,KIF5C,APOL2,APOC1 |
| GO:0071944 | CC | cell periphery | 2.30055E-05 | 4.638168411 | 177 | CXCL10,IFI6,SIGLEC1,CXCL9,BST2,SERPING1,LAG3,PLVAP,OAS3,IFITM1,GZMB,IFITM3,PTPRCAP,PRF1,OTOF,CD80,DDX58,GZMA,C1QA,MX1,IFIT5,ACE2,TRO,PTPRO,KCNA3,NKG7,IL4I1,SLC38A5,TICAM2,CD2,LRRN2,SLC43A1,MUC13,IGLL5,ADAMDEC1,C1QC,KLRC2,FAP,CCR5,PARP14,PTN,BTN3A3,C4B,LGR6,LY6E,PDCD1LG2,H2BC11,BOLL,ADCY5,HLA-DOB,FPR3,DPP4,TNFSF10,HAPLN3,PPAN-P2RY11,FASLG,HLA-DMB,ABCB1,MS4A4A,RNF43,BTN3A1,CD3D,CD3E,IGSF9B,P2RX5,CMKLR1,ICOS,CALHM6,GBP1,GRAP2,PRLR,LAMP3,COL5A1,PDCD1,SUCNR1,PLK4,HLA-F,GABRB2,LGALS3BP,GPR171,ADGRF5,ATP10A,CD8A,HLA-G,CD8B,MDK,GGT1,CACNA2D1,SIRPG,C4A,ZAP70,TIGIT,RGS1,PCDH17,HLA-DRB5,GSDMB,CD244,MUSK,KLRC1,CD69,RIMBP2,HLA-DMA,SLC16A1,CTLA4,CD38,TLR3,POSTN,CX3CL1,VAMP5,CD247,MYO7A,NOS2,ASPM,DUOXA2,SFRP1,HLA-DRB1,DGKI,CFB,TLR7,MARCHF1,TNFSF13B,SLAMF1,SLAMF7,CCR1,TPSAB1,IL7,FCGR3A,RHEX,CD300E,MLC1,CD48,SYT13,MX2,CLCNKB,KCNA2,GBP4,CD40LG,CD226,CXCL12,TNFRSF13B,DPP10,STAT2,CD72,TGM2,CRTAM,RARRES2,CD52,GNGT2,GRIK1,CARD11,COL4A3,IL15RA,FXYD6,FYN,RASGRF2,RYR1,PLEK2,ADGRB2,SHISA8,SLC12A3,TIMP4,ESR1,GPC4,ENPP2,FGD2,RAMP1,SLC26A1,MTNR1A,ACY3,SIT1,CD96,IRS1,CD33,EXOC3L4,LGALS9,CD74,TACR1 |
| GO:0005886 | CC | plasma membrane | 0.000248423 | 3.604808763 | 160 | CXCL10,IFI6,SIGLEC1,CXCL9,BST2,LAG3,PLVAP,OAS3,IFITM1,GZMB,IFITM3,PTPRCAP,PRF1,OTOF,CD80,DDX58,GZMA,MX1,IFIT5,ACE2,TRO,PTPRO,KCNA3,NKG7,IL4I1,SLC38A5,TICAM2,CD2,SLC43A1,MUC13,IGLL5,KLRC2,FAP,CCR5,PARP14,PTN,BTN3A3,C4B,LGR6,LY6E,PDCD1LG2,H2BC11,BOLL,ADCY5,HLA-DOB,FPR3,DPP4,TNFSF10,PPAN-P2RY11,FASLG,HLA-DMB,ABCB1,MS4A4A,RNF43,BTN3A1,CD3D,CD3E,IGSF9B,P2RX5,CMKLR1,ICOS,CALHM6,GBP1,GRAP2,PRLR,LAMP3,PDCD1,SUCNR1,PLK4,HLA-F,GABRB2,GPR171,ADGRF5,ATP10A,CD8A,HLA-G,CD8B,GGT1,CACNA2D1,SIRPG,C4A,ZAP70,TIGIT,RGS1,PCDH17,HLA-DRB5,GSDMB,CD244,MUSK,KLRC1,CD69,RIMBP2,HLA-DMA,SLC16A1,CTLA4,CD38,TLR3,CX3CL1,VAMP5,CD247,MYO7A,NOS2,ASPM,DUOXA2,SFRP1,HLA-DRB1,DGKI,CFB,TLR7,MARCHF1,TNFSF13B,SLAMF1,SLAMF7,CCR1,FCGR3A,RHEX,CD300E,MLC1,CD48,SYT13,MX2,CLCNKB,KCNA2,GBP4,CD40LG,CD226,CXCL12,TNFRSF13B,DPP10,STAT2,CD72,TGM2,CRTAM,CD52,GNGT2,GRIK1,CARD11,IL15RA,FXYD6,FYN,RASGRF2,RYR1,PLEK2,ADGRB2,SHISA8,SLC12A3,ESR1,GPC4,ENPP2,FGD2,RAMP1,SLC26A1,MTNR1A,ACY3,SIT1,CD96,IRS1,CD33,CD74,TACR1 |
| GO:0005488 | MF | binding | 0.000131237 | 3.881945019 | 385 | CXCL11,CXCL10,ISG15,IFI6,SIGLEC1,CXCL9,IFI44L,IFIT3,IFIT1,BST2,IFI27,RSAD2,OASL,SERPING1,IFIT2,USP18,EPSTI1,UBD,IFI44,CMPK2,KLHDC7B,CCL8,LAG3,PLVAP,OAS3,IFITM1,RUFY4,PLAAT2,GZMB,RTP4,IFITM3,OAS2,GIMAP7,PTPRCAP,ANXA10,SMTNL1,OAS1,DDX60,TFEC,CXCL13,CCL2,PRF1,SAA4,ETV7,OTOF,SAMD9L,GIMAP8,IDO1,CD80,GIMAP5,DDX58,HERC5,GZMA,C1QA,BATF2,MX1,C1QB,IFIT5,GIMAP6,ACE2,TRO,PRRX2,PTPRO,KCNA3,NKG7,GIMAP4,IL4I1,H2AC16,IFNG,APOL3,PCSK5,H3C7,H2AC19,IFIH1,JCHAIN,TICAM2,CD2,LRRN2,ZNF831,H3C12,SLC43A1,MUC13,IGLL5,ADAMDEC1,PARP9,C1QC,MYBL2,STAT1,KLRC2,UBQLNL,H2AC14,FAP,CCR5,PARP14,NRSN1,PTN,UBE2L6,BTN3A3,SAA1,SAA2,H1-5,C4B,LGR6,LY6E,LAP3,SAMD9,PDCD1LG2,H2BC11,BOLL,IFI35,ADCY5,HLA-DOB,MT2A,WDR74,FBXO39,DPP4,TNFSF10,H3C2,HAPLN3,TBX21,H2BC14,H2BC17,H2AC12,PPAN-P2RY11,FAM83A,PSMB9,CAMK4,FASLG,DHX58,HLA-DMB,ABCB1,H4C13,MS4A4A,RNF43,BTN3A1,CD3D,CD3E,IGSF9B,P2RX5,CMKLR1,ICOS,CALHM6,H2BC3,GBP1,H4C14,GRAP2,XAF1,PRLR,TRIM22,TPX2,IQSEC3,H3C3,NMI,EOMES,COL5A1,THEMIS,PDCD1,ISG20,NT5C3A,SUCNR1,PSAT1,PLK4,IRF7,HLA-F,MGAT3,PPM1J,HESX1,LGALS3BP,ADGRF5,PARP12,ATP10A,DEFB1,SHFL,CD8A,H3C8,GRAP,KRT39,HLA-G,HSH2D,CD8B,C17ORF67,FUOM,MDK,GGT1,H3C10,CACNA2D1,RBM11,IRF4,SIRPG,ASCL3,C4A,EIF2AK2,SH2D1A,ZAP70,TIGIT,RGS1,ZNF683,KIF15,PCDH17,HLA-DRB5,GSDMB,UBASH3A,H2AC17,CD244,RORB,IGFBP4,MUSK,C1R,STAP1,KLRC1,CD69,PSME2,PML,SLAMF8,AGR3,LKAAEAR1,SPATS2L,RIMBP2,WARS1,CENPE,HLA-DMA,SLC16A1,TAP1,H2AC13,CTLA4,CD38,TLR3,POSTN,CX3CL1,VAMP5,HDX,PLLP,CD247,GBP5,MYO7A,ENDOD1,NOS2,MUC19,TMEM187,H2BC7,ASPM,DUOXA2,SFRP1,CDCA7,HLA-DRB1,DGKI,ABI3,SP140,TRANK1,AIM2,CFB,TLR7,MARCHF1,TNFSF13B,SLAMF1,DTX3L,BUB1,IRF8,PLAAT4,SLAMF7,CCR1,CYP19A1,GFI1,NUPR1,UBE2T,TPSAB1,NUSAP1,IL7,ZBP1,DOK2,SH2D2A,FCGR3A,BPIFA1,B4GALT6,RHEX,TMEM176B,FAM241B,CD300E,MLC1,PSMB8,H4C11,BCL11B,TAP2,ZNF311,TEDC2,BATF3,CD48,UBQLN3,CYP2C18,SYT13,APOD,BCL2L14,MX2,GIMAP1,PYHIN1,PROX1,MB21D2,CCDC103,ZNF367,APOL1,H2BC10,CLCNKB,KCNA2,GBP4,H4C6,CD40LG,BACH2,SP110,CD226,CXCL12,TNFRSF13B,DPP10,STAT2,CD72,CFH,TGM2,WASF1,H2BC13,CRTAM,CASQ2,RARRES2,GNLY,DUS4L-BCAP29,RNASE6,CDC7,GNGT2,CARD11,COL4A3,H3C13,IL15RA,FXYD6,CPLX2,SLFN5,FYN,RASGRF2,IL32,HASPIN,HELZ2,RYR1,MS4A7,TUBA8,PLEK2,BUB1B,PAK1IP1,ADGRB2,H2BC6,ZSCAN23,SLC12A3,NLRC5,H2AC7,PDE1C,TIMP4,MAP10,ESR1,GPC4,ENPP2,FGD2,RAMP1,GAL3ST4,SLC26A1,GOLGA8Q,MTNR1A,H2AC20,TMEM42,FBXO6,ACY3,TOX,ZNF682,SIT1,STAT4,PPM1K,STAMBPL1,RNASE1,IRS1,CH25H,SEC16B,CD33,EXOC3L4,LGALS9,TMSB10,CD74,TACR1,KIF5C,APOL2,APOC1 |
| GO:0042802 | MF | identical protein binding | 0.000745002 | 3.127842328 | 72 | IFIT3,BST2,IFI27,PLVAP,GIMAP7,PRF1,DDX58,GZMA,MX1,ACE2,PTPRO,IFIH1,JCHAIN,CD2,MUC13,STAT1,FAP,CCR5,IFI35,DPP4,TNFSF10,FAM83A,CD3D,CD3E,GBP1,TRIM22,NMI,PSAT1,PLK4,HESX1,DEFB1,SHFL,HLA-G,RBM11,EIF2AK2,TIGIT,CD69,PSME2,PML,SLAMF8,WARS1,SLC16A1,TAP1,CD38,TLR3,CD247,GBP5,MYO7A,NOS2,H2BC7,SFRP1,ABI3,AIM2,SLAMF1,SLAMF7,TPSAB1,ZBP1,MLC1,CCDC103,H2BC10,GBP4,STAT2,CFH,TGM2,CRTAM,CASQ2,FYN,H2BC6,ESR1,ACY3,STAT4,CD74 |
| **Down regulated genes** | | | | | | |
| GO:0007154 | BP | cell communication | 2.39175E-06 | 5.621283626 | 165 | SCGB3A1,IL1R2,KLF15,PDGFRB,CXCL8,MUC7,KLF2,PADI2,DGAT2,FOS,ADM,MUC21,HBEGF,KLF7,DSC2,ICAM3,DUSP1,NAMPT,IL1R1,EGR1,ADRB3,FCER1A,ZFP36,EGR3,BTG2,OR2B11,C5AR2,IER3,ANPEP,ZNF219,CASS4,TRIB1,PLEKHG2,SCN2A,LITAF,KREMEN1,NEDD9,C5AR1,JUNB,GPR85,NDRG1,SOD2,ARRB2,OXSR1,UPP1,NR4A1,IVNS1ABP,METRNL,BCL6,IL6R,ATP2B1,CD44,TLE3,PPIF,MAP2K3,GAB2,NDEL1,REPS2,TNFRSF10D,NFKBIA,OR2L2,LRG1,CEBPB,MAPK13,FGD4,INPP5A,MYADM,PLK3,SHKBP1,SLC12A6,CNOT11,PRKDC,PID1,RALGDS,RASA2,PFKFB2,MAML3,PPP1R15A,STAT5B,LFNG,FOXO4,SPAG9,MAP4K4,MTURN,PER1,CDC42EP3,HIP1R,NR4A2,SH3BP2,PACSIN2,INPP5K,MYH9,IRS2,SDCBP,RASSF5,SH3BP5,TNIP1,SULF2,PTPN12,DOT1L,ATP6V1B2,CDC42BPG,CXCL16,PTEN,PELI1,MAP3K20,OSBPL8,RXRA,SEC14L1,SYAP1,MAP3K11,NIBAN2,MAPKAPK2,NUDT3,FRAT2,ZMIZ1,SLC44A2,SORT1,PIM3,NDRG2,STX3,TPT1,FOXO3,DDIT4,ZC3H3,MAP1LC3B,MAP3K8,SEMA4A,PXN,RHOQ,GABARAPL1,PINK1,SERINC3,UBE2D1,CPEB4,GABARAP,IQGAP1,BLOC1S6,ZFP36L1,RIOK3,ADIPOR1,PPARD,EIF4EBP2,MTMR3,GLUL,DNM2,PRKCD,USF2,STK38L,TFDP1,GNAQ,SGK1,WBP2,WDR45,RAB7A,DNAJC5,GNB2,EEF1D,MAPK1,KLHL24,PPP4R1,CHP1,CDK16,PRKACA,LAMP1 |
| GO:0051179 | BP | localization | 7.21663E-06 | 5.141665586 | 162 | KLF15,CD207,TULP2,RGPD2,PDGFRB,CXCL8,BEST1,PADI2,DGAT2,THBD,ADM,HBEGF,ATP8,STATH,KLF7,DSC2,ICAM3,DUSP1,FTH1,IL1R1,ZFP36,EGR3,ZSWIM4,C5AR2,ANPEP,CASS4,TRIB1,PYGL,SCN2A,ABHD5,TSC22D3,NEDD9,C5AR1,SOD2,DENND5A,ARRB2,OXSR1,ATG9B,SLC25A37,NR4A1,LAMB3,IMPDH1,TCN1,BCL6,IL6R,ATP2B1,CD44,PPIF,MAP2K3,GAB2,NDEL1,REPS2,TNFRSF10D,NFKBIA,ST3GAL4,LRG1,TP53INP2,CEBPB,SLC45A4,PNPLA8,CD302,MYADM,PLK3,SLC12A6,PID1,RABGEF1,PFKFB2,FAM53C,PPP1R15A,STAT5B,FOXO4,CEP68,NIN,MAN2B1,BICD2,SPAG9,MAP4K4,PER1,HIP1R,NR4A2,PACSIN2,PICALM,INPP5K,MYH9,IRS2,UBAP1,SSH2,SDCBP,RASSF5,RAB31,TMCO3,EIF2D,OSBPL1A,ATP6V1B2,CXCL16,PTEN,ERGIC1,ACTN1,OSBPL8,RXRA,SEC14L1,NIBAN2,STEAP4,MAPKAPK2,ZMIZ1,TUBA4A,SLC44A2,SORT1,LYST,PIM3,STX3,TPT1,NUP98,FOXO3,DDIT4,ZC3H3,VPS37B,TBC1D14,EEF2,ASAH1,SEMA4A,PXN,ARFGAP3,RHOQ,RAB3D,PINK1,UBE2D1,KLHL21,GABARAP,IQGAP1,BLOC1S6,ZFP36L1,ADIPOR1,PPARD,SLC25A40,SNX18,WDR1,RANBP2,GLUL,DNM2,PRKCD,USF2,DOP1B,TFDP1,ZSWIM6,YPEL5,CAMK1D,SGK1,WBP2,ANXA11,WDR45,CASC3,RAB7A,DNAJC5,B4GALT1,MAPK1,KLHL24,CHP1,CDK16,OSBPL2,PRKACA,LAMP1 |
| GO:0005829 | CC | cytosol | 9.83E-13 | 12.00737015 | 154 | SPRR3,BEST1,KRT13,PADI2,DGAT2,FOS,KLF7,FTH1,NAMPT,AMPD2,ARHGEF40,ZFP36,BTG2,IER3,TRIB1,PLEKHG2,PFKFB3,PYGL,ABHD5,TSC22D3,NEDD9,GYS1,NDRG1,ABTB1,DENND5A,ARRB2,OXSR1,UPP1,NR4A1,GPCPD1,IVNS1ABP,IMPDH1,CD44,PHLDA1,MAP2K3,GAB2,NDEL1,JOSD1,REPS2,NFKBIA,PI3,GSTO2,TP53INP2,USP32,MAPK13,FGD4,CNOT11,PRKDC,RALGDS,RABGEF1,RASA2,PFKFB2,PPP1R15A,STAT5B,FOXO4,CEP68,BICD2,SPAG9,NIBAN1,PER1,CDC42EP3,ETS2,HIP1R,PACSIN2,PICALM,INPP5K,MYH9,EIF3L,IRS2,KDM4B,UBAP1,SDCBP,TPM4,RAB31,KLHL2,TNIP1,EIF2D,OSBPL1A,PTPN12,ATP6V1B2,CDC42BPG,PTEN,PELI1,MAP3K20,ACTN1,OSBPL8,RXRA,SEC14L1,SYAP1,NIBAN2,MAPKAPK2,NUDT3,FRAT2,ERF,ZMIZ1,TUBA4A,SORT1,LYST,PIM3,NDRG2,TPT1,NUP98,FOXO3,DDIT4,TBC1D14,TFE3,MAP1LC3B,EEF2,MAP3K8,PXN,ARFGAP3,RNF144B,UBE2R2,RHOQ,GABARAPL1,PINK1,PABPC1,UBE2D1,KLHL21,IDI1,GABARAP,IQGAP1,BLOC1S6,ZFP36L1,RIOK3,WDR1,TKT,MTMR3,RANBP2,GLUL,DNM2,PRKCD,DOP1B,STK38L,TFDP1,SGK1,UBE2H,CMIP,ANXA11,WDR45,CASC3,RAB7A,MLLT1,DNAJC5,GNB2,EEF1D,MAPK1,CHP1,CDK16,FDPS,OSBPL2,PRKACA,LAMP1,DNAJB1 |
| GO:0005622 | CC | intracellular anatomical structure | 0.00246019 | 2.609031391 | 270 | TGM3,IL1R2,KLF15,FOSB,CD207,TULP2,RGPD2,SPRR3,PDGFRB,MUC7,BEST1,CAMK1G,SRSF12,KRT13,KLF2,PADI2,DGAT2,THBD,FOS,ADM,MUC21,HBEGF,DCUN1D3,ATP8,KLF7,DSC2,DUSP1,FTH1,PNRC1,NAMPT,EGR1,AMPD2,ARHGEF40,ZFP36,EGR3,BTG2,TMEM71,FOSL2,IER3,ANPEP,ZNF219,CASS4,NCCRP1,TRIB1,PLEKHG2,GK3P,PFKFB3,PYGL,LITAF,ABHD5,TSC22D3,NEDD9,ZNF442,C5AR1,JUNB,CTDSP1,GPR85,GYS1,NDRG1,SOD2,ABTB1,DENND5A,ARRB2,OXSR1,ATG9B,SLC25A37,UPP1,NR4A1,GPCPD1,IVNS1ABP,IMPDH1,TCN1,PLEKHG6,BCL6,ATP2B1,CD44,KLLN,PHLDA1,COL6A3,TLE3,PPIF,MAP2K3,GAB2,NDEL1,OPA3,JOSD1,REPS2,NFKBIA,PI3,GSTO2,ST3GAL4,LRG1,TP53INP2,CEBPB,USP32,KATNBL1,PNPLA8,MAPK13,FGD4,CD302,MYADM,PLK3,SHKBP1,CNOT11,PRKDC,MBOAT2,PID1,RALGDS,RABGEF1,RASA2,PFKFB2,FAM53C,MAML3,PPP1R15A,STAT5B,LFNG,FOXO4,CEP68,NIN,MAN2B1,BICD2,SPAG9,NIBAN1,MAP4K4,MTURN,PER1,CDC42EP3,ETS2,HIP1R,NR4A2,MAEA,PACSIN2,PICALM,INPP5K,MYH9,EIF3L,IRS2,KDM4B,UBAP1,NBPF9,SSH2,ABHD12B,SDCBP,RASSF5,TPD52L2,TPM4,SH3BP5,RAB31,KLHL2,TNIP1,SULF2,EIF2D,MORC4,OSBPL1A,PTPN12,DOT1L,ATP6V1B2,CDC42BPG,PTEN,PELI1,MAP3K20,ERGIC1,ACTN1,OSBPL8,RXRA,SEC14L1,SYAP1,MAP3K11,NIBAN2,STEAP4,CCNG2,MAPKAPK2,CCNQ,NUDT3,FRAT2,ERF,ZMIZ1,TUBA4A,SLC44A2,SORT1,LYST,PHF20L1,PIM3,NDRG2,STX3,ELL2,TPT1,R3HDM4,NUP98,FOXO3,DDIT4,ZC3H3,VPS37B,TBC1D14,TFE3,MAP1LC3B,FRMD4B,EEF2,ASAH1,MAP3K8,PXN,ARFGAP3,RNF144B,UBE2R2,RHOQ,IRF2BP2,GABARAPL1,RAB3D,PINK1,SERINC3,PABPC1,ZNF185,UBE2D1,CPEB4,KLHL21,IDI1,GABARAP,IQGAP1,BLOC1S6,ZFP36L1,RIOK3,PPARD,SLC25A40,YPEL3,SNX18,WDR1,MSL1,TKT,EIF4EBP2,MTMR3,EIF1,RANBP2,GLUL,DNM2,PRKCD,USF2,DOP1B,STK38L,TFDP1,YPEL5,CAMK1D,GNAQ,SGK1,UBE2H,RNF10,CMIP,WBP2,ANXA11,RBM47,KAT8,WDR45,CASC3,RAB7A,MLLT1,DNAJC5,B4GALT1,BZW1,RYBP,GNB2,EEF1D,MAPK1,KLHL24,CHP1,CDK16,FDPS,OSBPL2,PRKACA,LAMP1,DNAJB1,CCNI |
| GO:0005515 | MF | protein binding | 0.00811341 | 2.090796575 | 269 | SCGB3A1,IL1R2,KLF15,FOSB,CD207,RGPD2,SPRR3,PDGFRB,CXCL8,MUC7,BEST1,CAMK1G,SRSF12,KRT13,KLF2,PADI2,DGAT2,THBD,FOS,ADM,HBEGF,DCUN1D3,STATH,BTBD19,DSC2,ICAM3,DUSP1,LOXHD1,FTH1,PNRC1,NAMPT,IL1R1,EGR1,TREML2,ADRB3,ALPL,AMPD2,FCER1A,ZFP36,BTG2,TMEM71,C5AR2,FOSL2,IER3,ZNF219,CASS4,NCCRP1,TRIB1,PLEKHG2,PFKFB3,LYPD3,PYGL,SCN2A,A2ML1,LITAF,ABHD5,TSC22D3,KREMEN1,NEDD9,C5AR1,JUNB,CTDSP1,GPR85,GYS1,NDRG1,SOD2,ABTB1,DENND5A,ARRB2,OXSR1,SLC25A37,UPP1,NR4A1,LAMB3,IVNS1ABP,IMPDH1,METRNL,TCN1,PLEKHG6,BCL6,IL6R,ATP2B1,CD44,PHLDA1,COL6A3,TLE3,PPIF,MAP2K3,GAB2,NDEL1,JOSD1,REPS2,TNFRSF10D,NFKBIA,GSTO2,LRG1,TP53INP2,CEBPB,USP32,KATNBL1,MAPK13,FGD4,CD302,INPP5A,MYADM,GPX3,PLK3,SHKBP1,SLC12A6,CNOT11,PRKDC,PID1,RALGDS,RABGEF1,PFKFB2,FAM53C,PPP1R15A,STAT5B,FOXO4,CEP68,NIN,BICD2,SPAG9,NIBAN1,MAP4K4,MTURN,PER1,CDC42EP3,ETS2,MANSC1,HIP1R,NR4A2,MAEA,SH3BP2,PACSIN2,PICALM,INPP5K,MYH9,EIF3L,IRS2,UBAP1,SSH2,ABHD12B,SDCBP,RASSF5,TPD52L2,TPM4,SH3BP5,RAB31,KLHL2,TNIP1,MORC4,OSBPL1A,PTPN12,DOT1L,C16ORF72,ATP6V1B2,CDC42BPG,CXCL16,PTEN,PELI1,MAP3K20,ERGIC1,ACTN1,OSBPL8,NATD1,RXRA,SEC14L1,SYAP1,MAP3K11,NIBAN2,MAPKAPK2,CCNQ,NUDT3,ERF,ZMIZ1,TUBA4A,SORT1,LYST,PHF20L1,PIM3,NDRG2,STX3,ELL2,TPT1,NUP98,FOXO3,DDIT4,ZC3H3,VPS37B,TBC1D14,TFE3,MAP1LC3B,EEF2,ASAH1,MAP3K8,SEMA4A,PXN,ARFGAP3,RNF144B,UBE2R2,RHOQ,GABARAPL1,RAB3D,PINK1,SERINC3,PABPC1,ZNF185,UBE2D1,CPEB4,KLHL21,IDI1,GABARAP,IQGAP1,BLOC1S6,ZFP36L1,RIOK3,ADIPOR1,PPARD,SNX18,WDR1,MSL1,TKT,EIF4EBP2,MTMR3,EIF1,RANBP2,GLUL,DNM2,PRKCD,USF2,DOP1B,STK38L,TFDP1,YPEL5,CAMK1D,GNAQ,SGK1,UBE2H,RNF10,CMIP,WBP2,ANXA11,RBM47,KAT8,RNF141,WDR45,CASC3,RAB7A,MLLT1,DNAJC5,B4GALT1,ZCCHC14,BZW1,RYBP,GNB2,EEF1D,MAPK1,KLHL24,PPP4R1,CHP1,CDK16,FDPS,OSBPL2,PRKACA,LAMP1,DNAJB1,CCNI,RNASEK |
| GO:0043167 | MF | ion binding | 0.025064963 | 1.600932924 | 127 | TGM3,KLF15,PDGFRB,CAMK1G,KLF2,PADI2,THBD,KLF7,DSC2,FTH1,EGR1,ALPL,AMPD2,ZFP36,EGR3,ZSWIM4,ANPEP,ZNF219,TRIB1,GK3P,PFKFB3,PYGL,LITAF,ZNF442,CTDSP1,SOD2,OXSR1,NR4A1,IMPDH1,BCL6,ATP2B1,MAP2K3,GAB2,REPS2,USP32,PNPLA8,MAPK13,FGD4,PLK3,CPD,PRKDC,RABGEF1,RASA2,PFKFB2,LFNG,NIN,MAN2B1,MAP4K4,HIP1R,NR4A2,MAEA,PACSIN2,PICALM,MYH9,KDM4B,PCDHB1,SDCBP,RASSF5,TPM4,RAB31,SULF2,MORC4,ATP6V1B2,CDC42BPG,MAP3K20,ACTN1,RXRA,MAP3K11,STEAP4,MAPKAPK2,NUDT3,ZMIZ1,TUBA4A,PHF20L1,PIM3,STX3,TPT1,ZC3H3,EEF2,MAP3K8,PXN,ARFGAP3,RNF144B,UBE2R2,RHOQ,IRF2BP2,RAB3D,PINK1,ZNF185,UBE2D1,CPEB4,IDI1,IQGAP1,ZFP36L1,RIOK3,ADIPOR1,PPARD,YPEL3,SNX18,TKT,MTMR3,RANBP2,GLUL,DNM2,PRKCD,STK38L,ZSWIM6,YPEL5,CAMK1D,GNAQ,SGK1,UBE2H,RNF10,ANXA11,KAT8,RNF141,WDR45,RAB7A,B4GALT1,ZCCHC14,RYBP,MAPK1,CHP1,CDK16,FDPS,OSBPL2,PRKACA |

**Table 3** The enriched pathway terms of the up and down regulated differentially expressed genes

| **Pathway ID** | **Pathway Name** | | **Adjusted p value** | **Negative log10 of adjusted p value** | **Gene Count** | **Gene** |
| --- | --- | --- | --- | --- | --- | --- |
| **Up regulated genes** | | | | | | |
| REAC:R-HSA-168256 | Immune System | | 5.34E-25 | 24.27238886 | 127 | CXCL10,ISG15,IFI6,SIGLEC1,IFIT3,IFIT1,BST2,IFI27,RSAD2,OASL,SERPING1,IFIT2,USP18,LAG3,HERC6,OAS3,IFITM1,IFITM3,OAS2,OAS1,CCL2,CD80,DDX58,HERC5,C1QA,MX1,IFIT5,IFNG,H3C7,IFIH1,TICAM2,H3C12,MUC13,C1QC,STAT1,KLRC2,CCR5,UBE2L6,BTN3A3,SAA1,C4B,PDCD1LG2,IFI35,HLA-DOB,MT2A,H3C2,PSMB9,FASLG,DHX58,HLA-DMB,BTN3A1,CD3D,CD3E,GBP1,GRAP2,XAF1,PRLR,TRIM22,H3C3,PDCD1,ISG20,IRF7,HLA-F,DEFB1,CD8A,H3C8,HLA-G,CD8B,H3C10,IRF4,C4A,EIF2AK2,SH2D1A,ZAP70,KIF15,HLA-DRB5,KLRC1,PSME2,PML,CENPE,HLA-DMA,TAP1,CTLA4,TLR3,GBP5,NOS2,HLA-DRB1,AIM2,CFB,TLR7,TNFSF13B,DTX3L,IRF8,SLAMF7,CCR1,IL7,ZBP1,FCGR3A,BPIFA1,CD300E,PSMB8,TAP2,MX2,GBP4,CD40LG,CD226,TNFRSF13B,STAT2,CFH,WASF1,CRTAM,GNLY,RNASE6,CARD11,H3C13,IL15RA,FYN,IL32,TUBA8,NLRC5,FBXO6,STAT4,CD96,IRS1,CD33,LGALS9,CD74 |
| REAC:R-HSA-3214815 | HDACs deacetylate histones | | 1.06E-19 | 18.97494292 | 27 | H2AC16,H3C7,H2AC19,H3C12,H2AC14,H2BC11,H3C2,H2BC14,H2BC17,H2AC12,H4C13,H2BC3,H4C14,H3C3,H3C8,H3C10,H2AC17,H2AC13,H2BC7,H4C11,H2BC10,H4C6,H2BC13,H3C13,H2BC6,H2AC7,H2AC20 |
| REAC:R-HSA-5689880 | Ub-specific processing proteases | | 2.76E-07 | 6.558680932 | 22 | USP18,DDX58,H2AC16,H2AC19,IFIH1,H2AC14,H2BC11,H2BC14,H2BC17,H2AC12,PSMB9,H2BC3,H2AC17,PSME2,H2AC13,H2BC7,PSMB8,H2BC10,H2BC13,H2BC6,H2AC7,H2AC20 |
| REAC:R-HSA-1280218 | Adaptive Immune System | | 6.37E-08 | 7.195827239 | 48 | SIGLEC1,LAG3,HERC6,IFITM1,CD80,HERC5,UBE2L6,BTN3A3,PDCD1LG2,HLA-DOB,PSMB9,HLA-DMB,BTN3A1,CD3D,CD3E,GRAP2,PDCD1,HLA-F,CD8A,HLA-G,CD8B,SH2D1A,ZAP70,KIF15,HLA-DRB5,KLRC1,PSME2,CENPE,HLA-DMA,TAP1,CTLA4,HLA-DRB1,DTX3L,SLAMF7,FCGR3A,CD300E,PSMB8,TAP2,CD40LG,CD226,CRTAM,CARD11,FYN,TUBA8,FBXO6,CD96,CD33,CD74 |
| REAC:R-HSA-5663205 | Infectious disease | | 6.85E-08 | 7.164097591 | 52 | ISG15,ACE2,H2AC16,H3C7,H2AC19,H3C12,PARP9,H2AC14,CCR5,PARP14,H2BC11,ADCY5,H3C2,H2BC14,H2BC17,H2AC12,PSMB9,H4C13,H2BC3,H4C14,H3C3,PDCD1,H3C8,CD8B,GGT1,H3C10,EIF2AK2,H2AC17,PSME2,PML,H2AC13,NOS2,H2BC7,TLR7,ZBP1,FCGR3A,PSMB8,H4C11,H2BC10,H4C6,STAT2,WASF1,H2BC13,GNGT2,H3C13,FXYD6,FYN,TUBA8,H2BC6,H2AC7,RAMP1,H2AC20 |
| REAC:R-HSA-198933 | Immunoregulatory interactions between a Lymphoid and a non-Lymphoid cell | | 4.88047E-06 | 5.311538043 | 18 | SIGLEC1,IFITM1,CD3D,CD3E,HLA-F,CD8A,HLA-G,CD8B,SH2D1A,KLRC1,SLAMF7,FCGR3A,CD300E,CD40LG,CD226,CRTAM,CD96,CD33 |
| **Down regulated genes** | | | | | | |
| REAC:R-HSA-187037 | | Signaling by NTRK1 (TRKA) | 0.000139886 | 3.854225447 | 13 | FOSB,FOS,EGR1,EGR3,TRIB1,JUNB,MAPK13,RALGDS,IRS2,MAPKAPK2,DNM2,SGK1,MAPK1 |
| REAC:R-HSA-168256 | | Immune System | 0.000483116 | 3.315948984 | 68 | IL1R2,CD207,CXCL8,MUC7,PADI2,FOS,MUC21,ICAM3,FTH1,IL1R1,EGR1,TREML2,FCER1A,C5AR2,ANPEP,PYGL,C5AR1,JUNB,SOD2,IMPDH1,TCN1,BCL6,IL6R,CD44,MAP2K3,GAB2,NFKBIA,PI3,LRG1,MAPK13,PRKDC,STAT5B,MAN2B1,MYH9,IRS2,SDCBP,RAB31,KLHL2,OSBPL1A,PTPN12,ATP6V1B2,PTEN,PELI1,MAPKAPK2,TUBA4A,SLC44A2,STX3,FOXO3,EEF2,ASAH1,MAP3K8,RNF144B,UBE2R2,RAB3D,UBE2D1,KLHL21,IQGAP1,RANBP2,DNM2,PRKCD,YPEL5,UBE2H,RAB7A,DNAJC5,B4GALT1,MAPK1,PRKACA,LAMP1 |
| REAC:R-HSA-168249 | | Innate Immune System | 0.000483116 | 3.315948984 | 44 | MUC7,PADI2,FOS,MUC21,ICAM3,FTH1,FCER1A,C5AR2,ANPEP,PYGL,C5AR1,IMPDH1,TCN1,CD44,MAP2K3,GAB2,NFKBIA,PI3,LRG1,MAPK13,PRKDC,MAN2B1,MYH9,SDCBP,RAB31,ATP6V1B2,PELI1,MAPKAPK2,SLC44A2,EEF2,ASAH1,MAP3K8,RAB3D,UBE2D1,IQGAP1,DNM2,PRKCD,YPEL5,RAB7A,DNAJC5,B4GALT1,MAPK1,PRKACA,LAMP1 |
| REAC:R-HSA-6798695 | | Neutrophil degranulation | 0.003082493 | 2.511097929 | 24 | PADI2,FTH1,ANPEP,PYGL,C5AR1,IMPDH1,TCN1,CD44,LRG1,MAN2B1,SDCBP,RAB31,SLC44A2,EEF2,ASAH1,RAB3D,IQGAP1,PRKCD,YPEL5,RAB7A,DNAJC5,B4GALT1,MAPK1,LAMP1 |
| REAC:R-HSA-9006934 | | Signaling by Receptor Tyrosine Kinases | 0.003779389 | 2.422578418 | 24 | FOSB,PDGFRB,FOS,HBEGF,EGR1,EGR3,TRIB1,JUNB,LAMB3,COL6A3,GAB2,MAPK13,RALGDS,STAT5B,IRS2,PTPN12,ATP6V1B2,MAPKAPK2,PXN,DNM2,PRKCD,SGK1,MAPK1,PRKACA |
| REAC:R-HSA-199991 | | Membrane Trafficking | 0.021636488 | 1.664813234 | 26 | HBEGF,FTH1,DENND5A,ARRB2,REPS2,CPD,RABGEF1,BICD2,HIP1R,PACSIN2,PICALM,MYH9,UBAP1,RAB31,TUBA4A,SORT1,VPS37B,TBC1D14,MAP1LC3B,ARFGAP3,RHOQ,GABARAP,BLOC1S6,SNX18,DNM2,RAB7A |

**Table 4** Topology table for up and down regulated genes.

| **Category** | **Node** | **Degree** | **Betweenness** | **Stress** | **Closeness** |
| --- | --- | --- | --- | --- | --- |
| Up | ESR1 | 798 | 0.192517 | 1.77E+08 | 0.408682 |
| Up | UBD | 502 | 0.118413 | 96176834 | 0.369243 |
| Up | FYN | 273 | 0.061939 | 30015938 | 0.359012 |
| Up | STAT1 | 223 | 0.05064 | 18096366 | 0.397044 |
| Up | ISG15 | 188 | 0.027621 | 17693852 | 0.366029 |
| Up | NOS2 | 170 | 0.023391 | 17364304 | 0.360678 |
| Up | PML | 163 | 0.02224 | 15000134 | 0.36002 |
| Up | FBXO6 | 153 | 0.029751 | 15338582 | 0.345429 |
| Up | PTN | 108 | 0.019461 | 7122098 | 0.338033 |
| Up | FASLG | 83 | 0.014694 | 4374476 | 0.35841 |
| Up | EIF2AK2 | 81 | 0.010682 | 5028286 | 0.349214 |
| Up | IRF8 | 73 | 0.009166 | 3207394 | 0.351752 |
| Up | TGM2 | 69 | 0.008347 | 3649916 | 0.345615 |
| Up | IRF7 | 66 | 0.009403 | 3212578 | 0.348905 |
| Up | IRF4 | 60 | 0.007151 | 2266524 | 0.347959 |
| Up | IFIT3 | 60 | 0.003302 | 1795640 | 0.330435 |
| Up | DDX58 | 56 | 0.006449 | 2057946 | 0.346408 |
| Up | IRS1 | 56 | 0.003937 | 1608812 | 0.357287 |
| Up | BUB1B | 53 | 0.005921 | 2158524 | 0.336724 |
| Up | ZAP70 | 53 | 0.004798 | 1904462 | 0.34328 |
| Up | IFIT2 | 49 | 0.002087 | 1246440 | 0.324009 |
| Up | IFIT1 | 49 | 0.004345 | 1582522 | 0.334224 |
| Up | NMI | 47 | 0.007103 | 1727992 | 0.345941 |
| Up | STAT2 | 45 | 0.003261 | 1208362 | 0.349523 |
| Up | GFI1 | 45 | 0.005993 | 2163282 | 0.337699 |
| Up | PSMB8 | 42 | 0.00213 | 1084100 | 0.336879 |
| Up | UBE2L6 | 40 | 0.003621 | 1208660 | 0.334354 |
| Up | GZMB | 38 | 0.004228 | 1177768 | 0.321915 |
| Up | PSME2 | 37 | 0.002791 | 1085450 | 0.335272 |
| Up | MYBL2 | 36 | 0.003331 | 1073716 | 0.320627 |
| Up | LGALS3BP | 36 | 0.002906 | 1015894 | 0.35396 |
| Up | BUB1 | 34 | 0.003714 | 1122524 | 0.33668 |
| Up | TAP1 | 33 | 0.003564 | 1048476 | 0.340862 |
| Up | TPX2 | 32 | 0.003285 | 1158360 | 0.335272 |
| Up | CD247 | 32 | 0.002306 | 611176 | 0.343303 |
| Up | CCR5 | 31 | 0.004564 | 1252458 | 0.323988 |
| Up | GRAP2 | 30 | 0.003826 | 1039970 | 0.336018 |
| Up | PTPRO | 29 | 0.002308 | 639118 | 0.335162 |
| Up | CCL2 | 29 | 0.004485 | 1056792 | 0.320808 |
| Up | CD3E | 28 | 0.002918 | 706426 | 0.335425 |
| Up | ABCB1 | 28 | 0.002424 | 759344 | 0.342523 |
| Up | CDC7 | 28 | 0.002973 | 878770 | 0.335097 |
| Up | SH2D1A | 27 | 0.004057 | 742976 | 0.343855 |
| Up | BATF3 | 27 | 0.001939 | 569966 | 0.294124 |
| Up | TLR3 | 25 | 0.002148 | 559984 | 0.314863 |
| Up | PSAT1 | 24 | 0.002325 | 756588 | 0.334376 |
| Up | TNFSF10 | 24 | 0.00138 | 426574 | 0.312828 |
| Up | MTNR1A | 24 | 0.001564 | 1286488 | 0.287886 |
| Up | IFNG | 24 | 0.001178 | 365080 | 0.321189 |
| Up | PSMB9 | 24 | 0.001442 | 381174 | 0.352476 |
| Up | CARD11 | 23 | 0.001688 | 663288 | 0.333767 |
| Up | PRLR | 23 | 0.001176 | 469112 | 0.339172 |
| Up | C1QA | 23 | 0.002946 | 1093304 | 0.297122 |
| Up | WASF1 | 23 | 0.002571 | 472204 | 0.337388 |
| Up | IFIH1 | 22 | 0.001263 | 491138 | 0.335754 |
| Up | HERC5 | 22 | 0.002237 | 725500 | 0.354743 |
| Up | DTX3L | 22 | 0.001709 | 565192 | 0.334812 |
| Up | GZMA | 21 | 0.001501 | 1112872 | 0.304348 |
| Up | MT2A | 21 | 0.003383 | 615218 | 0.339352 |
| Up | CENPE | 20 | 0.001417 | 401202 | 0.339374 |
| Up | BCL11B | 20 | 0.001121 | 474760 | 0.334115 |
| Up | DOK2 | 20 | 0.00153 | 524938 | 0.333095 |
| Up | STAT4 | 20 | 7.57E-04 | 243488 | 0.309078 |
| Up | IFIT5 | 20 | 0.001119 | 410524 | 0.339217 |
| Up | STAMBPL1 | 18 | 0.001077 | 398602 | 0.332707 |
| Up | LAP3 | 18 | 0.001222 | 391612 | 0.331439 |
| Up | UBE2T | 18 | 6.27E-04 | 352842 | 0.334093 |
| Up | C7orf25 | 18 | 0.002466 | 690114 | 0.30653 |
| Up | KIF5C | 18 | 0.00143 | 447542 | 0.346642 |
| Up | CD8A | 18 | 0.001532 | 323014 | 0.332685 |
| Up | HLA-DRB1 | 18 | 0.001445 | 467308 | 0.333463 |
| Up | GBP1 | 18 | 3.90E-04 | 148994 | 0.324685 |
| Up | CD2 | 18 | 0.001782 | 349704 | 0.339172 |
| Up | LGALS9 | 17 | 0.00215 | 355432 | 0.305981 |
| Up | DHX58 | 17 | 0.001152 | 332654 | 0.309134 |
| Up | CXCL10 | 17 | 0.001168 | 350182 | 0.312314 |
| Up | PROX1 | 17 | 0.001217 | 289316 | 0.308465 |
| Up | OAS3 | 16 | 0.001148 | 310514 | 0.332836 |
| Up | MX1 | 16 | 0.002524 | 376922 | 0.336901 |
| Up | RYR1 | 16 | 0.002311 | 432186 | 0.331824 |
| Up | PLK4 | 15 | 0.001828 | 415966 | 0.33262 |
| Up | TAP2 | 15 | 9.76E-04 | 246018 | 0.335184 |
| Up | RNF43 | 15 | 0.001445 | 297884 | 0.337632 |
| Up | ABI3 | 15 | 0.00213 | 1231380 | 0.256422 |
| Up | CFH | 15 | 0.0012 | 263858 | 0.317709 |
| Up | KCNA2 | 15 | 6.23E-04 | 141946 | 0.289282 |
| Up | CYP19A1 | 14 | 3.07E-04 | 115996 | 0.303789 |
| Up | DPP4 | 14 | 0.002365 | 447304 | 0.277982 |
| Up | CCR1 | 14 | 0.001606 | 603916 | 0.246837 |
| Up | SP110 | 14 | 0.001349 | 851082 | 0.278148 |
| Up | NUSAP1 | 14 | 0.001144 | 264292 | 0.334463 |
| Up | CD40LG | 14 | 0.001271 | 269880 | 0.344247 |
| Up | FXYD6 | 14 | 0.002095 | 420154 | 0.332125 |
| Up | SERPING1 | 13 | 0.001794 | 274630 | 0.293771 |
| Up | CD33 | 13 | 6.01E-04 | 159724 | 0.332858 |
| Up | ZBP1 | 13 | 0.001638 | 1766968 | 0.269996 |
| Up | WDR74 | 13 | 0.001739 | 291056 | 0.338166 |
| Up | CTLA4 | 13 | 6.33E-04 | 132938 | 0.312485 |
| Up | KIF15 | 13 | 0.001676 | 307736 | 0.345266 |
| Up | MDK | 13 | 8.84E-04 | 565792 | 0.278874 |
| Up | CD3D | 13 | 7.11E-04 | 205180 | 0.328847 |
| Up | HLA-G | 13 | 0.00164 | 260418 | 0.327838 |
| Up | CD74 | 12 | 0.001173 | 235748 | 0.335075 |
| Up | BACH2 | 12 | 3.79E-04 | 146352 | 0.336261 |
| Up | ASPM | 12 | 5.62E-04 | 216470 | 0.332642 |
| Up | SLC16A1 | 11 | 0.001648 | 262392 | 0.33355 |
| Up | UBASH3A | 11 | 4.97E-04 | 160866 | 0.331996 |
| Up | KCNA3 | 11 | 9.51E-04 | 181440 | 0.331781 |
| Up | SLC12A3 | 11 | 4.71E-04 | 159292 | 0.327901 |
| Up | MYO7A | 11 | 9.03E-04 | 219204 | 0.332512 |
| Up | CXCL11 | 11 | 5.54E-04 | 192692 | 0.311177 |
| Up | TBX21 | 11 | 6.87E-04 | 139852 | 0.31065 |
| Up | GRAP | 10 | 7.31E-05 | 39208 | 0.287967 |
| Up | IFITM1 | 10 | 9.96E-04 | 194092 | 0.333225 |
| Up | APOD | 10 | 0.001108 | 491794 | 0.306493 |
| Up | BST2 | 10 | 9.30E-04 | 185754 | 0.33131 |
| Up | HLA-DRB5 | 10 | 2.46E-04 | 129660 | 0.326774 |
| Up | USP18 | 10 | 8.64E-04 | 134798 | 0.336283 |
| Up | TNFRSF13B | 10 | 6.43E-04 | 297044 | 0.28414 |
| Up | C4A | 10 | 0.001095 | 531836 | 0.286041 |
| Up | CD38 | 4 | 6.28E-05 | 20058 | 0.284171 |
| Up | CCL8 | 4 | 8.97E-05 | 52842 | 0.271611 |
| Up | CD244 | 3 | 1.01E-06 | 282 | 0.266809 |
| Up | CXCL9 | 3 | 4.32E-04 | 118944 | 0.307043 |
| Up | IFITM3 | 3 | 2.30E-05 | 5666 | 0.285866 |
| Up | NLRC5 | 2 | 3.30E-07 | 254 | 0.258476 |
| Up | IDO1 | 2 | 0 | 0 | 0.284581 |
| Up | C1R | 2 | 5.02E-06 | 1022 | 0.238252 |
| Up | SIT1 | 2 | 0 | 0 | 0.265264 |
| Up | IFI27 | 2 | 0 | 0 | 0.284518 |
| Up | IFI35 | 2 | 1.50E-05 | 6072 | 0.284676 |
| Up | PRF1 | 2 | 2.61E-06 | 974 | 0.250049 |
| Up | C1QB | 2 | 2.70E-04 | 32182 | 0.277501 |
| Up | CD48 | 2 | 8.18E-06 | 2242 | 0.259246 |
| Up | IFI6 | 2 | 0 | 0 | 0.284518 |
| Up | SLAMF1 | 2 | 0 | 0 | 0.266642 |
| Up | TICAM2 | 2 | 4.90E-06 | 1370 | 0.258789 |
| Up | HELZ2 | 2 | 2.73E-05 | 21354 | 0.274253 |
| Up | SLAMF7 | 2 | 7.09E-05 | 15008 | 0.282125 |
| Up | GRIK1 | 1 | 0 | 0 | 0.250171 |
| Up | TNFSF13B | 1 | 0 | 0 | 0.221278 |
| Up | C1QC | 1 | 0 | 0 | 0.229073 |
| Up | CD226 | 1 | 0 | 0 | 0.264185 |
| Up | CD80 | 1 | 0 | 0 | 0.238097 |
| Up | SAA2 | 1 | 0 | 0 | 0.275239 |
| Up | GSDMB | 1 | 0 | 0 | 0.276798 |
| Up | STAP1 | 1 | 0 | 0 | 0.276798 |
| Up | CDCA2 | 1 | 0 | 0 | 0.276798 |
| Up | BCL2L14 | 1 | 0 | 0 | 0.276798 |
| Up | TFEC | 1 | 0 | 0 | 0.237943 |
| Up | CXCL12 | 1 | 0 | 0 | 0.217526 |
| Up | TLR7 | 1 | 0 | 0 | 0.262187 |
| Up | PARP12 | 1 | 0 | 0 | 0.262187 |
| Up | ACY3 | 1 | 0 | 0 | 0.250207 |
| Up | XAF1 | 1 | 0 | 0 | 0.284219 |
| Up | RSAD2 | 1 | 0 | 0 | 0.284219 |
| Up | GGT1 | 1 | 0 | 0 | 0.251655 |
| Up | TRO | 1 | 0 | 0 | 0.259653 |
| Up | RORB | 1 | 0 | 0 | 0.242555 |
| Up | PARP9 | 1 | 0 | 0 | 0.250843 |
| Up | GNGT2 | 1 | 0 | 0 | 0.261759 |
| Up | AIM2 | 1 | 0 | 0 | 0.249951 |
| Up | CRTAM | 1 | 0 | 0 | 0.276798 |
| Up | OAS1 | 1 | 0 | 0 | 0.259011 |
| Up | CD8B | 1 | 0 | 0 | 0.249647 |
| Up | IL32 | 1 | 0 | 0 | 0.264785 |
| Up | TMSB10 | 1 | 0 | 0 | 0.277952 |
| Up | SLFN5 | 1 | 0 | 0 | 0.260682 |
| Up | ENPP2 | 1 | 0 | 0 | 0.269684 |
| Up | PTPRCAP | 1 | 0 | 0 | 0.276798 |
| Up | BATF2 | 1 | 0 | 0 | 0.256717 |
| Up | TUBA8 | 1 | 0 | 0 | 0.26916 |
| Up | ETV7 | 1 | 0 | 0 | 0.258151 |
| Down | EGR1 | 518 | 0.137274 | 73984308 | 0.382711 |
| Down | ARRB2 | 311 | 0.0549 | 47924518 | 0.368262 |
| Down | UBE2D1 | 238 | 0.047029 | 25945120 | 0.357486 |
| Down | PRKDC | 220 | 0.03817 | 19097472 | 0.370202 |
| Down | FOS | 214 | 0.043308 | 15087474 | 0.384921 |
| Down | MAPK1 | 187 | 0.036123 | 10770246 | 0.3953 |
| Down | PXN | 181 | 0.029271 | 18299458 | 0.360121 |
| Down | CEBPB | 154 | 0.026379 | 9111334 | 0.379737 |
| Down | PABPC1 | 152 | 0.024666 | 9387102 | 0.377279 |
| Down | RXRA | 150 | 0.027 | 7639360 | 0.379063 |
| Down | NFKBIA | 142 | 0.018107 | 11480668 | 0.359994 |
| Down | BCL6 | 139 | 0.026812 | 10769616 | 0.355332 |
| Down | EEF2 | 118 | 0.015181 | 5494630 | 0.379259 |
| Down | MAP1LC3B | 116 | 0.019024 | 7093962 | 0.343878 |
| Down | MYH9 | 114 | 0.018836 | 6847402 | 0.372568 |
| Down | PRKACA | 111 | 0.022264 | 7520924 | 0.354057 |
| Down | NR4A1 | 107 | 0.020235 | 8432996 | 0.357611 |
| Down | EEF1D | 99 | 0.012436 | 6378050 | 0.342569 |
| Down | MAPK13 | 96 | 0.010247 | 6734144 | 0.347205 |
| Down | IQGAP1 | 92 | 0.011294 | 4572162 | 0.367787 |
| Down | PTEN | 92 | 0.017001 | 4610068 | 0.373762 |
| Down | NDRG1 | 88 | 0.009487 | 4980860 | 0.357038 |
| Down | GABARAP | 87 | 0.010537 | 3939104 | 0.338479 |
| Down | ACTN1 | 87 | 0.012382 | 4256206 | 0.359389 |
| Down | RANBP2 | 82 | 0.008093 | 4767628 | 0.348953 |
| Down | SGK1 | 82 | 0.01006 | 5455636 | 0.342729 |
| Down | GABARAPL1 | 81 | 0.009201 | 3841908 | 0.339083 |
| Down | DNM2 | 81 | 0.013143 | 4359556 | 0.340072 |
| Down | UBE2H | 76 | 0.009903 | 4234864 | 0.338323 |
| Down | PDGFRB | 74 | 0.008996 | 3335988 | 0.353643 |
| Down | FOXO3 | 74 | 0.008196 | 3288092 | 0.369376 |
| Down | TUBA4A | 71 | 0.006319 | 3097846 | 0.347959 |
| Down | PRKCD | 70 | 0.009284 | 3892990 | 0.350694 |
| Down | ATP6V1B2 | 65 | 0.008552 | 3330612 | 0.340184 |
| Down | STAT5B | 62 | 0.006782 | 2481222 | 0.344733 |
| Down | JUNB | 62 | 0.007011 | 2447096 | 0.345359 |
| Down | PTPN12 | 57 | 0.007343 | 2176734 | 0.343832 |
| Down | CD44 | 56 | 0.008336 | 2860602 | 0.360247 |
| Down | RNF10 | 56 | 0.006127 | 2367936 | 0.341816 |
| Down | SOD2 | 53 | 0.008034 | 2443178 | 0.337544 |
| Down | RNF144B | 53 | 0.005069 | 2017554 | 0.336724 |
| Down | DNAJB1 | 50 | 0.006088 | 2062626 | 0.339936 |
| Down | EIF3L | 48 | 0.00468 | 1955518 | 0.332125 |
| Down | NDEL1 | 47 | 0.007994 | 1584292 | 0.340591 |
| Down | GNB2 | 46 | 0.008922 | 1639312 | 0.354547 |
| Down | RAB7A | 46 | 0.006877 | 1677636 | 0.35396 |
| Down | ETS2 | 46 | 0.004714 | 1536566 | 0.353691 |
| Down | TPD52L2 | 46 | 0.006465 | 1995176 | 0.335469 |
| Down | PINK1 | 45 | 0.00464 | 1826106 | 0.345895 |
| Down | TNFRSF10D | 44 | 0.003308 | 1603192 | 0.337855 |
| Down | USF2 | 44 | 0.006294 | 2069964 | 0.335645 |
| Down | TFE3 | 44 | 0.003494 | 2757172 | 0.312219 |
| Down | KAT8 | 43 | 0.005464 | 1597380 | 0.323865 |
| Down | FTH1 | 43 | 0.004707 | 1493774 | 0.335689 |
| Down | TPM4 | 42 | 0.003909 | 1867058 | 0.339217 |
| Down | TKT | 41 | 0.003604 | 1344208 | 0.361848 |
| Down | RYBP | 39 | 0.005665 | 1677814 | 0.335272 |
| Down | GAB2 | 38 | 0.001218 | 711446 | 0.323111 |
| Down | TNIP1 | 38 | 0.004626 | 1381986 | 0.351438 |
| Down | MAP3K11 | 38 | 0.003908 | 1706650 | 0.336194 |
| Down | NUDT3 | 38 | 0.005959 | 1613400 | 0.330457 |
| Down | GNAQ | 37 | 0.007581 | 1632484 | 0.330138 |
| Down | IRS2 | 37 | 0.001379 | 715496 | 0.355949 |
| Down | NUP98 | 35 | 0.003469 | 1591646 | 0.334965 |
| Down | MAP3K8 | 35 | 0.001498 | 677496 | 0.33368 |
| Down | TFDP1 | 34 | 0.004349 | 1304508 | 0.334703 |
| Down | PPARD | 34 | 0.002647 | 976048 | 0.339509 |
| Down | NEDD9 | 33 | 0.002263 | 650036 | 0.344779 |
| Down | RASSF5 | 33 | 0.00421 | 853638 | 0.337788 |
| Down | PACSIN2 | 32 | 0.003634 | 812032 | 0.338501 |
| Down | MAPKAPK2 | 32 | 0.003363 | 726882 | 0.341953 |
| Down | TPT1 | 32 | 0.002106 | 917626 | 0.347888 |
| Down | WDR1 | 32 | 0.004819 | 1012580 | 0.360703 |
| Down | ZFP36 | 31 | 0.004454 | 1090138 | 0.350072 |
| Down | DUSP1 | 30 | 0.002677 | 870036 | 0.354939 |
| Down | NR4A2 | 30 | 0.003467 | 814860 | 0.311215 |
| Down | TSC22D3 | 29 | 0.002154 | 635518 | 0.360931 |
| Down | DOT1L | 29 | 0.002244 | 720134 | 0.322319 |
| Down | SPAG9 | 28 | 0.002956 | 613724 | 0.339554 |
| Down | MAP2K3 | 28 | 0.002387 | 736730 | 0.334638 |
| Down | SDCBP | 27 | 0.005582 | 1210604 | 0.333615 |
| Down | FOSL2 | 27 | 0.002273 | 705484 | 0.354498 |
| Down | PPP1R15A | 27 | 0.001703 | 1037016 | 0.311328 |
| Down | UBE2R2 | 27 | 0.003032 | 777410 | 0.334376 |
| Down | MLLT1 | 26 | 0.001525 | 390820 | 0.296213 |
| Down | PPIF | 26 | 0.003794 | 881822 | 0.333615 |
| Down | PICALM | 26 | 0.002516 | 757500 | 0.336569 |
| Down | MAP4K4 | 25 | 0.002758 | 917576 | 0.333117 |
| Down | TLE3 | 25 | 0.003223 | 628204 | 0.335403 |
| Down | SH3BP2 | 24 | 0.001216 | 410836 | 0.339666 |
| Down | BLOC1S6 | 24 | 0.004921 | 2205528 | 0.298297 |
| Down | IL1R1 | 24 | 0.003639 | 728952 | 0.344895 |
| Down | USP32 | 24 | 0.002234 | 766640 | 0.333095 |
| Down | LYST | 24 | 0.004414 | 1162826 | 0.32765 |
| Down | RALGDS | 23 | 0.003002 | 711328 | 0.301611 |
| Down | LAMP1 | 22 | 0.003428 | 700048 | 0.333225 |
| Down | FOXO4 | 22 | 0.001106 | 455348 | 0.356394 |
| Down | GYS1 | 22 | 0.003454 | 678486 | 0.349785 |
| Down | OXSR1 | 22 | 0.00287 | 500966 | 0.352573 |
| Down | PELI1 | 21 | 0.001214 | 477420 | 0.338613 |
| Down | GLUL | 21 | 0.003434 | 834082 | 0.357188 |
| Down | IVNS1ABP | 21 | 0.001213 | 471132 | 0.346105 |
| Down | PYGL | 20 | 0.00345 | 602684 | 0.332663 |
| Down | NAMPT | 20 | 0.002212 | 380268 | 0.336261 |
| Down | BTG2 | 20 | 0.002656 | 551622 | 0.332018 |
| Down | PNRC1 | 19 | 0.001899 | 443316 | 0.320207 |
| Down | CAMK1D | 19 | 0.002449 | 987922 | 0.296162 |
| Down | RAB31 | 19 | 0.002284 | 614060 | 0.327671 |
| Down | BZW1 | 19 | 0.00303 | 357544 | 0.352283 |
| Down | ELL2 | 18 | 0.00274 | 321386 | 0.338859 |
| Down | INPP5K | 18 | 0.002393 | 526368 | 0.333117 |
| Down | CDK16 | 18 | 5.60E-04 | 330784 | 0.334725 |
| Down | RIOK3 | 17 | 0.001547 | 464048 | 0.327713 |
| Down | FOSB | 17 | 8.83E-05 | 66880 | 0.296864 |
| Down | PLK3 | 16 | 0.001804 | 408658 | 0.333702 |
| Down | CCNG2 | 16 | 8.68E-04 | 216670 | 0.354229 |
| Down | ANXA11 | 16 | 0.001671 | 301984 | 0.331889 |
| Down | FDPS | 15 | 7.11E-04 | 275776 | 0.332642 |
| Down | ERGIC1 | 15 | 0.004069 | 571924 | 0.325881 |
| Down | SH3BP5 | 15 | 8.56E-04 | 284762 | 0.327566 |
| Down | EGR3 | 3 | 3.96E-06 | 1002 | 0.283465 |
| Down | IER3 | 3 | 1.70E-05 | 7456 | 0.289429 |
| Down | HBEGF | 2 | 2.31E-05 | 7130 | 0.274576 |
| Down | ASAH1 | 2 | 3.49E-05 | 12710 | 0.257503 |
| Down | IDS | 2 | 6.65E-05 | 88902 | 0.300199 |
| Down | CAMK1G | 2 | 1.73E-04 | 159302 | 0.292448 |
| Down | KDM4B | 1 | 0 | 0 | 0.290133 |
| Down | NDRG2 | 1 | 0 | 0 | 0.290133 |
| Down | NIN | 1 | 0 | 0 | 0.261492 |
| Down | KLF2 | 1 | 0 | 0 | 0.276798 |
| Down | PHLDA1 | 1 | 0 | 0 | 0.253708 |
| Down | C5AR1 | 1 | 0 | 0 | 0.26916 |
| Down | OSBPL1A | 1 | 0 | 0 | 0.262187 |
| Down | SNX18 | 1 | 0 | 0 | 0.263859 |
| Down | SERINC3 | 1 | 0 | 0 | 0.250134 |
| Down | HIP1R | 1 | 0 | 0 | 0.251828 |
| Down | PHF20L1 | 1 | 0 | 0 | 0.244648 |
| Down | IFRD1 | 1 | 0 | 0 | 0.258841 |
| Down | ANPEP | 1 | 0 | 0 | 0.261292 |
| Down | B4GALT1 | 1 | 0 | 0 | 0.259312 |
| Down | ZFP36L1 | 1 | 0 | 0 | 0.25483 |
| Down | TGM3 | 1 | 0 | 0 | 0.261292 |
| Down | UPP1 | 1 | 0 | 0 | 0.263358 |
| Down | PFKFB3 | 1 | 0 | 0 | 0.26916 |
| Down | MSL1 | 1 | 0 | 0 | 0.244648 |
| Down | NCCRP1 | 1 | 0 | 0 | 0.290133 |
| Down | MAEA | 1 | 0 | 0 | 0.269684 |
| Down | PID1 | 1 | 0 | 0 | 0.276798 |
| Down | TP53INP2 | 1 | 0 | 0 | 0.252896 |
| Down | BICD2 | 1 | 0 | 0 | 0.258697 |
| Down | SEC14L1 | 1 | 0 | 0 | 0.257296 |
| Down | MYADM | 1 | 0 | 0 | 0.276798 |
| Down | OSBPL8 | 1 | 0 | 0 | 0.269684 |
| Down | PLEKHG2 | 1 | 0 | 0 | 0.276798 |
| Down | SPRR3 | 1 | 0 | 0 | 0.264717 |

**Table 5** miRNA - target gene and TF - target gene interaction

| **Regulation** | **Target Genes** | **Degree** | **MicroRNA** | **Regulation** | **Target Genes** | **Degree** | **TF** |
| --- | --- | --- | --- | --- | --- | --- | --- |
| Up | IRF4 | 140 | hsa-mir-4484 | Up | ESR1 | 22 | STAT3 |
| Up | ESR1 | 98 | hsa-mir-3668 | Up | IRF7 | 14 | KLF5 |
| Up | EIF2AK2 | 98 | hsa-mir-190b | Up | IRF8 | 12 | MAX |
| Up | STAT1 | 63 | hsa-mir-4693-5p | Up | EIF2AK2 | 10 | POU2F2 |
| Up | FYN | 61 | hsa-mir-4326 | Up | FYN | 10 | ARID3A |
| Up | UBD | 34 | hsa-mir-4287 | Up | IRF4 | 10 | JUN |
| Up | FBXO6 | 30 | hsa-mir-5095 | Up | ISG15 | 10 | NR2E3 |
| Up | ISG15 | 29 | hsa-mir-206 | Up | STAT1 | 9 | GATA2 |
| Up | TGM2 | 27 | hsa-mir-1285-3p | Up | TGM2 | 7 | USF2 |
| Up | PML | 26 | hsa-mir-378a-3p | Up | FBXO6 | 6 | HNF4A |
| Up | FASLG | 25 | hsa-mir-7703 | Up | UBD | 6 | TEAD1 |
| Up | IRF8 | 18 | hsa-mir-4686 | Up | PTN | 6 | FOXC1 |
| Up | IRF7 | 17 | hsa-mir-520c-3p | Up | FASLG | 5 | NR3C1 |
| Up | PTN | 12 | hsa-mir-449a | Up | PML | 5 | NFIC |
| Up | NOS2 | 6 | hsa-mir-1291 | Up | NOS2 | 4 | NFYA |
| Down | MAPK1 | 286 | hsa-mir-1470 | Down | BCL6 | 17 | SREBF2 |
| Down | EEF2 | 229 | hsa-mir-2110 | Down | CEBPB | 12 | E2F6 |
| Down | MYH9 | 226 | hsa-mir-3615 | Down | EGR1 | 12 | SRF |
| Down | PRKDC | 164 | hsa-mir-4511 | Down | RXRA | 12 | GATA3 |
| Down | PABPC1 | 151 | hsa-mir-1825 | Down | ARRB2 | 12 | USF1 |
| Down | EGR1 | 132 | hsa-mir-500b-5p | Down | FOS | 11 | FOXD1 |
| Down | FOS | 105 | hsa-mir-204-3p | Down | EEF2 | 10 | NRF1 |
| Down | CEBPB | 94 | hsa-mir-573 | Down | PXN | 10 | IRF2 |
| Down | UBE2D1 | 84 | hsa-mir-326 | Down | UBE2D1 | 9 | HOXA5 |
| Down | MAP1LC3B | 68 | hsa-mir-3180-5p | Down | MAPK1 | 9 | TP53 |
| Down | PXN | 60 | hsa-mir-1255a | Down | MAP1LC3B | 8 | ZNF354C |
| Down | NFKBIA | 54 | hsa-mir-548f-3p | Down | NFKBIA | 8 | TFAP2C |
| Down | ARRB2 | 54 | hsa-mir-4298 | Down | PABPC1 | 7 | SOX5 |
| Down | BCL6 | 43 | hsa-mir-409-5p | Down | MYH9 | 5 | CREB1 |
| Down | RXRA | 41 | hsa-mir-6782-3p | Down | PRKDC | 3 | E2F1 |
